# Anaerobic protist survival in microcosms is dependent on microbiome metabolic function

**DOI:** 10.1101/2025.03.26.644077

**Authors:** Karla Iveth Aguilera-Campos, Julie Boisard, Viktor Törnblom, Jon Jerlström-Hultqvist, Ada Behncké-Serra, Elena A Cotillas, Courtney W Stairs

**Affiliations:** Department of Biology, Lund University, Lund, Sweden; Department of Cell and Molecular Biology, Uppsala University, Uppsala, Sweden

## Abstract

Anaerobic environments serve as habitats for diverse microorganisms, including unicellular eukaryotes (protists) and prokaryotes. To thrive in low-oxygen environments, protists and prokaryotes often establish specialized metabolic cross-feeding associations, such as syntrophy, with other microorganisms. Previous studies show that the breviate protist *Lenisia limosa* engages in a mutualistic association with a denitrifying *Arcobacter* bacterium based on hydrogen exchange. Here, we investigate if the ability to form metabolic interactions is conserved in other breviates by studying five diverse breviate microcosms and their associated bacteria We show that five laboratory microcosms of marine breviates live with multiple hydrogen-consuming prokaryotes that are predicted to have different preferences for terminal electron acceptors using genome-resolved metagenomics. Growth of the prokaryotes and protists within the microcosms respond differently to electron acceptors depending on the make-up of the prokaryotic community. We find that the metabolic capabilities of the bacteria and not their taxonomic affiliations determine protist growth and survival and present new potential protist-interacting bacteria from the *Arcobacteraceae*, *Desulfovibrioaceae* and *Terasakiella* lineages. This investigation uncovers potential nitrogen and sulfur cycling pathways within these bacterial populations, hinting at their roles in syntrophic interactions with the protists via hydrogen exchange.

## INTRODUCTION

Anaerobic environments serve as habitats for a myriad of microorganisms, including unicellular eukaryotes (protists) and prokaryotes. To thrive in low-oxygen environments, protists and prokaryotes often establish specialized metabolic cross-feeding associations such as syntrophy with other microorganisms[1–3]. Some syntrophic interactions in low-oxygen environments are key to biogeochemical functions[3, 4], agriculture[5], and bioelectricity generation[6]. For example, nearly all biological methane emission on our planet derives from syntrophic interactions between methanogenic archaea and hydrogen-producing bacteria[7]. From an evolutionary standpoint, establishing syntrophic relationships allow collaborating organisms to persist in a new environment that could not be accessed individually and can lead to the evolution of new lineages shaped by co-evolution of each partner[3, 8].

Syntrophic interactions between protists and bacteria have been described across the eukaryotic tree of life within the metamonads[9], diatoms[10], discobids[11], ciliates[12–14], foraminifera[15], amoebozoans[16], and breviates[17]. In these cases, the anaerobic eukaryote hosts prokaryotic endo- or ectosymbionts that likely utilize hydrogen produced by their hosts to fuel energy conservation via methanogenesis, nitrate, or sulfate reduction yielding methane, nitrate, ammonium or hydrogen sulfide.

One of these examples includes the breviate protist *Lenisia limosa*[17]. Breviates are free-living amoeboflagellate protists that have only been identified in hypoxic or anoxic environments. They are part of the eukaryotic supergroup ‘Obazoa’ where they branch sister to the Opisthokonts (animals and fungi) and Apusomonads[18]. Breviates possess specialized anaerobic mitochondria, so-called mitochondrion-related organelles (MRO), that likely couple ATP synthesis to hydrogen production rather than oxidative phosphorylation[17, 19, 20]. Attempts to establish axenic cultures have been unsuccessful suggesting that the protist’s growth might rely on interactions with prokaryotes. Indimngseed, a previous study proposed that the breviate *Lenisia limosa* engages in a mutualistic facultative association with an epibiont bacterial partner, *Arcobacter* sp. EP1[17]. In this system, the protist likely produces hydrogen which is used by the epibiont to fuel denitrification in the presence of nitrous oxide or nitrate. Critically, Hamman *et al.* proposed that *L. limosa,* like other protists and some bacteria, can perform electron confurcation whereby a low potential (*e.g.,* ferredoxin) and higher potential (*e.g.,* NAD(P)H) electron donor are used *in concert* to fuel hydrogen production by a specialized [FeFe]-hydrogenase. Such a reaction requires a low partial pressure of hydrogen which can be facilitated by a hydrogen-scavenging epibiont[21].

Here, we examine the diversity of breviate-associated bacteria (including *Arcobacteraceae*) and cross-feeding potential of five diverse breviate-containing microcosms. We tested the hypothesis that different electron acceptors would influence the growth of the protists depending on the metabolic capabilities of the prokaryotic community[3, 22, 23]. Using metagenomics, fluorescent *in situ* hybridization and amplicon sequencing analysis, we identify prokaryotes that might be important for the protist’s survival under different conditions. We find that one breviate species can only grow in the presence of sulfate while another species favours growth with nitrate over sulfate, suggesting that differences in the nature of the prokaryotic community (e.g., sulfate reducers or nitrate reducers) can impact the ability of the protist to survive in anoxia.

## MATERIALS AND METHODS

### Microcosm maintenance and identification

Breviate microcosms PCE, FB10N2, LRM1b, LRM2N6, and *Pygsuia biforma* were provided by Yana Eglit (University of Victoria) and Alastair Simpson (Dalhousie University). All breviate microcosms were collected from marine anoxic environments (Supplementary Datafile S1). Microaerophilic breviates cultures were maintained at 20°C in LBSW (3% lysogenic broth prepared in instant ocean, Aquarium Systems 216034), in 15 ml plastic tubes with minimal headspace and passed weekly. Microaerophilic cultures were maintained in plastic containers with minimal headspace while anoxic cultures were maintained in glass serum flasks where headspace was replaced by argon. The DIC images of the breviates were captured using a Zeiss Axio Observer Z1 motorized inverted fluorescence microscope and processed with Fiji. DNA isolation for metagenomic sequence and amplicon sequencing are described in the Supplementary Material. The eukaryotic 18S was amplified using EukA and EukB primers[24, 25].

### 18S environmental sequencing survey and phylogenetics

The breviate 18S sequences from the microcosms (PCE, LRM1b, FB10N2, LRM2N6) and previously reported sequences (*Pygsuia biforma,* KC433554.1; *Breviata anathema,* AF153206.1; *Subulatomonas tetraspora,* HQ342676.1; *Lenisia limosa,* KT023596.1) were used as a query against the Integrated Microbial Next Generation Sequencing (IMNGS) platform[26] longer than 200 bp and >97% (Supplementary Datafile S1). The *Lenisia limosa* query failed to retrieve any sequences from the IMNGS database. To validate the retrieved sequences, we performed an additional taxonomic assignment of all the putative ‘breviate’ sequences using SILVA SINA search-and-classify[27] using a 97% cut-off and removed all non-Obazoan, Fungi, and Opisthokonta sequences. The putative breviate environmental sequences were added to a dataset of annotated breviate sequences available on Genbank and metaPR2[28] together with a selection of apusomonad sequences as an outgroup. Sequences were aligned using SSU-align and masked using SSU-mask[29] using default settings. Phylogenies were inferred using IQTREE v2.0[30, 31] under the best scoring model of evolution decided by ModelFinder and 500 non-parametric bootstraps (-b 500) (Supplementary Datafile S2).

### Amplicon sequencing

To explore the changes for the prokaryotic community in response to nitrate, we collected DNA from anoxic and microaerophilic microcosms grown in LBSW or LBSW- NIT (supplemented with 2 mM KNO_3_) for 7 d. The V4 region of the 16S rRNA gene was amplified from each biological replicate in three technical replicates using the barcoded primers 515F[32] and 806R[33] (Supplementary Material). Library preparation and sequencing with MiSeq paired-end were performed by Eurofins with their NGSelect Amplicon 2nd PCR service. Adapter sequence removal and read merging were performed by Eurofins using Cutadapt v2.7[34] and FLASH v2.2.00[35], respectively. The resulting data was processed using qiime2 v2023.9[36] (Supplementary Material). We calculated the relative abundance[36], visualized the data in R and calculated differential abundance across conditions with ANCOM[37].

### Breviates growth curves with different electron acceptors in anoxia

To test which electron acceptors impact breviate growth, we grew anoxic microcosms in sulfate-free and nitrate-free 2 mM HEPES-buffered (pH 7.0) defined seawater supplemented with 3% LB (dSW): dSW (no added electron acceptors), dSW-NIT (supplemented with 2 mM KNO_3_) and dSW-SULF (supplemented with 10 mM MgSO_4_) each supplemented with prey bacteria (*Klebsiella pneumoniae,* 1 x 10^9^ cells/ml). Klebsiella pneumoniae was a gift from Dr. Elisabeth Gauger, Lund University. dSW was prepared with 27.72 g of NaCl, 0.67 g of KCl, 1.36 g of CaCl_2_.2H2O, 9.32 g MgCl_2_.6H_2_O and NaHCO_3_ to 1 L, modified from[9] and supplemented with 30 mL of LB. Three biological replicates were prepared by inoculating 5 ml of *P. biforma* or LRM1b breviate cultures in 55 ml of dSW, dSW-NIT or dSW-SULF in 100 mL serum flasks.

To generate the growth curves, 1 ml of each culture was collected anaerobically with a needle and a syringe at 0, 3, 5, 7, and 10 d and analyzed immediately using a Benchtop B3 series FlowCAM (Yokogawa Fluid Imaging Technologies Inc.), a fluid particle imaging system designed to capture and analyze particles in liquid samples. For each sample, the FlowCAM captured 1000 particles in the 3–40 μm diameter range, with 3 technical replicates generated per sample. All images are available at figshare.com/s/6a5781edc1a4e0e722bc. To distinguish images containing a protist-like object from non-protist material, we developed a Gradient Boosting Classifier (https://github.com/theLabUpstairs/FlowCam_Image_Classifier) by training on the image metadata variables with the most relevant features selected using Recursive Feature Elimination based on a Random Forest Classifier. The protist concentration in each sample was estimated by dividing the number of identified protist images by the volume of fluid imaged (Supplementary Datafile S4) and statistical analyses were performed using a two-way ANOVA (Supplementary Datafile S5).

### Metagenomic sequencing, assembly and binning

DNA samples of PCE, FB10N2, LRM1b and LRM2N6 were sequenced by Eurofins (Standard genomic library, NovaSeq 6000 S4 PE150 XP). Each strain resulted in 1 paired-end libraries except for strain FB10N2 which resulted in 2 paired-end libraries that were concatenated (R1, R2) (Supplementary Datafile S6). Read quality was assessed using fastqc 0.11.9[38] and reads were trimmed using Trimmomatic 0.39[39]. All strains were assembled independently using Spades 3.15.2 (options: – meta)[40]. Long-read metagenomic sequencing of microcosm DNA was performed by the National Genomics Infrastructure Sweden (PCE, LRM1b, LRM1N6, FB10N2; kit: SQK-LSK109) or in-house (*Pygsuia biforma,* kit: SQK-NBD114.24) with ONT ligation library prep with barcoding on one ONT PromethION flowcell (NBIS: FLO-PRO002 and in-house: FLO-PRO114M). Details on base-calling and adaptor trimming can be found in Supplementary material. DNA from each microcosm was assembled independently using Flye 2.9.1 (--meta)[41]. Contig clustering, manual binning and gene calls were performed using anvi’o 8[42].

Metagenome-assembled genomes (MAGs) were reassembled into high-quality genomes for *Arcobacteraceae*, *Desulfovibrionaceae* and *Terasakiella* by mapping back the reads from each sample to each MAG using bowtie2 2.4.2 (short reads)[43], minimap2 2.24-r1122 (long reads) and samtools 1.14 (both). Reads were processed through iterative assemblies and contig clustering using Trycycler v0.5.4 to generate consensus long-read assemblies[44]. Read correction was performed using medaka v1.7.2 (https://github.com/nanoporetech/medaka) (long reads for PCE, FB10N2, LRM1b, LRM2N6) or medaka v2 (*Pygsuia biforma*) and polypolish 0.5.0 (short reads for PCE, FB10N2, LRM1b and LRM2N6)[45].

### Taxonomic assignment, phylogenomic tree construction and annotation of the assembled genomes

Taxonomic assignment of *Arcobacteraceae, Desulfovibrionaceae* and *Terasakiella* genomes was performed using GTDBtk 2.4.0[46]. All genomes were placed in GTDB phylogenomic tree with pplacer 1.1[46, 47] (--classify_wf) to estimate the taxonomic placement. Taxonomic inference was refined using the --denovo_wf with defined outgroups (doi.org/10.17044/scilifelab.28254575) and visualized with iTOL 7[48]. Gene predictions were performed with prodigal through anvi’o 8[49]; metabolic annotations were performed by searching the KEGG database[50] with hmmsearch[51] through anvi’o 8. Genome statistics are summarized in Supplementary Datafile S6.

### Fluorescence in situ hybridization

Breviates were grown on a slide and incubated overnight in a moist chamber in an anaerobic jar under anaerobic conditions with Anaerocult A (Millipore 113829, Sparks, Maryland, USA) and subsequently fixed in 4% formaldehyde (ThermoScientific 28906, Rockford, IL, USA) for 15 min. Slides were rinsed with water, immersed in ethanol and incubated with hybridization buffer (Suppementary Material) containing 5 ng/μl of FISH probes targeting *Arcobacteraceae*[52], deltaproteobacteria[9] and *Terasakiella* (Supplementary Datafile S2). After incubation, slides were washed and stained with DAPI. Slides were mounted with SlowFade Diamond Antifade Mountant (ThermoScientific P36970, Eugene, Oregon, USA) and imaged on a Zeiss Axio Imager.z2 microscope, using a Plan-Apochromat 100×/1.40 Oil Ph 3 M27 immersion oil objective lens. The following filters were used: set 38 for Atto 488 (excitation, BP 470/40 nm; emission, BP 525/50 nm), set 31 for Atto 550 (excitation, BP 565/30 nm; emission, BP 620/60 nm), set 50 for Atto 633 (excitation, BP 640/30 nm; emission, BP 690/50 nm) and set 49 for DAPI (excitation, G 365 nm; emission, BP 445/50 nm). Images were processed using linear adjustments (e.g., brightness/contrast) in Fiji.

## RESULTS & DISCUSSION

### Breviate protists are diverse and found in low-oxygen environments

Here, we focus on five laboratory-maintained microcosms containing breviate protists (PCE, FB10N2, LRM1b, LRM2N6, and *Pygsuia biforma*) and prokaryotes isolated from brackish environments. The breviate cells show the characteristic pear-like cell shape with two flagella (usually only one is visible) emerging from the cell (Fig. 1A). All species were observed to have pronounced and sometimes branched filopodia as previously described for other breviates[17, 18, 20, 53]. We amplified and sequenced the 18S rRNA gene and used these as a query to retrieve environmental sequences >97% identical available on the IMNGS database[26]. We recovered 78 putative environmental breviate sequences that derive from 68 projects targeting low-oxygen marine intertidal sediments, deep marine sediments, freshwater sediments, compost, a decommissioned mine and soil (Fig. 1B, Supplementary Datafile S1). We reconstructed a phylogenetic tree incorporating the microcosm, environmental and breviate annotated MetaPR2 sequences and found that the microcosm breviate sequences resolved into three distinct groups most closely related to *Subulatomonas tetraspora* (FB10N2), *Pygsuia biforma* (LRM1b) and a clade of environmental sequences derived from marine sediments (PCE, LRM2N6; Fig. 1C).

**Figure 1.**
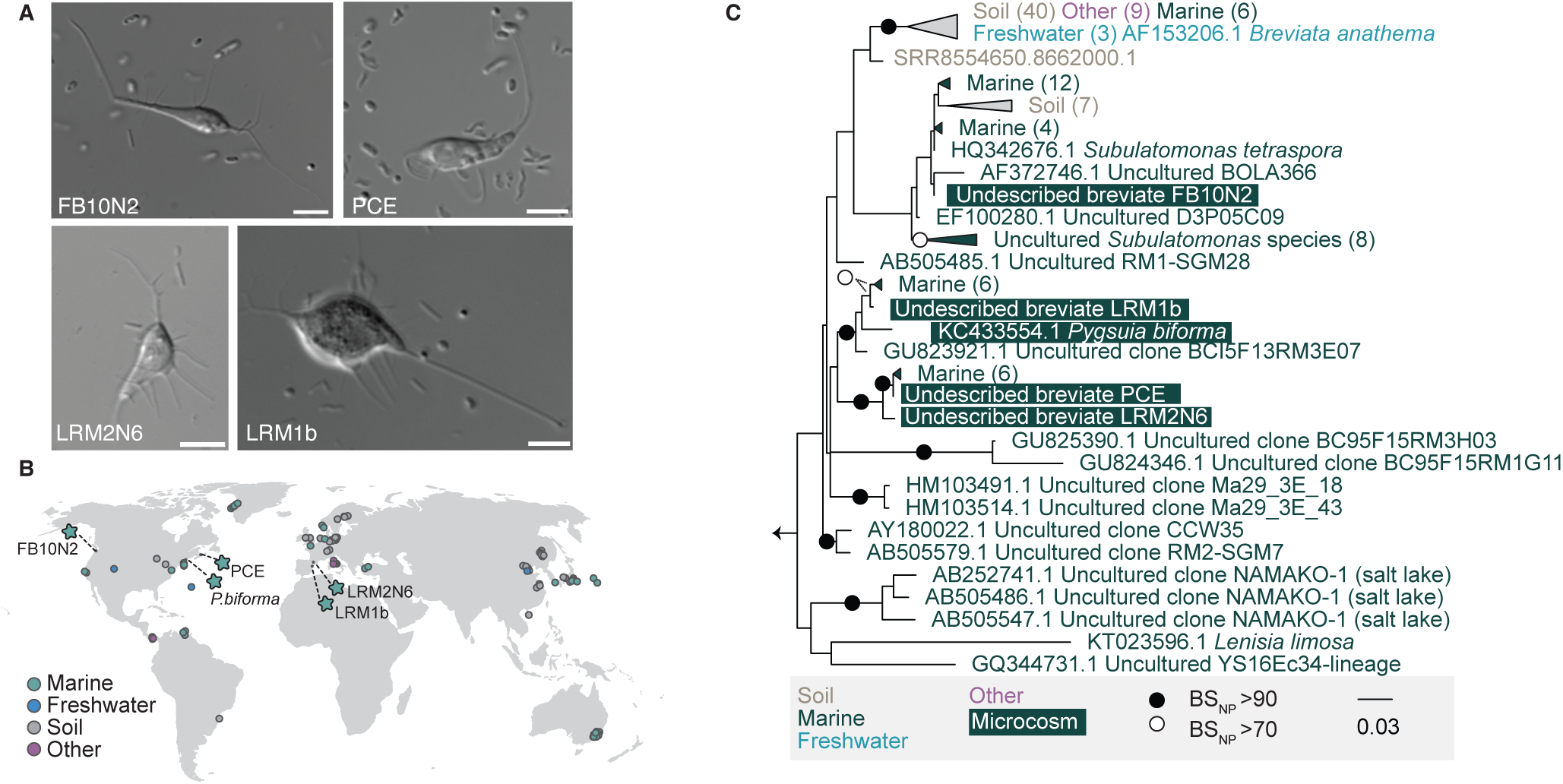
Small subunit ribosomal RNA phylogenetic analysis reveals that cultured breviates resolve into three clades within Breviatea. **a.** Unrooted maximum-likelihood phylogenetic analysis was computed on an alignment of 156 sequences with 1542 sites. Non-parametric bootstrap support greater than 70% or 90% are mapped onto each bipartition with an open or closed circle, respectively. Breviate sequences from microcosms are shaded in dark green. All other sequences derive from environmental sequences (coloured based on sampling location) from publicly available data. **b.** Differential interference contrast images of the four undescribed breviate species in this study. Scale bar 5 µm. **c.** Geographical locations where one or more breviate 18S sequence was identified in circles, breviate strain isolation sites in stars, and nature of the environment are indicated with colours.

### Breviate growth and relative abundance of bacteria varies with availability of electron acceptors

*Lenisia limosa* was shown to display higher growth rates in nitrate and nitrous oxide in comparison to when no electron acceptors were added[17]. To explore if there was a correlation of the bacterial composition, protist growth, and electron acceptors, we evaluated the cell density of *P. biforma* and LRM1b in anoxic conditions when supplemented with nitrate, sulfate or no added electron acceptor (noE) over time (Fig 2A,B). These protists were selected because they were closely related but had distinct microbial communities (Fig 2C, discussed below) allowing us to test if community composition can influence protist dynamics. Both protists were able to grow in media with no added electron acceptors (containing only those electron acceptors present in the LB and cell inoculum). *P. biforma* grew significantly better in media with sulfate in comparison with noE (Fig. 2B, Supplementary Data File S5), reaching its peak at day 3. *P. biforma* was unable to grow in media with nitrate suggesting that *P. biforma*’s growth is favoured in the presence of active sulfate reducers and inhibited in the presence of active nitrate reducers. LRM1b grew better in media with nitrate in comparison with noE reaching its peak at day 5 (Fig. 2C, Supplementary Data File S5). The growth of LRM1b in media with sulfate or noE was similar, and reached its peak at day 3. This suggests that growth of the LRM1b breviate is likely unaffected by the activity of sulfate reducers. However, unlike *P. biforma*, the LRM1b breviate can grow in the presence of nitrate with distinct growth dynamics compared to sulfate-reducing conditions.

**Figure 2:**
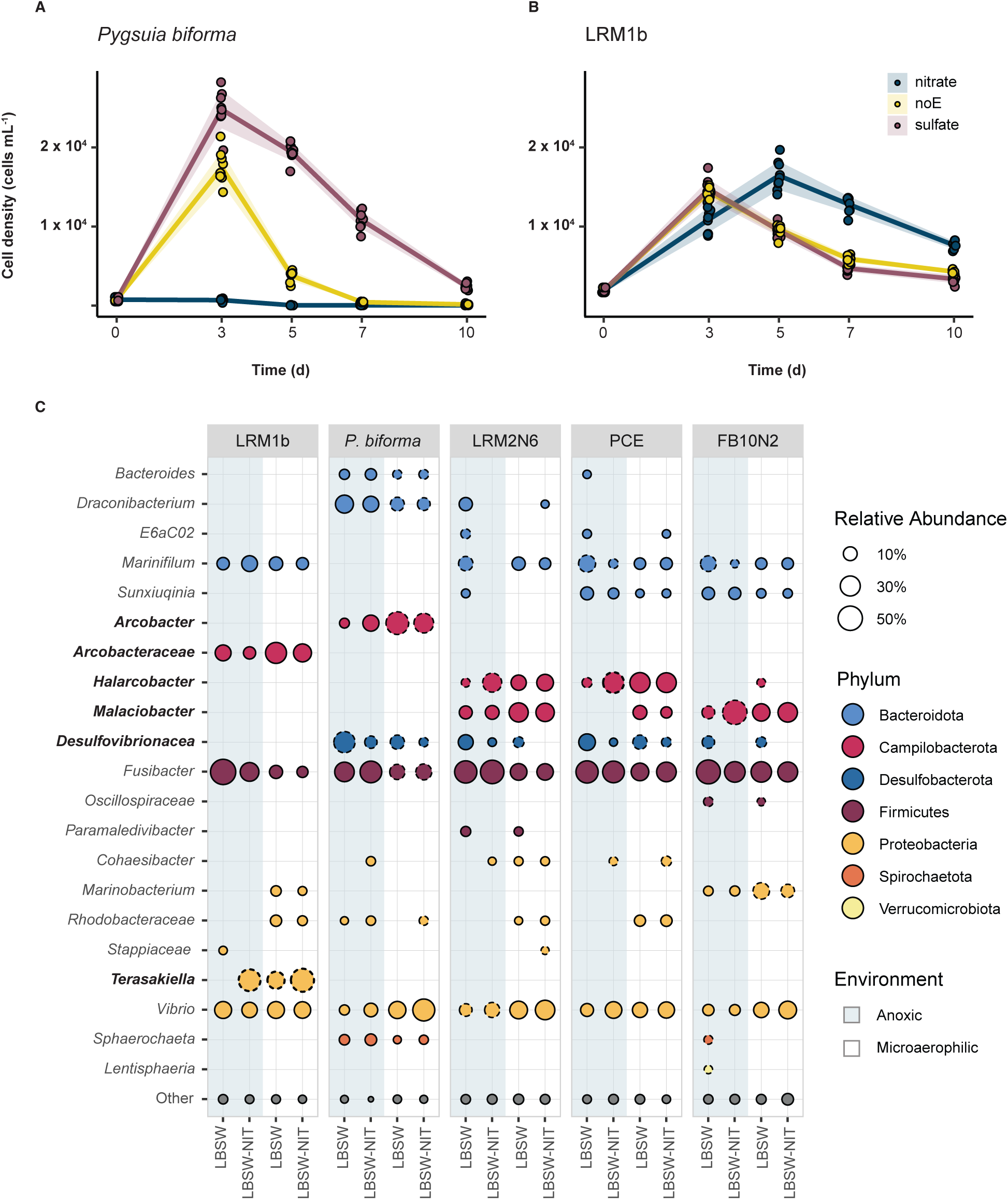
The relative abundance of the breviate microcosms varies with different electron acceptors and correlates with breviate growth. **a.** *P. biforma* and **b.** LRM1b growth curves. Breviate cultures were grown in triplicates in defined seawater (dSW), dSW- NIT or dSW-SULF in anoxia. Protist concentration per ml was calculated at 0, 3, 5, 7 and 10 d. Each datapoint represents a measurement and standard deviations shown with shading. Growth rates of breviates in each microcosm cultured with different electron acceptors were compared using ANOVA and post-hoc test, the growth rates of *P. biforma* were statistically different when grown with the different electron acceptors (p-value < 0.001), while the growth rates or LRM1b were only statistically different between dSW-NIT and dSW-SULF (p-value < 0.05), suggesting a similar growth rate between the conditions. **c.** Bubble plot showing percentage of relative abundance (> 1%) of bacteria present in each indicated microcosm (LRM1b, *P. biforma*, LRM2N6, PCE, FB10N2 columns), growing in anoxic (blue shadow), or microaerophilic conditions (no shadow) in LBSW or LBSW-NIT as indicated. Taxonomic assignment was made at the genus level when possible and coloured based on their phylum designation. Bubbles with dashed outlines indicate taxa with statistically significant differences in relative abundance (ANCOM) between cultures within the same oxic regime with or without nitrate.

To understand the bacterial community associated with the breviates, we performed 16S rRNA gene amplicon analysis under microaerophilic and anoxic conditions supplemented with nitrate or sulfate as electron acceptors (Fig. 2A) and assessed alpha diversity, evenness and beta diversity of the communities (Supplementary Discussion, Supplementary Fig. S1). *Arcobacteraceae* species were among the most abundant species in all the breviate cultures alongside with *Vibrio* and *Fusibacter*, regardless of the presence of oxygen (Fig. 2A). In nitrate-supplemented anoxic breviate microcosms (with the exception of *P. biforma*), at least one amplicon sequence variant (ASV) displayed a statistically significantly increase in relative abundance when compared to the anoxic cultures without nitrate (Fig. 2A, Supplementary Fig. S1). Most of these organisms belong to the *Arcobacteraceae* including *Malaciobacter* (FB10N2) and *Halarcobacter* (LRM2N6 and PCE) (Fig. 2A; Supplementary Data File S3). In the LRM1b microcosm, we detected a higher relative abundance of *Terasakiella*, and not *Arcobacteraceae,* in the presence of nitrate. In *P. biforma,* LRM2N6, PCE, and FB10N2 microcosms, the relative abundance of the taxa classified as *Desulfovibrionaceae* was higher in cultures supplemented with sulfate in comparison to the cultures supplemented with nitrate in both microaerophilic and anoxic microcosms (Fig. 2C; Supplementary Data File S3). *Desulfovibrionaceae* sequences were detected in LRM1b microcosm but below the threshold for inclusion (1% relative abundance). Predictably, we observed that the addition of nitrate or sulfate under anoxia lead to changes in the relative abundance of bacteria typically associated with nitrate reduction (*e.g., Arcobacteraceae* or *Terasakiella*) or sulfate reduction (e.g., *Desulfovibrionaceae*), respectively.

### Genome-resolved metagenomics predict Arcobacteraceae, Desulfovibrionaceae and Terasakiella species are nitrate or sulfate respiring

To understand the metabolic potential of each microcosm, we used metagenomic sequencing to reconstruct genomes of their prokaryotic communities (Fig. 3). For some of the microcosms, we isolated bacteria into pure cultures and performed whole genome sequencing. From these data, we reconstructed high-quality, often circular, genomes from nine *Arcobacteraceae* (Fig. 3A), six *Desulfovibrionaceae* (Fig. 3B), and one *Terasakiella* sp. (Fig. 3C) (Figure 3D, Supplementary Datafile S6). Nine new species names and one new genus were deposited on SeqCode[54] (**<u>seqco.de/r:dhfmnria</u>**) and classified based on the Microbial Genome Atlas guidelines (Supplementary Datafile S6)[55, 56]. Below we discuss the metabolic potential of the community focusing on the *Arcobacteraceae, Desulfovibrionaceae*, and *Terasakiella* members specifically related to hydrogen- and end product-sharing metabolisms.

**Figure 3:**
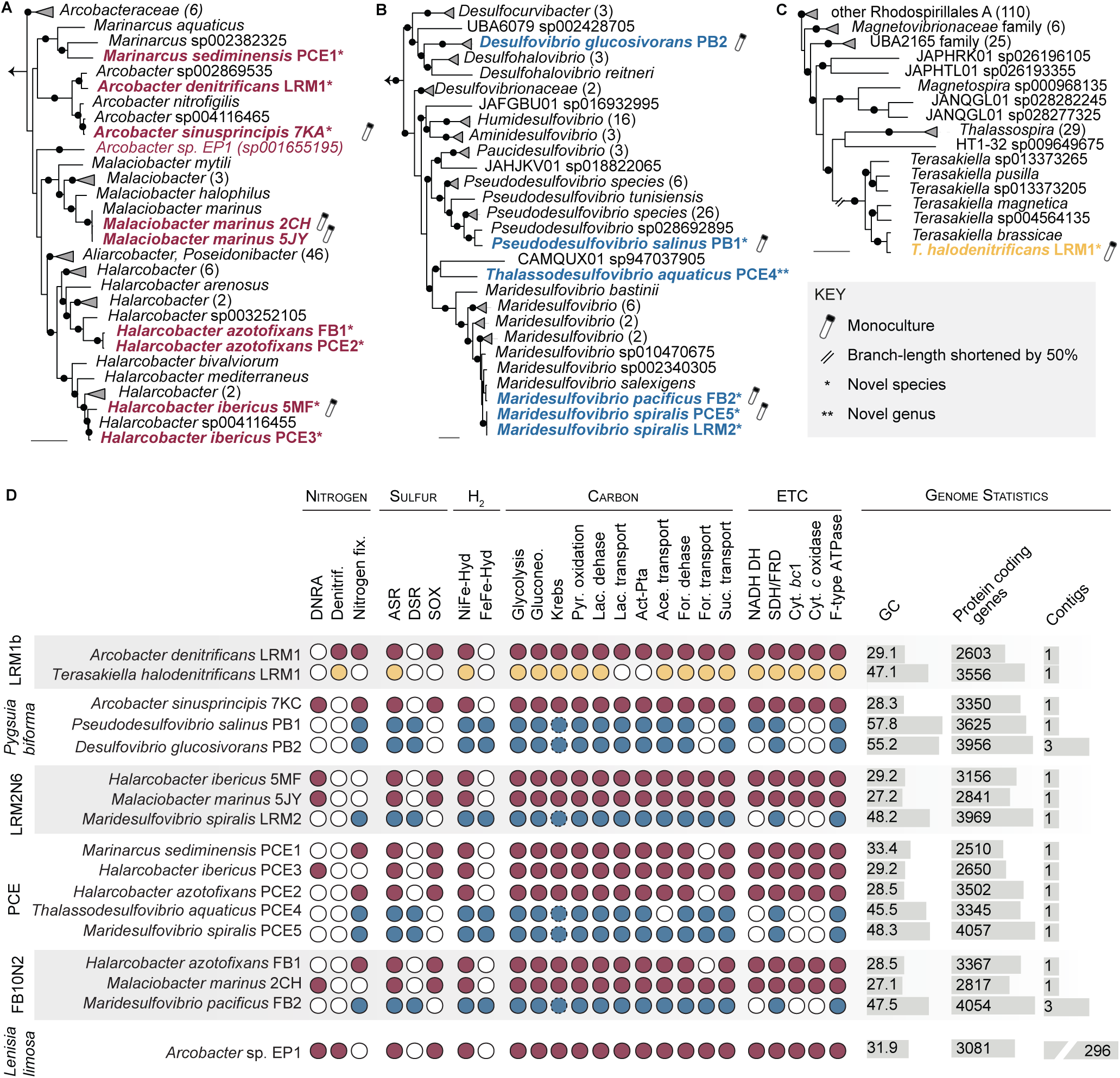
Phylogenetic relationships and metabolic potential of breviate-associated bacteria. GTDB-tk computed phylogenomic trees of breviate-associated bacteria (bold) and their closest relatives (black) across **a.** *Arcobacteraceae*, **b.** *Desulfovibrionaceae*, **c.** *Rhodospirillales* with novel genera (**) or novel species (*) according to [55, 56]. Symbols denote monocultures (test tube icon) and branches shortened by 50% (//). Bootstrap support is indicated at key nodes (filled circles), scale bars represent 0.1 substitutions per site. and 16S phylogenies are shown in Supplementary Datafile S7 **d.** Metabolic potential of breviate-associated bacteria. Presence (coloured circles) or absence (white circles) of genes and pathways for nitrogen, sulfur, and hydrogen metabolism, carbon cycling, and electron transport chains (ETC) are shown for each species. Dotted lines indicate the pathway is incomplete. DNRA, dissimilatory nitrate reduction to ammonium; Denitrif., denitrification; Nitrogen fix., nitrogen fixation; ASR, assimilatory sulfate reduction; DSR, dissimilatory SR; Sox, sulfur oxidation; pyr., pyruvate; lac., lactate; Act-Pta, acetate kinase –phosphate acetyltransferase pathway; ace., acetate; for., formate; cyt., cytochorome; and SDH/FRD, succinate dehydrogenase/fumarate reductase. Genome statistics include GC content, number of predicted protein-coding genes, and number of contigs for each assembly.

#### Breviate-associated *Arcobacteraceae* are facultative anaerobes capable of dissimilatory nitrate reduction or denitrification

We recovered at least one *Arcobacteraceae* MAG in each microcosm (Fig. 3A and Supplementary Datafile S7). Formal species descriptions for *Arcobacter sinusprincipis nov. sp.* and *Halarcobacter ibericus nov. sp.* is in preparation (Aguilera-Campos et al.).

Genes encoding hydrogen-uptake [NiFe]-hydrogenases were identified in all *Arcobacteraceae* MAGs (Figure 3D and Supplementary Datafile S6) and are predicted to donate electrons to the quinol pool. *Halarcobacter ibericus* PCE3 and 5MF, *Malaciobacter marinus* 2CH and 5JY, and *Halarcobacter sinusprincipis* 7KC encode the potential to reduce nitrate for ATP production by dissimilatory nitrate reduction to ammonium (DNRA; via NapGH, NapAB, and NrfAH) using electrons ultimately derived from hydrogen similar to other Campylobacterota[57] (Fig. 3D). This is consistent with their increased relative abundance in media supplemented with nitrate (Fig. 2A). Unlike the *Lenisia limosa*-associated *Arcobacter* sp. EP1, most of our breviate-associated *Arcobacteraceae* lack the genetic capacity for denitrification with the exception of *Arcobacter denitrificans* LRM1 (NAP-β, NirS, NorBC and NosZ; Fig. 3D). Some of the *Arcobacteraceae* genomes encode for nitrogen fixation via NifDKH (Fig. 3D). With respect to sulfate and sulfide-related metabolism, each *Arcobacteraceae* MAG encodes for assimilatory sulfate reduction, sulfide-quinone oxidoreductase (Sqr) and a complete sulfur oxidation (SOX) system. Collectively, these pathways suggest that the *Arcobacteraceae* are able to oxidize hydrogen sulfide, sulfite, thiosulfate or elemental sulfur, common compounds of marine coastal environments[58–60].

All *Arcobacteraceae* encode for glycolysis, gluconeogenesis, the Krebs cycle, pyruvate oxidation to acetyl-CoA, acetate import (ActP), acetate oxidation (Ack-Pta), lactate import (LctP) and lactate oxidation (LldEFG, Dld; Fig. 3D). This suggests that the *Arcobacteraceae* can use acetate or lactate as a carbon source. We also detected the potential for formate oxidation (FdhABCD) which can transfer electrons from formate to the quinone pool to fuel DNRA like other *Arcobacter*[61], although not all the strains encode for a formate transporter. The majority of *Arcobacteraceae* genomes encode a TRAP-type or DcuAB system that is predicted to transport succinate, fumarate and similar molecules. Succinate could serve as an electron or carbon source by the bacteria in the microcosms, indeed *M. marinus*, *H. ibericus* and *A. sinusprincipis* isolates can grow on succinate as the sole carbon source (Aguilera-Campos et al. in preparation). All the genes for aerobic respiration were found in the *Arcobacteraceae* genomes, including NADH:quinone oxidoreductase, fumarate reductase, cytochrome bc1 complex, cytochrome c oxidase, cbb3-type and F-type ATPase, in agreement with their microaerophilic capabilities.

#### Breviate-associated *Desulfovibrionaceae* have the capacity for dissimilatory sulfate reduction

We detected *Desulfovibrionaceae* species in the *P. biforma*, FB10N2, PCE and LRM2N6 microcosm. Some of these genomes likely derive from new species within the *Maridesulfovibrio* and *Pseudodesulfovibrio* genera and at least one likely derives from a new genus *Thalassodesulfovibrio aquaticus* (Fig. 3B). All *Desulfovibrionaceae* genomes encode for hydrogen-uptake [NiFe]-hydrogenases and formate dehydrogenases that are predicted to transfer electrons to a type-I cytochrome *c*_3_[62, 63]. These electrons likely feed into the dissimilatory sulfate reduction (DSR) pathway via the quinone reductase (QrcABCD) and modifying (QmoABC) complexes, adenylylsulfate reductase (AprAB), and dissimilatory sulfate reductase (DsrABCD, DsrMKJOP; Fig. 3D and Supplementary Datafile S6)[62, 64, 65]. Combined with their increased relative abundance in microcosms supplemented with sulfate (Fig. 2A), we suspect all the breviate-associated *Desulfovibrionaceae* are sulfate reducing bacteria (SRB). We also detected the potential for nitrogen fixation in all the *Desulfovibrionaceae* genomes, like other SRB from marine sediments[66].

With respect to carbon metabolism, all of the *Desulfovibrionaceae* genomes encode genes for glycolysis, gluconeogenesis, pyruvate oxidation, Ack-Pta, acetyl-CoA synthetase (ACS) pathways and an F-type ATPase. They have a truncated Krebs cycle (lacking malate dehydrogenase, aconitate hydratase, and succinyl-CoA synthetase), and lack aerobic respiration similar to other *Desulfovibrionaceae*[67, 68]. All breviate-associated *Desulfovibrionaceae* genomes encode for lactate import (LutP) and oxidation (LDH). This suggests that the *Desulfovibrionaceae* could accept electrons from either the hydrogen-uptake machinery above or lactate to fuel DSR. All *Desulfovibrionaceae* genomes encode a TRAP-type transport system small permease protein for succinate/fumarate transport and a fumarate reductase (FrdABC) which can contribute to the quinol pool.

#### Breviate-associated Terasakiella halodenitrificans is a facultative anaerobe with the potential for complete denitrification

We only recovered a *Terasakiella* MAG (complemented with an isolated genome) in the LRM1b microcosm. This likely represents a new species within the genus, *Terasakiella halodenitrificans* (Fig. 3C, Supplementary Datafile S7). The *Terasakiella halodenitrificans* genome encodes a hydrogen-uptake [NiFe]-hydrogenase that is predicted to pass electrons via the quinol pool to the cytochrome *bc*1 complex ultimately yielding reduced cytochrome *c*[69]. We also detected genes for Nap-mediated complete denitrification (via NapAB, NapCDH, NirS, NorBC and NosZ; Fig. 3D, Supplementary Datafile S6) and observed an increase in the relative abundance of the *Terasakiella halodenitrificans* ASV in nitrate-supplemented microcosms (Fig. 2C). *Terasakiella halodenitrificans* encodes all the genes for glycolysis, gluconeogenesis and the Krebs cycle, formate transport, lactate dehydrogenase, and formate dehydrogenase, however, the electron carriers for these systems are unknown. *Terasakiella halodenitrificans* can likely import and metabolize acetate via an ActP and ACS, respectively. We also detected a succinate TRAP-transporter and a succinate dehydrogenase which likely funnel electrons to the quinone pool. Collectively, this suggests that *Terasakiella halodenitrificans* is a facultative anaerobe that can use succinate and acetate as carbon sources.

### Arcobacteraceae, Desulfovibrionaceae and Terasakiella bacteria are associated with breviate cells

To visualize if bacteria were directly interacting with the breviates, we performed fluorescence *in situ* hybridization (FISH) using probes directed against the 16S rRNA of *Arcobacteraceae*, *Desulfobacterota* (formerly ‘deltaprotebacteria’), and *Terasakiella halodenitrificans*. We detected the signal of *Arcobacteraceae* (Fig. 4A, 4C, Supplementary Fig. S2) and *Desulfobacterota* (Fig. 4B, 4D, Supplementary Fig. S3) probes in all the breviate cultures, and the signal of the *Terasakiella* probe in the LRM1b culture (Fig. 4E). While the signal of the bacterial probes was often associated with the protist cells (Fig. 4A, 4B, 4D), we observed signal in non-protist associated bacteria (Fig. 4C). This suggests the interactions between protists and bacteria might be transient. For those microcosms that contained multiple *Desulfobacterota* species, we designed probes to target specific genera of *Pseudodesulfovibrio* (PD1, Supplementary Fig. S4) and *Maridesulfovibrio* (MD1, MD2, Supplementary Fig. S3). *Pseudodesulfovibrio* was detected in all the breviate cultures except FB10N2 (Supplementary Fig. S4). Using these probes, we observed that both *Maridesulfovibrio* and *Pseudodesulfovibrio* reactive cells form associations with breviate cells. It is difficult to know if the potential association of the bacteria with the breviates is due to an interaction or just coincidence. In the *Lenisia limosa* study, there is no discussion of non-protist associated *Arcobacter* and only one cropped micrograph was provided[17].

**Figure 4:**
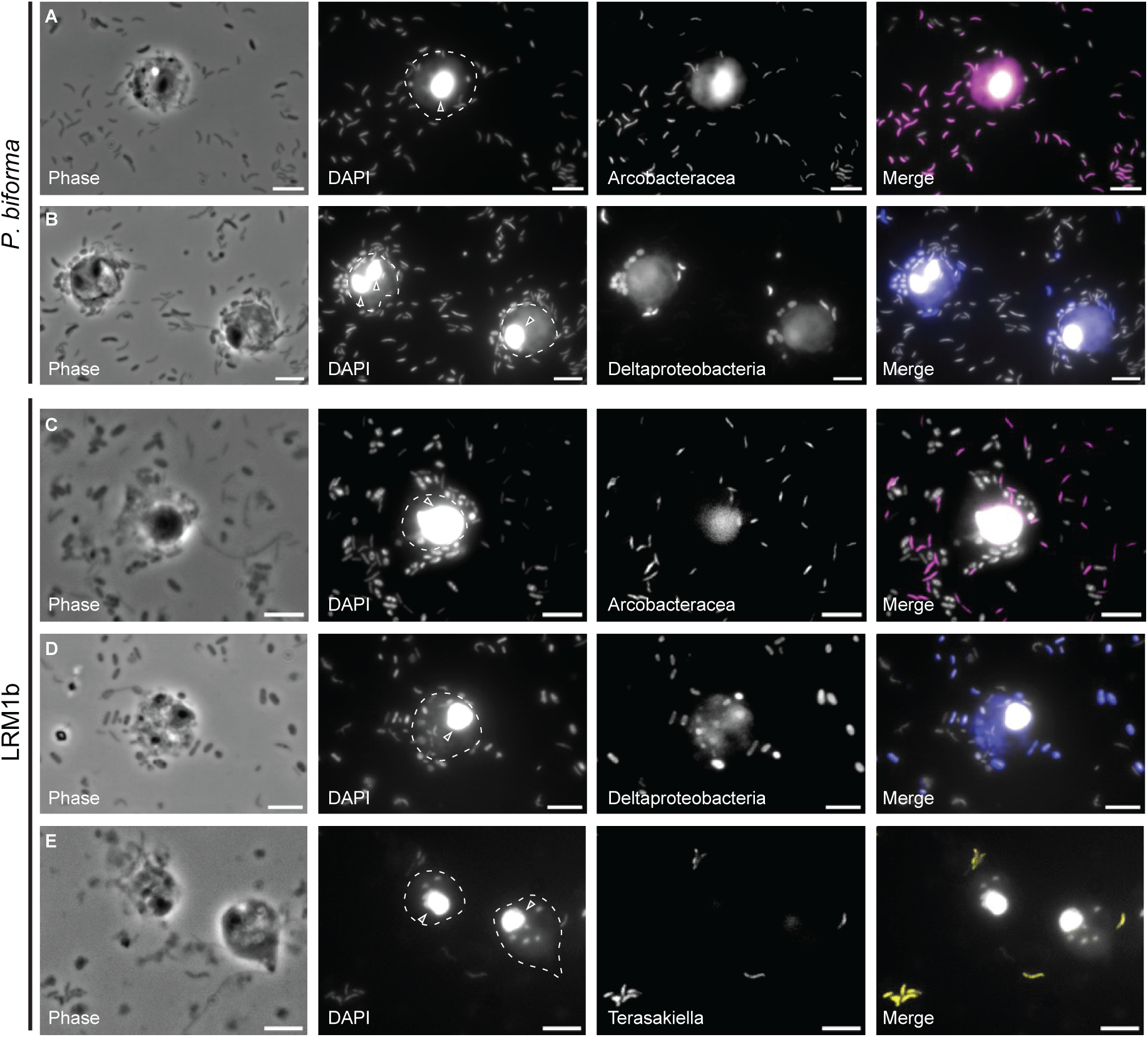
Fluorescence *in situ* hybridization (FISH) suggests a non-obligate direct interaction of *P. biforma* and LRM1b with hydrogen-consuming bacteria. *P. biforma* (**a** and **b**) and LRM1b (**c**, **d** and **e**) cultures were incubated on a slide overnight in an anaerobic glove box. Cultures were fixed (4% formaldehyde) and hybridized with 16S rRNA probes: Arc1430-Atto 488 and Arc94-Atto 488 targeting *Arcobacteracea* cells (magenta, **a** and **c**), Delta495a-Atto 550 probe targeting Desulfobacterota (formerly ‘Deltaproteobacteria’; dark blue, **b** and **d**) and Tera537-Atto 550 probe targeting *Terasakiella halodenitrificans* LRM1 (yellow, **e**). Cell body (outlines) and nuclei (white arrow) of the protists are shown in the DAPI panel. Panels left to right: Phase, DNA stained with DAPI (the big circles correspond to the breviate nucleus), specific FISH probes, and merge channel of DAPI with FISH probes. Scale bar 5 µm

Extracellular relationships between prokaryotes and protists can vary in the nature of the interaction. For example, some protists have sophisticated cellular structures that coordinate their prokaryotic partners[9, 11, 70]. However, some extracellular interactions of *Desulfovibrio* or *Arcobacter* with other protists are sometimes transient[71]. Future experiments should interrogate the contact-dependence of breviate:prokaryote interactions, specifically to examine the function of the chemotaxis and cell:cell adhesion machinery identified herein (Supplementary Material).

### Breviates are predicted to produce hydrogen that can be used by Arcobacteraceae, Desulfovibrionaceae and Terasakiella

Previous studies in *P. biforma* and *L. limosa* predict that breviates produce hydrogen as an end product of metabolism by the action of confurcating NAD(P)H-dependent [FeFe]-hydrogenases which likely require a low-partial pressure of hydrogen to function[17, 19] (Fig. 5A). Therefore, the metabolism of the protist would benefit a hydrogen-scavenging prokaryote, which could therefore serve as a hydrogen sink to keep the partial pressure of hydrogen low as previously proposed in ciliates[72], metamonads[9, 68, 73], amoebozoans[16], and *L. limosa*[17]. Here, we propose that nitrate-reducing, denitrifying or sulfate-reducing bacteria, could serve as the hydrogen sink in the breviate microcosms. For example, the LRM1b-associated *Terasakiella halodenitrificans* and *Arcobacter denitrificans* may couple hydrogen oxidation with denitrification (Fig. 5D). Similarly, the breviate-associated *Arcobacteraceae* (*Halarcobacter ibericus*, *Malaciobacter marinus* and *Halarcobacter sinusprincipis*) might use hydrogen as an electron donor for DNRA (Fig. 5D), as was observed for other *Campylobacterota* isolates[57]. Finally, the *Desulfovibrionaceae* SRB serve as the hydrogen sink via DSR (Fig. 5B) as has been proposed for other symbionts of protists[68]. These microcosm inferences might reflect the ecology of these organisms in their natural environments as sequences from breviates recovered from environments known to host DNRA, denitrifying and sulfate-reducing bacteria (*e.g.,* Marmara sea sediment[74]:HM103514, HM103491; Disko island tidal sediments[75]: EF100226, EF100280, EF100389, EF100391, EF100407, EF100410).

**Figure 5:**
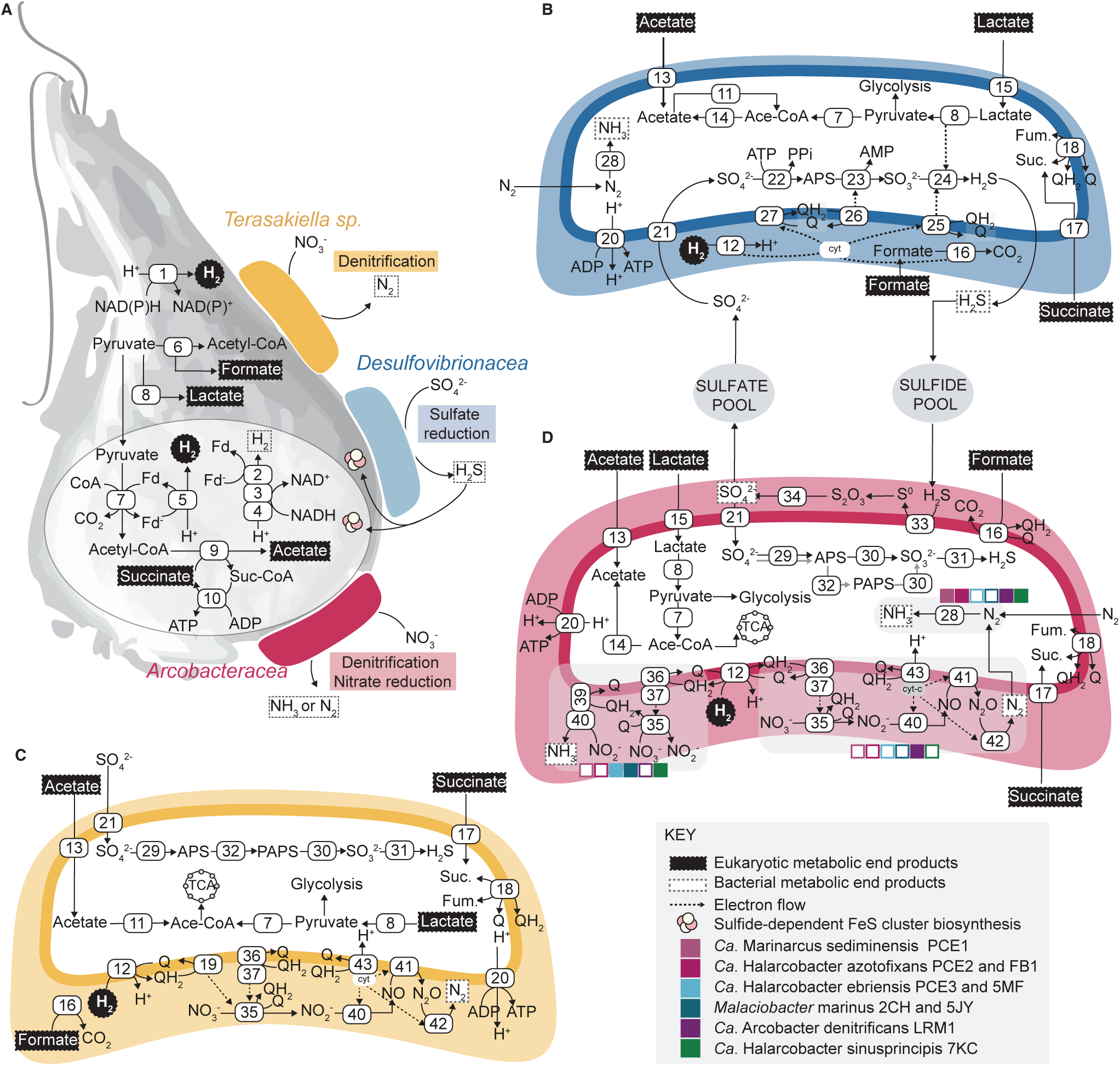
Prediction of potential metabolic cross-feeding of breviate-associated bacteria with breviates. **a** Schematic representation of a potential metabolic interaction of the breviate cell (in grey) with the associated bacteria, *Desulfovibrionaeae* (in blue), *Arcobacteraceae* (in red) and *Terasakiella sp* (in yellow). The breviate cell produces acetate, succinate, lactate, formate and H_2_ as end products of pyruvate and energy metabolism in the cytoplasm and/or in the mitochondrion related organelle (MRO), these products can then be used by the associated bacteria as carbon or energy source, for instance for dissimilatory sulfate reduction, dissimilatory nitrate reduction or denitrification. The metabolism of the breviate cell is predicted from transcriptomes of the *Pygsuia biforma*[19] and *Lenisia limosa*[17] already reported. The metabolism of the breviate-associated bacteria is predicted from our assembled genomes. Predicted metabolism of *Desulfovibrionaceae* (**b**), *Terasakiella sp*. (**c**) and *Arcobacteraceae* (**d**). Within each cell is represented the predicted pathways involved in Nitrogen, Sulfur, Carbon and Hydrogen metabolism. In dashed squared metabolic end products. 1) NAD(P)H-dependent FeFe-hydrogenase, 2) electron bifurcating FeFe- Hydrogenase, 3) NuoE: NADH:ubiquinone oxidoreductase subunit E, 4) NuoF: NADH:ubiquinone oxidoreductase subunit F, 5) ferredoxin-dependent Hydrogenase, 6) PFL: pyruvate formate-lyase, 7) PFO/POR: pyruvate-ferredoxin oxidoreductase, 8) LDH: lactate dehydrogenase, 9) ASCT: Acetate:succinate CoA transferase, 10) SCS: succinyl coenzyme A synthetase, 11) ACS: Acetyl-coenzyme A synthetase, 12) NiFe-hydrogenase, 13) Acetate transporter (e.g. ActP), 14) Ack/Pta: acetate kinase phosphate acetyltransferase pathway, 15) LutP: lactate permease, 16) FDH: formate dehydrogenase, 17) TRAP: TRAP-type transport system small permease protein, 18) FRD: fumarate reductase/SDH succinate dehydrogenase; 19) NapC: cytochrome c-type protein. 20) ATP synthase, 21) SulP: sulfate permease, 22) Sat: sulfate adenylyltransferase, 23) AprAB: adenylylsulfate reductase subunit A and B, 24) DsrABCD: dissimilatory sulfite reductase subunits 25) DsrMKJOP dissimilatory sulfite reductase subunits, 26) qmoABC: quinone-modifying oxidoreductase, 27) QrcABCD: menaquinone reductase, 28) NifDKH: nitrogenase, 29) CysN: sulfate adenylyltransferase subunit 1, 30) CysH: phosphoadenosine phosphosulfate reductase, 31) CysJI: sulfite reductase, 32) CysC: adenylylsulfate kinase, 33) Sqr: sulfide:quinone oxidoreductase, 34) SOX: sulfide oxidation system, 35) NapAB: nitrate reductase, 36) NapH: ferredoxin-type protein, 37) NapG: ferredoxin-type protein, 38) NrfA: nitrite reductase, 39) NrfH: cytochrome *c* nitrite reductase, 40) NirS: nitrite reductase, 41) NorBC: nitric oxide reductase, 42) NosZ: nitrous oxide reductase, 43) Cyt bc1: cytochrome *bc*1 complex, Other abbreviations: TCA: tricarboxylic acid cycle, cyt-c: cytochrome c, Ace-CoA: Acetyl coenzyme A, Suc-CoA: succinyl coenzyme A.

In contrast to *L. limosa*, four of our microcosms lack bacteria capable of complete denitrification and *P. biforma* was unable to grow in the presence of nitrate as the electron acceptor. We suggest that the denitrification potential of the most abundant prokaryote in the LRM1b microcosm, *Terasakiella halodenitrificans* (and less abundant *Arcobacter denitrificans*) explains why LRM1b and not *P. biforma* can grow with nitrate. This suggests that, in general, denitrification is compatible with LRM1b survival or that the byproducts of dissimilatory nitrate reduction (*e.g.,* ammonia) can be toxic to *P. biforma*, and not *L. limosa* or LRM1b microcosms. If nitrate is available, then nitrate-reducing or denitrifying bacteria (*Arcobacteracea*, *Terasakiella*) will use LRM1b microcosm end-products and outcompete the sulfate-reducing bacteria due to more favorable energy potential of DNRA and DN over DSR[76, 77]. If sulfate is available, sulfate-reducing *Desulfovibrionaceae* and sulfide-oxidizing *Arcobacteraceae* species can use microcosm end-products and likely maintain sulfide homeostasis compatible with *P. biforma* growth.

While sulfide can be toxic to eukaryotes[78–80], we suspect that the breviates might rely on sulfide for the biosynthesis of Fe-S clusters. Most eukaryotes synthesize this essential biological cofactor using the Iron-Sulfur Cluster (ISC) system that relies on cysteine-derived sulfur. *P. biforma* lacks the ISC system and instead encodes an archaeal minimal sulfur mobilization system (SMS)[19] that, in archaea, relies on sulfide[81]. Therefore, we suspect that the breviates might be able to directly utilize environmental sulfide for Fe-S cluster metabolisms and have increased resilience in sulfidic environments such as deep sea[74] or tidal [75] sediments. In addition, sulfide homeostasis could be regulated by sulfide-oxidizing *Arcobacteraceae* via their SOX system (Fig. 5D) as seen in marine invertebrate symbioses involving *Arcobacteracaea*[71] and other bacteria[82–84].

### Breviates likely produce formate, acetate, lactate and succinate that can be used by Arcobacteraceae, Desulfovibrionaceae and Terasakiella

To examine if the protists can produce metabolites compatible with the predicted metabolism of the bacteria, we queried the genome and transcriptome data for *P. biforma* and *L. limosa* for genes related to formate, acetate, lactate and succinate production as a proxy for breviate metabolism (Fig. 5A, Supplementary Datafile S6). *P. biforma* and *L. limosa* encode a pyruvate formate lyase and formate transporter (FocA) protein suggesting that the protist might be capable of producing and transporting formate. Both protists are predicted to produce lactate (LDH), acetate (acetate:succinate CoA transferase), and succinate (fumarate-reducing complex II[19, 85] and succinyl CoA synthetase), although specific export transporters could not be confidently identified.

These end-products are compatible with all of the examined bacteria and are consistent with known growth dependencies of related species. For example, most of the breviate-associated *Desulfovibrionaceae* genomes described here encode genes for lactate, formate and succinate/fumarate transport and utilization (Fig. 3D). Previous studies have shown that related species (e.g., *Desulfovibrio glucosivorans*, *D. desulfuricans)*, can grow on media supplemented with lactate, hydrogen/acetate or formate [67, 86]. Therefore, we suspect the breviate-associated *Desulfovibrioaceae* can likely use community-derived lactate, formate and acetate/hydrogen as substrates for growth (Fig. 5B). Similarly, the protist-associated *Arcobacteraceae* and *Teraskiella* can likely uptake lactate, acetate, and succinate (Fig. 3D, 5D) likely using formate or hydrogen as electron donors as previously reported in other *Arcobacteraceae*[17, 60, 87] and *Terasakiella*[88, 89].

### No evidence for co-evolution of the breviate protists and their associated bacteria

The nature of known eukaryote:prokaryote interactions is diverse, ranging from transient associations to obligate symbioses[90], and is influenced by the environment, as well as the metabolic needs of the interacting partners. For example, in some anaerobic ciliates, the metabolic activities of symbiotic methanogenic archaea or sulfate-reducing bacteria boost the ciliate’s metabolic efficiency[91]. Over evolutionary time, symbiotic interactions can lead to co-evolution of both host and symbiont, including the reduction of symbiont genomes[12, 92], mutual dependence[16] and even vertical transmission or replacement of symbionts across speciation events[93, 94]. We failed to detect obvious prokaryotic genome reduction or co-speciation patterns in the five studied breviate microcosms. However, given the diverse relationships of the breviate species (22% variation in identity at the 18S) and limited sampling, it is difficult to establish clear co-speciation patterns between members of the phylum.

The diverse nature of the *Desulfovibrionaceae* and *Arcobacteraceae* across phylogenomic trees (Fig. 3A) suggests that their co-occurrence with breviates may not be the result of co-speciation, but rather a propensity or metabolic capabilities for these lineages to engage in interactions with eukaryotes. Indeed, *Arcobacteraceae* species have been shown to associate with other eukaryotes as extracellular epibionts or ectosymbionts[17, 71, 95], and in some cases, these associations are transient[71]. *Desulfovibrionaceae* are also metabolic partners of protists[68, 73]. Even *Terasakiella* species, like *Terasakiella pusilla* (known as *Oceanospirillum pusillum* or *Spirillum pusillum)* was isolated from a marine shellfish[88] and other species from the Rhodospirillales order have been found in association with other eukaryotes[96, 97].

Here, we present the microbial complement of five laboratory-maintained protist microcosms. We demonstrate that protist growth is influenced by the metabolism of the bacteria in the consortium based on the availability of different electron acceptors like sulfate and nitrate. Our findings corroborate previous observations of the *Arcobacter:Lenisia* microcosm, and expand this association to five diverse breviate species, identifying *Desulfovibrionaceae* and *Terasakiella* as new potential partners. Several questions remain about the metabolic interactions between anaerobic breviates and bacteria with respect to the contact-dependence, metabolic nature and cellular biological nature of these interactions. By increasing the sampling of known breviate diversity, it is evident that, much like *Lenisia limosa*, breviate anaerobic growth is influenced by its surrounding prokaryotic community. These interactions likely depend on the metabolism of the bacterial community and availability of electron acceptors and not necessarily on the taxonomic affiliation of the bacteria. Future research should focus on how these organisms might interact in the natural environment and whether this relationship has evolved into a specialized contact-dependent interaction. Furthermore, distinct pathways of adaptation to anoxia in endobiotic, marine, and freshwater environments, where predation, competition, and nutrient availability vary, might also play a role.

## Supporting information

Supplementary Discussion

Supplementary DataFile S1

Supplementary DataFile S2

Supplementary DataFile S3

Supplementary DataFile S4

Supplementary DataFile S5

Supplementary DataFile S6

Supplementary DataFile S7

## ACKNOWLEDGMENTS

The authors would like to thank Yana Eglit, Alastair Simpson and Andrew Roger for material and helpful discussions; Humberto Itriago, Marco Fantini, Anna Castellet and Olena Starostina-Hommel for technical support. This project was supported by funds from the Swedish Research Council Swedish Research Council (Vetenskapsrådet starting grant 2020-05071), the European Research Council (ERC) under the European Union’s Horizon 2020 research and innovation programme (grant agreement ERC Starting grant 101078476 to CWS), the Royal Physiographic Society of Lund grant 43432 (to KIAC) and 44180 (to JB) and Jörgen Lindström’s Foundation grant (to KIAC). The authors acknowledge support from the National Genomics Infrastructure in Stockholm, funded by Science for Life Laboratory, the Knut and Alice Wallenberg Foundation, and the Swedish Research Council. Computational resources and data handling were enabled by the National Academic Infrastructure for Supercomputing in Sweden (NAISS), partially funded by the Swedish Research Council through grant agreement no. 2022-06725, under projects NAISS 2023/5-47, NAISS 2023/6-39, NAISS 2024/5-77, and NAISS 2024/6-43, with access to the UPPMAX and PDC computational infrastructures.

## DATA AVAILABILITY STATEMENT

Amplicon reads, metagenomic reads and assembled genomes of the prokaryotes in the breviate microcosm have been deposited to NCBI in BioProject PRJNA1084235. Descriptions of the datasets are available Supplementary Datafile S3 and S6. Breviate 18S sequences have been deposited under the Genbank Accessions PV036398 (PCE), PV036399 (FB10N2), PV036400 (LRM1b) and PV036401 (LRM2N6). Metagenome assemblies, metagenome-assembled genomes (MAGs) generated through anvi’o binning and reassembled using Trycycler, gene calls, taxonomic classification, metabolic annotation, and quality assessment, raw microscopy pictures and flowCAM data can be found in figshare (figshare.com/s/6a5781edc1a4e0e722bc). Scripts for the flowCAM data classifier can be found https://github.com/theLabUpstairs/FlowCam_Image_Classifier.

