## Supplementary Discussion for "Anaerobic protist survival in microcosms is dependent on microbiome metabolic function"

### SUPPLEMENTARY MATERIALS AND METHODS

#### DNA isolation and preparation for metagenomic sequencing

Cells were collected by centrifugation at 2500 x g for 5 min, and the cell pellet was resuspended in 500 µl of extraction buffer (10 mM Tris-HCl pH 8.0, 10 mM EDTA pH 8.0 and 2% SDS). Proteinase K (20 mg/ml) (ThermoScientific AM2542, Germany) was added to the solution to a final concentration of 770 µg/ml. The tubes were mixed by inversion and incubated at 56°C for 3 h. The DNA was extracted by standard phenol:chloroform extraction using equal volume phenol: chloroform: isoamyl alcohol (ThermoScientific 327111000, Israel) followed by a chloroform extraction to remove residual phenol. The DNA was precipitated with 1/10 volume of 3 M sodium acetate buffer solution pH 5.2 (ThermoScientific R1181, Lithuania) followed by the addition of 2.5 volumes of ice-cold absolute ethanol. The samples were mixed and stored at -20° overnight. To recover the DNA, the samples were collected by centrifugation at 13000 x g for 30 min at 4°C and the resulting pellet was washed with 70% ethanol twice by collection at 13000 x g for 30 min at 4°C and ultimately resuspended in 10 mM Tris buffer, pH 8.0. DNA for metagenome sequence was isolated from PCE, FB10N2, LRM1b, LRM2N6 and *Pygsuia biforma* breviate cultures, cultured in 50 ml conical tubes with 45 ml of LBSW at room temperature for seven days (microaerophilic). For nanopore sequencing, the DNA was treated with 1 mg/ml RNase A (ThermoScientific LT-02241, Lithuania), followed by incubation at room temperature for 10 min. The DNA was quantified with Qubit dsDNA HS assay kit (Invitrogen Q32854, Eugene, Oregon, USA).

#### 18S rRNA gene amplification and sequencing

To recover the 18S rRNA gene sequences of the undescribed breviates, we isolated the DNA from PCE, FB10N2, LRM1b and LRM2N6 breviate cultures as described above. The 18S rRNA gene was amplified from 1 ng of DNA using eukaryote-specific primers EukA and EukB [1] and GoTaq G2 Green Master Mix (Promega M7823, Madison, WI, USA). The thermocycler conditions are outlined in Supplementary Datafile S3. PCR products were separated on a 1% agarose gel, and the products of ~1800 bp were purified using the Nucleospin Gel and PCR clean-up (Macherey-Nagel 740609.50, Germany). Fragments were cloned into the pGEM-T easy vector systems (Promega A3600, Madison, WI, USA) and transformed into chemically competent DH5α *Escherichia coli* chemical competent cells using standard methods [2]. Plasmids were isolated from positive clones using Nucleospin plasmid purification (Macherey-Nagel 740588.50, Germany) and sequenced using Sanger sequencing (Eurofins).

**Amplicon sequencing**

To explore the changes for the prokaryotic community in response to nitrate under microaerophilic conditions, four biological replicates of each breviate microcosm were grown in LBSW or LBSW-NIT (supplemented with 2 mM $KNO_3$) and allowed to acclimatize for 7 d at room temperature. Thereafter cells were passed once more into LBSW or LBSW-NIT and grown for 7 days. DNA was obtained as described above. For the anoxic data, microaerophilic breviate cultures were inoculated in 75 ml of LBSW-HEPES (2 mM HEPES, pH 8.0) or LBSW-NIT-HEPES in 100 ml serum flasks in three biological replicates. Unlike the microaerophilic conditions, we supplemented the anoxic cultures with prey bacteria (*Klebsiella pneumoniae,* 1 x $10^9$ cells/ml) because the prokaryotic community did not grow to sufficient density under anoxic conditions to maintain breviate growth. We supplemented with HEPES so as to best compare with the *Lenisia limosa* study [3]. Samples were incubated for 7 d at room temperature, and DNA was extracted from 250 µl of each microcosm. Because of these differences in experimental design, we do not compare the microaerophilic and anoxic data.

The V4 region of the 16S rRNA gene was amplified from each biological replicate in three technical replicates using the barcoded primers 515F [4] and 806R [5]. The gene was amplified with Phire Hot Start II DNA polymerase (ThermoScientific F122S, Lithuania), dNTP mix (VWR 733-1363, Denmark) and 2 µl of 1 ng/µl DNA, according to the protocol of the polymerase manufacturer. For details about primers and PCR conditions see Supplementary Datafile S2. Technical replicates were pooled together and purified with AMPure XP beads (Beckman Coulter A63880, Brea, California, USA), the DNA was eluted in 5 mM Tris-HCl pH 8.5. Library preparation and sequencing with MiSeq paired-end (2 x 300 bp) (Illumina) were performed by Eurofins with their NGSelect Amplicon 2nd PCR service. Adapter sequence removal and read merging were performed by Eurofins using Cutadapt v2.7 [6] and FLASH v2.2.00 [7], respectively. Raw reads were deposited to NCBI, accession numbers can be found in Supplementary Datafile S3.

The resulting data was processed using the qiime2 (v 2023.9) pipeline [8]. We imported the reads as single end sequences, demultiplexed the data and denoised with DADA2 pipeline, we truncated the sequences at position 250 [8, 9] and filtered out reads that were in less than 3 samples and that have a frequency lower than 500 [8]. In addition we excluded the f__Enterobacteriaceae taxa corresponding to the food source that was added to the cultures. We explored the taxonomic composition of the samples using the pretrained silva 138 classifier [10, 11]. We used q2-gcn-norm plugin to normalize the data to copy number variation based on the rrnDB database (version 5.6) (https://github.com/Jiung-Wen/q2-gcn-norm). We converted the absolute abundance to relative abundance [8] and visualized the data in R.

Differential abundance (Supplementary Datafile S3) across conditions was evaluated with ANCOM [12].

### Amplicon sequencing diversity analysis

ASV data were further analyzed in R (v4.2.2) for compositional analysis of relative abundance and visualization of alpha and beta diversity. Prior to calculating alpha diversity metrics, the data were rarefied to 40620 reads per sample. The alpha diversity metrics included Shannon diversity, which was used to calculate Pielou's evenness as a complementary measure. Rarefaction and calculation of Shannon diversity were performed using the phyloseq package (v1.48.0). Statistical analysis of alpha diversity was performed to test for differences between conditions. Normality was tested with the Shapiro-Wilk test, and due to non-normal distributions, significant differences between conditions were identified using the Kruskal-Wallis test followed by Dunn's test with Bonferroni correction (Supplementary datafile S3). Beta diversity was assessed using the Bray-Curtis dissimilarity metric, and Principal Coordinates Analysis (PCoA) was performed for visualization using the phyloseq package. Pairwise differences between conditions were tested using PERMANOVA via the pairwise.adonis2 function from the vegan package (v2.6.8). Statistical significance was determined at $p \leq 0.05$, with $R^2$ values and F-statistics reported.

### Using DiSCo to predict dissimilatory sulfate reduction metabolism

To identify genes associated with dissimilatory sulfate reduction in the MAGs, DiSCo (version 1.0.0) was used, a Perl 5-based tool designed to automatically detect and classify proteins involved in Dsr-dependent dissimilatory sulfur metabolism (Supplementary Datafile S6). Predicted proteins were analyzed using DiSCo with a predefined Hidden Markov Model (HMM) library and an automated filtering step to retain high-confidence protein predictions [13].

### *Metagenomic sequencing, assembly and binning*

Long-read metagenomic sequencing of PCE, LRM1b, LRM1N6, FB10N2 microcosm DNA was performed by the National Genomics Infrastructure Sweden with ONT ligation kit SQK-LSK109 on one ONT PromethION FLO-PRO002 flowcell. Demultiplexing, adaptor trimming and basecalling in super high accuracy was performed using Guppy 6.1.5 (dna_r9.4.1_450bps_hac_prom) . Sequencing of *Pygsuia biforma* DNA was performed in-house with ONT ligation kit SQK-NBD114.24 and sequenced on one ONT PromethION FLO-PRO114M flowcell. Demultiplexing, adaptator trimming and basecalling was performed in super high accuracy using Dorado 0.7.1 (r1041_e82_400bps_sup_v5.0.0). Reads were trimmed with chopper (-q 9 -l 500) https://github.com/wdecoster/chopper. DNA from each microcosm was assembled independently using Flye 2.9.1 (--meta)[15]. Quality was checked using metaquast 5.2[16]. Reads of each microcosm were mapped onto the corresponding assembly using minimap2 2.24-r1122[17] and samtools 1.14[18]. Contigs clustering, manual binning and gene calls were performed using anvi'o 8[19].

### Bacterial isolation and genome sequencing

*Desulfovibrio glucosivorans* PB2 and *Pseudodesulfovibrio salinus* PB1 and LRM1 were isolated from the *P. biforma* microcosm, by diluting the culture and spreading it on SRBS (to 1 L: 2.32 g of Sulfate Reducing Broth Base Millipore 28228, India, 10 g of sodium thiosulfate, 4 ml of of 60% sodium DL-lactate solution Sigma-aldrich L4263, St. Louis, MO, USA, 33 g of instant ocean and 2% agar) and MBS (For 1 L: 37.4 g of Marine Broth BD Difco 2216, France, 4 ml of 60% sodium DL-lactate solution, and 2% agar. Adjust pH 7.2-7.5) agar plates, respectively. Plates were incubated in a glove box under an 80% $N_2$, 10% $CO_2$ and 10% $H_2$ atmosphere at room temperature for around 7 d. To obtain pure cultures, single colonies were re-streaked in SRBS and MBS agar plates and subsequently grown in SRBS and MBS liquid media (without agar) for DNA purification. All the incubations were done in a glove box at room temperature. The DNA from *Desulfovibrio glucosivorans* PB2 was purified using Gene Jet kit for Gram negative bacteria (Thermo Scientific K0721, Lithuania), and further concentrated with AMPure XP beads according to the manufacturer's instructions. The DNA from *Pseudodesulfovibrio salinus* PB1 was obtained with the phenol-chloroform method described above.

*Terasakiella halodenitrificans* LRM1 was isolated from the LRM1b microcosm, by diluting the culture and spreading it on marine broth (MB) plates supplemented with 2 mM $KNO_3$. The inoculated plate was incubated in microaerophilic conditions (Anaerocult C, Millipore 1.32383.0001, Germany, in a BD BBL™ GasPak™ jar) for 96 h at room temperature. To obtain pure cultures, single colonies were re-streaked in MB and subsequently grown in MB liquid media for DNA purification. All the incubations were done in microaerophilic conditions. The DNA was purified with phenol-chloroform. DNA was quantified with Qubit dsDNA HS assay kit. DNA was sent to Eurofins for bacterial genome sequencing using Oxford Nanopore Technology. Bacterial genome assemblies and quality control of the assemblies were done by Eurofins: short and low-quality reads from the raw nanopore sequencing were removed using Filtlong v0.2.1 (https://github.com/rrwick/Filtlong). Bacterial genome de novo assemblies were done using Flye v2.9.3 [20], and the resulting contigs were polished with Medaka v1.8 (https://github.com/nanoporetech/medaka). The quality of the assembled genomes were assessed using various tools, including QUAST v5.2 [21], CheckM2 v1.0.1 [22] and Mash v2.3

[23]. To ensure the purity of the samples, the reads were mapped onto the assembly with minimap2 v2.24 [17] and to call variations within the assembled genome Clair3 v1.0.4 was used [24].

**16S rRNA phylogenetic trees**

The 16S sequence from each bacterium of interest was extracted from the MAGs and from select type species from each of the major lineages under investigation were used as a query against the SILVA SINA search-and-classify [6] tool and retrieved 20 neighbours using a 90% cut-off value. These results were parsed into two datasets for each lineage: a dataset with only described bacteria and a dataset including environmental sequences. Each dataset was aligned using SINA to the global SSU alignment and gaps were removed using Wasabi default settings for each of the lineages of interest *Desulfovibrionaceae*, *Arcobacteraceae*, and *Terasakiella*. Phylogenies were inferred using IQTREE v2.0 [25, 26] under the best scoring model of evolution decided by ModelFinder and 100 non-parametric bootstraps (-b 100) (Supplementary Datafile S7).

***Average Nucleotide Identity (ANI) comparisons between genomes***

To perform the taxonomic assignment of *Arcobacteraceae, Desulfovibrionaceae* and *Terasakiella* genomes was performed using GTDBtk 2.4.0 [27]. To compare ANI between the new genomes, we used the EZBioCloud ANI calculator [28] (Supplementary Datafile S6). This allowed us to classify MAGs from different microcosms into the same species. For example *Halarcobacter azotofixans* PCE2 and *Halarcobacter azotofixans* FB1 (ANI of 99.49%), *Maridesulfovibrio spiralis* PCE5 and *Maridesulfovibrio spiralis* LRM2 (ANI of 99.46%), and *Halarcobacter ibericus* PCE3 and *Halarcobacter ibericus* 5MF (ANI of 94.66%). In addition, this method was used to compare *Desulfovibrio glucosivorans* PB2 with *Desulfovibrio glucosivorans* DMSS-1 (Reference genome: 2576861818) which is not present in the GTDB, but was investigated due to the high percent of identity of their 16S rDNA sequences (ANI of 97.61%).

***Fluorescence in situ hybridization***

Breviate cultures were grown on a slide and incubated overnight in a moist chamber in an anaerobic jar under anaerobic conditions with Anaerocult A (Millipore 113829, Sparks, Maryland, USA, in a BD BBL™ GasPak™ jar). Breviate cells were fixed with a final concentration of 4% formaldehyde (ThermoScientific 28906, Rockford, IL, USA) for 15 min in multiwell slides. Slides were rinsed in 2 × 50 mL distilled water for a total of 3 min and then air-dried. Slides were immersed in 50%, 80% and 100% ethanol for 3 min in each tube and air-dried. Dried cells on the slide were incubated with hybridization buffer (20 mM Tris-HCl, pH

7.6, 0.01% SDS, 900 mM NaCl), 5 ng/µ of FISH probe and an appropriate concentration of formamide (ThermoScientific 17899, Rockford, IL, USA, Supplementary Datafile S2), in a moist chamber at 46°C for 2 h [29, 30].

FISH probes were synthesized commercially (Supplementary Datafile S2), with identical fluorophores at the 5´- and 3´-end of the oligonucleotide [31]. After incubation, the slides were rinsed and incubated with 50 ml pre-warmed washing buffer (20 mM Tris-HCl, pH 7.5, 0.01% SDS, 5 mM EDTA and a suitable NaCl concentration depending on the %FA in the hybridization buffer, Supplementary Datafile S2) at 48°C for 30 min, and subsequently rinsed in distilled water for 40 s and air-dried [29, 30]. DAPI was added to each well with a final concentration of 2 µg/ml, slides were rinsed with distilled water and air-dried. Slides were mounted with SlowFade Diamond Antifade Mountant (ThermoScientific P36970, Eugene, Oregon, USA). Cells were imaged on a Zeiss Axio Imager.z2 microscope, using a Plan-Apochromat 100×/1.40 Oil Ph 3 M27 immersion oil objective lens. The following filters were used: set 38 for Atto 488 (excitation, BP 470/40 nm; emission, BP 525/50 nm), set 31 for Atto 550 (excitation, BP 565/30 nm; emission, BP 620/60 nm), set 50 for Atto 633 (excitation, BP 640/30 nm; emission, BP 690/50 nm) and set 49 for DAPI (excitation, G 365 nm; emission, BP 445/50 nm). Images were processed using linear adjustments (e.g., brightness/contrast) in fiji.

### SUPPLEMENTARY RESULTS & DISCUSSION

All the *Arcobacteraceae*, *Desulfovibrionaceae* and *Terasakiella* genomes described here encode for bacterial chemotaxis and flagellar proteins, this implies a chemosensory system similar to the one happening in *Escherichia coli*. The pathway is composed of chemoreceptors, the histidine protein kinase chemotaxis protein (CheA) and two diffusible response regulators (CheY and CheB). CheY controls flagellar motor switching, whereas CheB controls chemoreceptor adaptation [32]. The encoded genes might be relevant for bacterial motility and invasion, and can contribute to syntrophic growth, as the flagella can facilitate chemotaxis to the syntrophic partner [3, 33]. There might be other genes and pathways besides the ones described here involved in the interactions with breviates, similar to the genes involved in colonization or virulence like the ones used by their animal-pathogenic relatives [34, 35].

### SUPPLEMENTARY FIGURES & TABLES

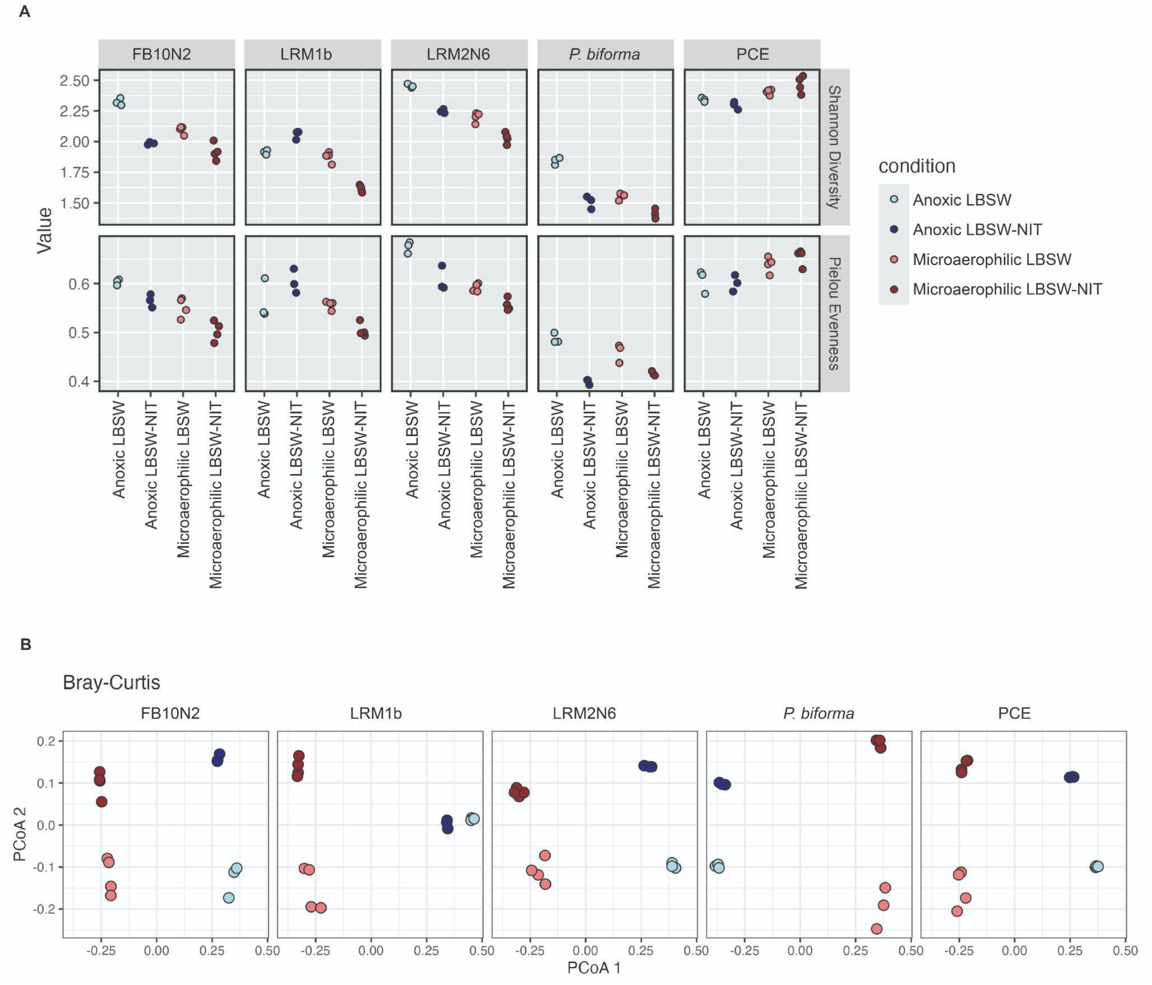

**Supplementary figure S1: Differences in alpha and beta diversity of the bacterial community in the breviate microcosm.** a) Alpha diversity plots of each breviate microcosm growing in anoxic or microaerophilic conditions, with or without nitrate, from left to right: FB10N2, LRM1b, LRM2N6, *P. biforma* and PCE. In the top, Shannon diversity, in the bottom, Pielou eveness. In terms of alpha diversity, Shannon's index was slightly higher in anoxic samples with sulfate for FB10N2, LRM2N6 and *P. biforma* while Shannon's index was higher in anoxic samples supplemented with nitrate for LRM1b. For PCE, the Shannon index was higher in microaerophilic samples with nitrate. In all the cases, the differences in Shannon's index is explained by the evenness, as observed with Pielou's, and not by the richness of the species. b) Beta diversity PCoA plots of each breviate microcosm, same order and conditions as for alpha diversity. The comparisons among communities were made using Bray-Curtis, the results indicated that all conditions (anoxic or microaerophilic, with or without nitrate) in all the samples differed significantly, with the exception of LRM1b anoxic samples with and without nitrate that clustered together in the PCoA plot, suggesting that the presence of sulfate in the anoxic sample is not changing the species. For statistical comparisons see Supplementary Datafile S3.

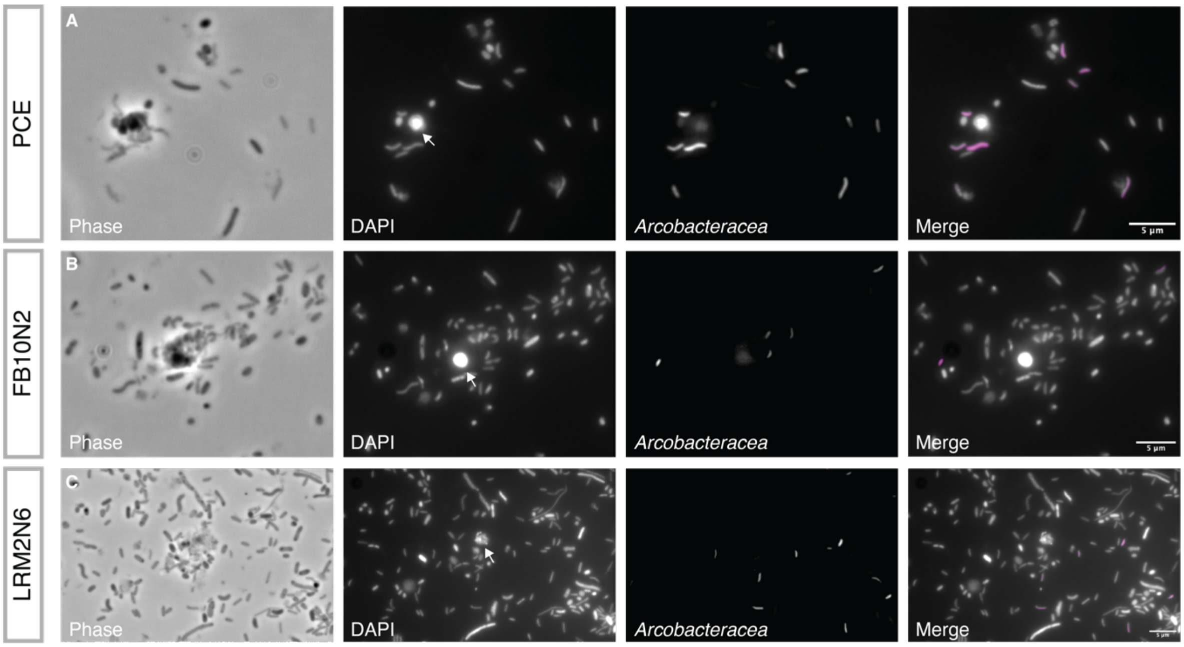

**Supplementary figure S2: Fluorescence *in situ* hybridization (FISH) suggests a direct interaction of PCE, FB10N2 and LRM2N6 with *Arcobacteracea*e.**

a) PCE, b) FB10N2 and c) LRM2N6 breviate microcosm were incubated on a slide overnight in anoxic conditions. Cultures were fixed (4% formaldehyde) and hybridized with 16S rRNA probes (20% formamide): Arc1430-Atto 488 and Arc94-Atto 488 targeting *Arcobacteraceae* cells, then stained with DAPI. Panels from left to right: Phase, DNA stained with DAPI (white arrows pointing to the breviate nucleus), *Arcobacteraceae* probes, and merge channel of DAPI (gray) with FISH probes (magenta). Scale bar 5 µm.

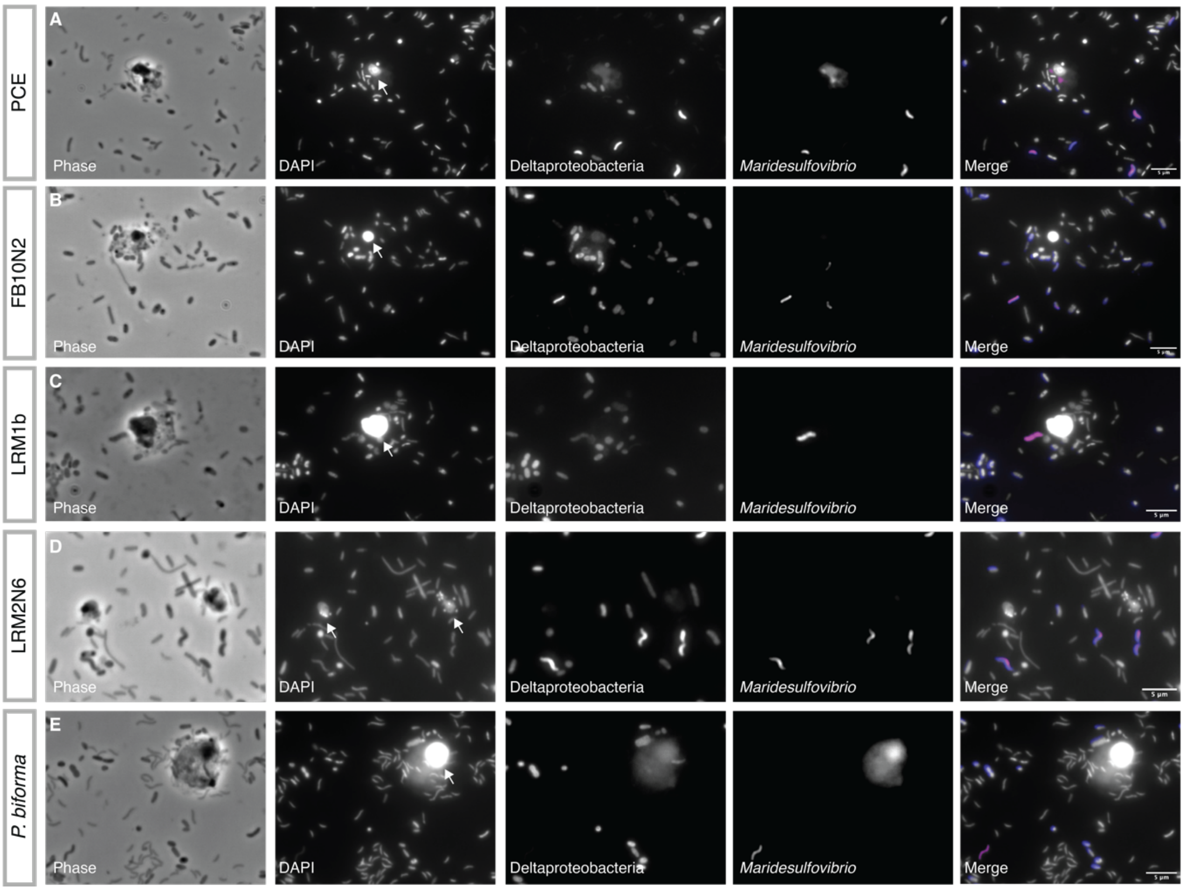

**Supplementary figure S3: Fluorescence *in situ* hybridization (FISH) suggests a direct interaction of PCE, FB10N2, LRM1b, LRM2N6 and *P. biforma* with *Maridesulfovibrio* species.**

a) PCE, b) FB10N2, c) LRM1b, d) LRM2N6 and e) *P. biforma* breviate microcosm were incubated on a slide overnight in anoxic conditions. Cultures were fixed (4% formaldehyde) and hybridized with 16S rRNA probes (35% formamide): MD1 Atto 633 + MD2 Atto 633 targeting different *Maridesulfovibrio* species and delta495a Atto 550 targeting Deltaproteobacteria, then stained with DAPI. Panels from left to right: Phase, DNA stained with DAPI ( white arrows pointing to the breviate nucleus), Deltaproteobacteria probes, *Maridesulfovibrio* probes, and merge channel of DAPI (gray) with FISH probes (Deltaproteobacteria in blue, and *Maridesulfovibrio* in magenta). Scale bar 5 µm.

259

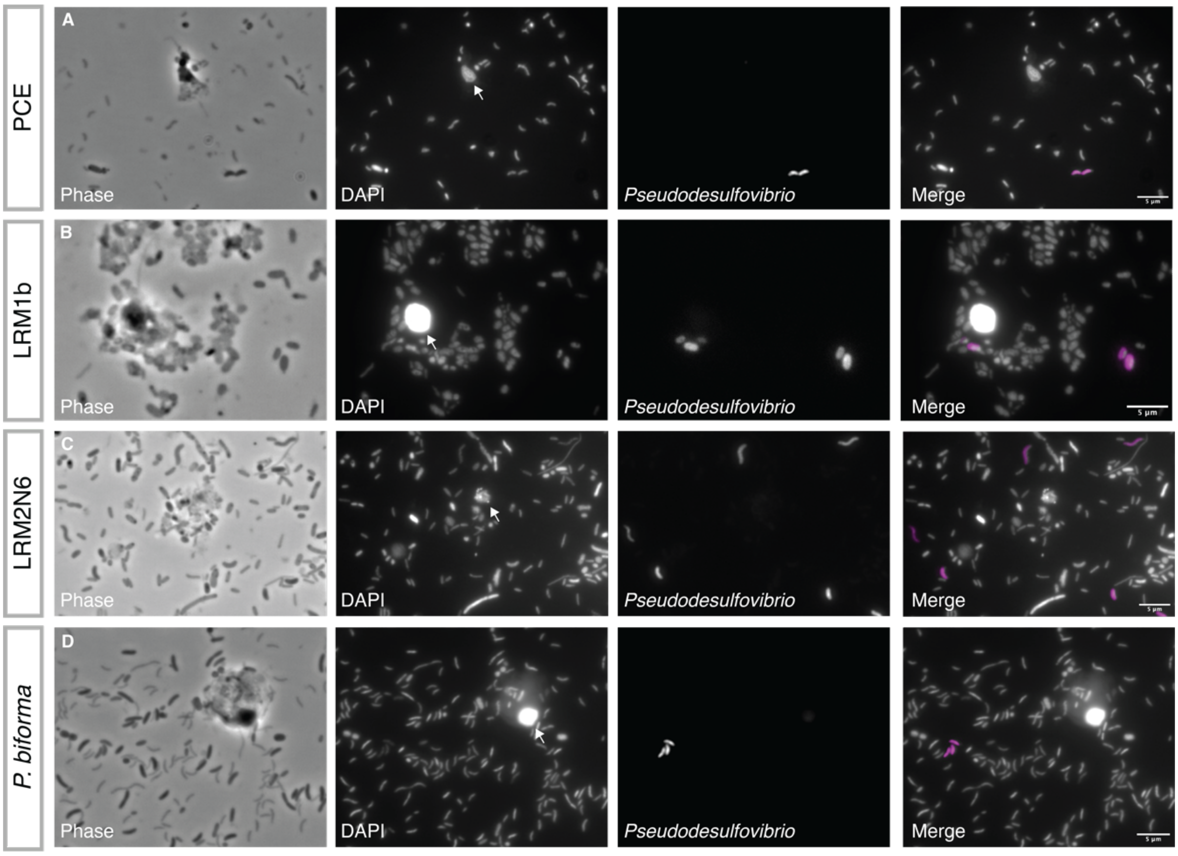

**Supplementary figure S4: Fluorescence *in situ* hybridization (FISH) suggests a direct interaction of PCE, LRM1b, LRM2N6 and *P. biforma* with *Pseudodesulfovibrio* species.**

a) PCE, b) LRM1b, c) LRM2N6 and d) *P. biforma* breviate microcosm were incubated on a slide overnight in anoxic conditions. Cultures were fixed (4% formaldehyde) and hybridized with 16S rRNA probes (20% formamide): PD1-Atto 633 targeting different *Pseudodesulfovibrio* species, then stained with DAPI. Panels from left to right: Phase, DNA stained with DAPI (white arrows pointing to the breviate nucleus), *Pseudodesulfovibrio* probes, and merge channel of DAPI (gray) with *Pseudodesulfovibrio* probes (in magenta). Scale bar 5 µm.

260

**Table S1: Etymology of new prokaryotic taxa in breviate microcosms submitted to SeqCode**

| Proposed name | Strain | New genus or species | Etymology | Meaning |
|---|---|---|---|---|
| *Marinarcus sediminensis* | PCE1 | species | se.di.mi.nen’sis N.L. neut. n. *sediminensis*, from L. *sedimen*, sediment, and *-ensis*, suffix denoting origin | Bacterium isolated from sediment |
| *Halarcobacter azotofixans* | PCE2, FB1 | species | a.zo.to.fix’ans. N.L. neut. n. *azotofixans*, from L. *azote*, nitrogen, and L. *fixans*, fixing | Bacterium with metabolic potential for nitrogen-fixation |
| *Halarcobacter ibericus* | PCE3, 5MF | species | i.be'ri.cus. N.L. masc. adj. *ibericus*, of or belonging to Iberia | Bacterium originated from the Iberian region. Description in preparation (Aguilera-Campos et al.) |
| *Maridesulfovibrio spiralis* | PCE5, LRM2 | species | spi.ra’lis. N.L. neut. n. *spiralis*, from L. *spiralis*, coiled or spiral-shaped | Spiral-shaped bacterium |
| *Pseudodesulfovibrio salinus* | PB1 | species | sa.li’nus. N.L. masc. adj. *salinus*, from L. *salinus*, salty | Bacterium that lives in saline environments |
| *Arcobacter denitrificans* | LRM1 | species | de.ni.tri’fi.cans. N.L. neut. n. *denitrificans*, denitrifying | Bacterium with metabolic potential for denitrification |
| *Terasakiella halodenitrificans* | LRM1 | species | ha.lo.de.ni.tri’fi.cans. N.L. neut. n. *halodenitrificans*, salt-denitrifying | A halophilic bacterium capable of denitrification |
| *Arcobacter sinusprincipis* | 7KC | species | si.nus.prin.ci’pis. N.L. masc. adj. *sinusprincipis*, originating from Prince Cove. | Bacterium originated from Prince Cove. Description in preparation (Aguilera-Campos et al.) |
| *Maridesulfovibrio pacificus* | FB2 | species | pa.ci’fi.cus. N.L. masc. adj. *pacificus*, from the Pacific | Bacterium originated from the Pacific Ocean |
| *Thalassodesulfovibrio aquaticus* | PCE4 | genus | Tha.las.so.de.sul.fo.vi’bri.o. N.L. masc. n. *Thalassodesulfovibrio*, | A marine bacterium capable of sulfate reduction |

| Proposed name | Strain | New genus or species | Etymology | Meaning |
|---|---|---|---|---|
| | | | from G. *thalassa*, sea, L. *de*, from, L. *sulfur*, sulfur, and N.L. *vibrio*, a curved bacterium | |
| *Thalassodesulfovibrio aquaticus* | PCE4 | species | a.qua'ti.cus. N.L. masc. adj. *aquaticus*, from L. *aqua*, water | |

### DESCRIPTION OF SUPPLEMENTARY DATA FILES

**Supplementary Datafile S1 (.xlsx)**

This spreadsheet file includes multiple sheets with information about the sampling sites for the microcosms and the coordinates for the 18S sequences found in publicly available data.

**Supplementary Datafile S2 (.xlsx)**

This spreadsheet contains multiple sheets summarizing the primer sequences, PCR conditions, FISH probe sequences and conditions and phylogenetic tree information.

**Supplementary Datafile S3 (.xlsx)**

This spreadsheet includes multiple sheets with information about the 16S amplicon analysis in the different breviate cultures grown with different conditions including the metadata file and accession numbers to the raw reads in NCBI. It also includes information about statistical analysis between the conditions (ANCOM) and diversity analysis .

**Supplementary Datafile S4 (.xlsx)**

This spreadsheet shows the raw FlowCAM data for the breviate growth curve experiment in dSW (noE- no electron acceptor), dSW-NIT (nitrate) and dSW-SULF (sulfate) in anoxic conditions. Protists concentration was calculated at days 0, 3, 5, 7 and 10, using flowCAM.

**Supplementary Datafile S5 (.html)**

This html file shows the workflow for calculating growth rates and statistical analysis of the FlowCAM cell count data.

**Supplementary Datafile S6 (.xlsx)**

This spreadsheet contains multiple worksheets detailing the prokaryotic assembled genomes, Genbank accession numbers, the taxonomy classification using GTDBtk, average nucleotide identity comparisons between our assembled genomes and metabolic reconstruction.

**Supplementary Datafile S7 (.pdf)**

This PDF shows the phylogenetic analysis of the 16S from breviate-associated bacteria from the *Arcobacteraceae*, *Desulfovibrionaceae,* and *Terasakiella* genomes. For *Arcobacteraceae* and *Desulfovibrionaceae* we performed an additional analysis of only cultured representatives. Breviate-associated sequences are coloured. Uncultured sequences from public databases are shown in grey.

**Figshare contents**

Link for review: figshare.com/s/6a5781edc1a4e0e722bc

- ➔ **16S_Breviate_Associated_Trees** - 16S rRNA phylogenetic trees for breviate-associated bacteria, including alignments, IQ-TREE outputs, and colorized tree visualizations.
- ➔ **18S_Breviate_Associated_Trees** - 18S rRNA sequences from multiple sources, alignments processed with SSU-align and SSU-mask, and phylogenetic trees generated using IQ-TREE.
- ➔ **Arcobacter_EP1_Annotation** - Gene prediction and annotation of *Arcobacter* sp. EP1 (GCA_001655195.1) using Prokka.
- ➔ **DIC_breviates** - Differential interference contrast (DIC) microscopy images of breviates.
- ➔ **FlowCam_classified_images** - classified FlowCam generated images, used for evaluating breviate concentrations.
- ➔ **FISH_breviate_microscosm** - Fluorescence *in situ* hybridization (FISH) images of breviate microcosms.
- ➔ **GTDBtk_Classification_and_Phylogeny** - GTDB-tk classification and phylogenetic placement of *Arcobacteraceae*, *Desulfovibrionaceae*, and *Terasakiella* genomes.
- ➔ **Metagenomic_MAGs** - Metagenome-assembled genomes (MAGs) generated through anvi'o binning and reassembled using Trycycler, with taxonomic classification, metabolic annotation, and quality assessment.
