## Supplementary DataFile S5 for "Anaerobic protist survival in microcosms is dependent on microbiome metabolic function": Supplementary_DataFileS5_anova_growth_rates.html

Pygsuia and LRM1b - Growth Data


#### Table of contents

- Setup
- Growth parameters
  - Initial growth rate
- Comparing treatments and strains
  - ANOVA
  - Check assumptions
  - Post-Hoc tests
- Figures
  - Preliminary figures

### Pygsuia and LRM1b - Growth Data

 Code

Data analysis

Author

Ada Behncké Serra

Published

November 10, 2024

### Setup

Load required packages

```
if(!require(pacman)){install.packages("pacman")} #'pacman' eases loading/installation of packages
p_load(readr, #for data import 
       tidyverse, #for data wrangling and visualisation
       car, #tools and functions for linear regression analysis
       rstatix, #pipe-friendly stats tools
       ggpubr) #for additional plotting aesthetic options
```

Load data

```
Data_Growth <- read_delim("Data/growthcurve_anoxic_Pygsuia_LRM1b_sep2024.csv",
                          delim = ";", escape_double = FALSE, 
                          col_types = cols(
                            Treatment = col_factor(levels = c("Control", "Nitrate", "Sulfate")),
                            Culture = col_factor(levels = c("Pygsuia", "LRM1b")), 
                            Biorep = col_factor(levels = c("A", "B", "C"))
                            ),
                          trim_ws = TRUE)
```

---

### Growth parameters

We start by calculating means and standard error of the mean (SEM) of the technical replicates. W aree interested in keeping the biological replicates separate, as each one may behave differently. While the 95% confidence interval is a more informative measure of where the true mean of the population lies, with such a small sample (n = 3) it will be even more imprecise than the SEM. To give an idea of the goodness of our mean estimate, given our small sample, the SEM will suffice.

```
Data_growth_means <- Data_Growth %>% 
  group_by(Culture, Treatment, Time_d, Biorep) %>% 
  summarise(
    Cell_density_mean = mean(Cells_ul),
    Cell_density_SEM = (sd(Cells_ul)/3)
  )
head(Data_growth_means)
```

```
# A tibble: 6 × 6
# Groups:   Culture, Treatment, Time_d [2]
  Culture Treatment Time_d Biorep Cell_density_mean Cell_density_SEM
  <fct>   <fct>      <dbl> <fct>              <dbl>            <dbl>
1 Pygsuia Control        0 A                   749.             100.
2 Pygsuia Control        0 B                   749.             100.
3 Pygsuia Control        0 C                   749.             100.
4 Pygsuia Control        3 A                 16327.             604.
5 Pygsuia Control        3 B                 16433.             101.
6 Pygsuia Control        3 C                 19684.             495.
```

#### Initial growth rate

To compare growth rates we are interested in the rate of increase at the exponential phase of microbial growth. We do not have the time resolution to fit an exponential curve to our initial data (almost all cultures are peaking already at our second time point, i.e. 3 days), so we are just going to linearly estimate a growth rate that we will call `µ`

We will estimate the initial growth rate as `µ = (Nmax - N0)/(t_Nmax - t0)`, where `Nmax`is the observed peak cell density (cells/mL), `N0`is the initial cell density (cells/mL), `t_Nmax`is the time point at which the peak cell density is reached (d), and `t0`is the initial time point (d = 0). Since t0 is always 0 (in our case), we can ignore the term in the formula.

To estimate the SEM associated with `µ`, we propagate the uncertainties of `Nmax`and `N0` (`t_Nmax` and `t0`do not contribute to the uncertainty propagation because they do not have an associated uncertainty). Observe that, to propagate the uncertainty, we are assuming that N0 and Nmax are independent (which they probably are not entirely), but we will have to live with that.

We start by creating functions to calculate `µ` and estimate its standard error.

```
calc_growthrate <- function (Nmax, Nmin, N0, t_Nmax, t_Nmin) {
  #create an empty vector to fill with values
  mu <- rep(0, length(t_Nmax))
  for (i in length(t_Nmax)) {
     #if t_Nmax is zero, we want to calculate a death rate
    if (t_Nmax[i] == 0) {
      mu[i] <- (Nmin[i] - N0[i])/(t_Nmin[i])
      } 
    else {
      mu[i] <- (Nmax[i] - N0[i])/(t_Nmax[i])
      }
  }
 
  return(mu)
}

calc_growthrate_SEM <- function (Nmax_SEM, Nmin_SEM, N0_SEM, t_Nmax, t_Nmin) {
  #create an empty vector to fill with values
  muSEM <- rep(0, length(t_Nmax))
  for (i in length(t_Nmax)) {
     #if t_Nmax is zero, we want to calculate a death rate
    if (t_Nmax[i] == 0) {
      muSEM[i] <- sqrt((Nmin_SEM[i]^2 + N0_SEM[i]^2)/t_Nmin[i]^2)
      } 
    else {
      muSEM[i] <- sqrt((Nmax_SEM[i]^2 + N0_SEM[i]^2)/t_Nmax[i]^2)
      }
  }
 
  return(muSEM)
}
```

Now we use the functions we defined above to make a table summarising the growth parameters.

```
Growth_params <- Data_growth_means %>% 
  group_by(Culture, Treatment, Biorep) %>% 
  summarise(
    N0 = Cell_density_mean[Time_d == 0], #cell density at time t=0 and its SEM
    N0_SEM = Cell_density_SEM[Time_d == 0],
    Nmax= max(Cell_density_mean), #peak cell density
    Nmin = min(Cell_density_mean), #min cell density
    Nmax_SEM = Cell_density_SEM[which.max(Cell_density_mean)], #corresponding SEM
    Nmin_SEM = Cell_density_SEM[which.min(Cell_density_mean)],
    t_Nmax = Time_d[which.max(Cell_density_mean)], #time of the peak cell density
    t_Nmin = Time_d[which.min(Cell_density_mean)], #time of the min cell density
    µ = calc_growthrate(Nmax, Nmin, N0, t_Nmax, t_Nmin), #apply the function we defined
    µ_SEM = calc_growthrate_SEM(Nmax_SEM, Nmin_SEM, N0_SEM, t_Nmax, t_Nmin)
  )
```

```
`summarise()` has grouped output by 'Culture', 'Treatment'. You can override
using the `.groups` argument.
```

```
head(Growth_params)
```

```
# A tibble: 6 × 13
# Groups:   Culture, Treatment [2]
  Culture Treatment Biorep    N0 N0_SEM   Nmax  Nmin Nmax_SEM Nmin_SEM t_Nmax
  <fct>   <fct>     <fct>  <dbl>  <dbl>  <dbl> <dbl>    <dbl>    <dbl>  <dbl>
1 Pygsuia Control   A       749.   100. 16327.  50.4    604.     7.62       3
2 Pygsuia Control   B       749.   100. 16433. 250.     101.    27.9        3
3 Pygsuia Control   C       749.   100. 19684. 132.     495.    13.5        3
4 Pygsuia Nitrate   A       749.   100.   814.  32.1     40.6    3.77       3
5 Pygsuia Nitrate   B       749.   100.   749.  49.2    100.     8.98       0
6 Pygsuia Nitrate   C       749.   100.   749.  18.9    100.     0.135      0
# ℹ 3 more variables: t_Nmin <dbl>, µ <dbl>, µ_SEM <dbl>
```

---

### Comparing treatments and strains

#### ANOVA

Now we can use ANOVA to compare growth rates between treatments and strains. Since wee have two grouping factors (Culture and Treatment), we will be doing a two way ANOVA. We are also interested in the interaction between the two factors, i.e., whether the treatments affect the strains differently. Keep in mind, though, that we are comparing calculated growth rates with big uncertainties.

```
Growthrate_anova <- aov(µ ~ Culture * Treatment, data = Growth_params)

summary(Growthrate_anova)
```

```
                  Df   Sum Sq  Mean Sq F value   Pr(>F)    
Culture            1  2454504  2454504   11.64  0.00516 ** 
Treatment          2 71699111 35849556  169.97 1.57e-09 ***
Culture:Treatment  2 35144931 17572465   83.31 9.19e-08 ***
Residuals         12  2531026   210919                     
---
Signif. codes:  0 '***' 0.001 '**' 0.01 '*' 0.05 '.' 0.1 ' ' 1
```

#### Check assumptions

We can only trust the results of a model it the data satisfies the assumptions that the model builds on. In the case of an ANOVA, these assumptions are the normality of residuals and the homogeneity of variances.

```
#Normality of residuals check: formally (Shapiro-Wilk's test) and visually

shapiro.test(residuals(Growthrate_anova)) #Shapiro-Wilk test
```

```
    Shapiro-Wilk normality test

data:  residuals(Growthrate_anova)
W = 0.93032, p-value = 0.1967
```

```
qqnorm(residuals(Growthrate_anova)) #plot residuals
qqline(residuals(Growthrate_anova))
```

```
#Homogeneity of variances check: formally (Levene's test)
leveneTest(µ ~ Culture * Treatment, data = Growth_params)
```

```
Levene's Test for Homogeneity of Variance (center = median)
      Df F value Pr(>F)
group  5  0.7608 0.5949
      12
```

Since the p-value of the Shapiro-Wilk normality test is >0.05, we can’t say that the data is not normally distributed. The residuals do not look bad either. The normality assumption is thus satisfied.

Th p-value for the Levene’s test is also >0.05. Thus, we can’t reject the null hypothesis and we consider the variances to be equal amongst groups. The homogeneity of variances assumption is thus satisfied.

#### Post-Hoc tests

Since we found significant differences among groups with the ANOVA test, we use Post-Hoc tests to examine the specific group differences. We will use the Tukey’s HSD method for pairwise comparisons.

```
summary(Growthrate_anova)
```

```
                  Df   Sum Sq  Mean Sq F value   Pr(>F)    
Culture            1  2454504  2454504   11.64  0.00516 ** 
Treatment          2 71699111 35849556  169.97 1.57e-09 ***
Culture:Treatment  2 35144931 17572465   83.31 9.19e-08 ***
Residuals         12  2531026   210919                     
---
Signif. codes:  0 '***' 0.001 '**' 0.01 '*' 0.05 '.' 0.1 ' ' 1
```

```
post_hoc_tukey_Growthrate <- as.data.frame(tukey_hsd(Growthrate_anova))

post_hoc_tukey_Growthrate
```

```
                term          group1          group2 null.value   estimate
1            Culture         Pygsuia           LRM1b          0  -738.5428
2          Treatment         Control         Nitrate          0 -3446.0844
3          Treatment         Control         Sulfate          0  1279.9820
4          Treatment         Nitrate         Sulfate          0  4726.0664
5  Culture:Treatment Pygsuia:Control   LRM1b:Control          0 -1433.9263
6  Culture:Treatment Pygsuia:Control Pygsuia:Nitrate          0 -5652.2814
7  Culture:Treatment Pygsuia:Control   LRM1b:Nitrate          0 -2673.8137
8  Culture:Treatment Pygsuia:Control Pygsuia:Sulfate          0  2443.1037
9  Culture:Treatment Pygsuia:Control   LRM1b:Sulfate          0 -1317.0659
10 Culture:Treatment   LRM1b:Control Pygsuia:Nitrate          0 -4218.3551
11 Culture:Treatment   LRM1b:Control   LRM1b:Nitrate          0 -1239.8874
12 Culture:Treatment   LRM1b:Control Pygsuia:Sulfate          0  3877.0300
13 Culture:Treatment   LRM1b:Control   LRM1b:Sulfate          0   116.8604
14 Culture:Treatment Pygsuia:Nitrate   LRM1b:Nitrate          0  2978.4677
15 Culture:Treatment Pygsuia:Nitrate Pygsuia:Sulfate          0  8095.3851
16 Culture:Treatment Pygsuia:Nitrate   LRM1b:Sulfate          0  4335.2154
17 Culture:Treatment   LRM1b:Nitrate Pygsuia:Sulfate          0  5116.9174
18 Culture:Treatment   LRM1b:Nitrate   LRM1b:Sulfate          0  1356.7478
19 Culture:Treatment Pygsuia:Sulfate   LRM1b:Sulfate          0 -3760.1696
      conf.low   conf.high    p.adj p.adj.signif
1  -1210.24869  -266.83681 5.16e-03           **
2  -4153.47727 -2738.69152 5.52e-08         ****
3    572.58916  1987.37491 1.11e-03           **
4   4018.67355  5433.45931 1.62e-09         ****
5  -2693.46589  -174.38671 2.28e-02            *
6  -6911.82096 -4392.74179 4.32e-08         ****
7  -3933.35329 -1414.27411 1.35e-04          ***
8   1183.56411  3702.64329 3.20e-04          ***
9  -2576.60551   -57.52634 3.86e-02            *
10 -5477.89467 -2958.81549 1.18e-06         ****
11 -2499.42700    19.65218 5.46e-02           ns
12  2617.49041  5136.56959 2.94e-06         ****
13 -1142.67922  1376.39996 9.99e-01           ns
14  1718.92808  4238.00726 4.65e-05         ****
15  6835.84549  9354.92467 7.56e-10         ****
16  3075.67586  5594.75504 8.70e-07         ****
17  3857.37782  6376.45700 1.35e-07         ****
18    97.20819  2616.28737 3.23e-02            *
19 -5019.70922 -2500.63004 4.09e-06         ****
```

We can see that, strictly speaking in terms of p-value, all groups are significantly different except the control vs. sulfate treatment and control vs. nitrate treatment for LRM1b.

However, given that our growth rate estimates are HIGHLY INACCURATE, significant differences with p-values in the order of 10^-2 or 10^-3 should be regarded skeptically, especially if coupled with relatively small effect sizes.

### Figures

#### Preliminary figures

We want to plot the two strains separately, so we will need separate datasets for each strain. We start by preparing these two datasets. We will also want to add some jitter to the x-axis to avoid overlapping of the data points.

```
Data_plot_Pygsuia <- Data_growth_means %>% 
  filter(Culture == 'Pygsuia') %>% 
  mutate(
    Time_d_jittered = Time_d + rnorm(n(), mean = 0, sd = 0.15),  # Jitter on the x-axis
  )


Data_plot_LRM1b <- Data_growth_means %>% 
  filter(Culture == 'LRM1b') %>% 
  mutate(
    Time_d_jittered = Time_d + rnorm(n(), mean = 0, sd = 0.15),  # Jitter on the x-axis
  )
```

Now, we make the plots. We add some jitter to x-axis to avoid overlap between the points

```
ggplot(data = Data_plot_Pygsuia,
       aes(x = Time_d_jittered, y = Cell_density_mean, col = Treatment)) +
  geom_point() +  # Plot jittered points
  geom_line(aes(group = interaction(Biorep, Treatment), color = Treatment)) +
  geom_errorbar(aes(ymin = Cell_density_mean - Cell_density_SEM, 
                    ymax = Cell_density_mean + Cell_density_SEM), 
                width = 0.01) + # Add error bars
  scale_x_continuous(breaks = c(0, 3, 5, 7, 10), #set x-axis breaks
                     labels = c(0, 3, 5, 7, 10)) +
  coord_cartesian(ylim = c(0, 30000)) + #adjust zoom
  scale_y_continuous(breaks = c(0,10000, 20000, 30000),
                     labels = c(0,
                                expression(10^4),
                                expression("2·"~10^4),
                                expression("3·"~10^4)
                                )) +
  labs(x = "Time (d)",
       y = expression("Cell Density (cells·"~mL^-1~")")) +
  theme_pubr()+
  scale_colour_viridis_d(option = "magma",
                         begin = 0, end =  0.7) +
  theme(axis.title.y = element_text(margin = margin(r = 10)),
        axis.title.x = element_text(margin = margin(t = 10)),
        legend.position = "right"
        )
```

```
ggplot(data = Data_plot_LRM1b,
       aes(x = Time_d_jittered, y = Cell_density_mean, col = Treatment)) +
  geom_point() +  # Plot jittered points
  geom_line(aes(group = interaction(Biorep, Treatment), color = Treatment)) +
  geom_errorbar(aes(ymin = Cell_density_mean - Cell_density_SEM, 
                    ymax = Cell_density_mean + Cell_density_SEM), 
                width = 0.01) + # Add error bars
  scale_x_continuous(breaks = c(0, 3, 5, 7, 10), #set x-axis breaks
                     labels = c(0, 3, 5, 7, 10)) +
  coord_cartesian(ylim = c(0, 20000)) + #adjust zoom
  scale_y_continuous(breaks = c(0,10000, 20000),
                     labels = c(0,
                                expression(10^4),
                                expression("2·"~10^4)
                                )) +
  labs(x = "Time (d)",
       y = expression("Cell Density (cells·"~mL^-1~")")) +
  theme_pubr()+
  scale_colour_viridis_d(option = "magma",
                         begin = 0, end =  0.7) +
  theme(axis.title.y = element_text(margin = margin(r = 10)),
        axis.title.x = element_text(margin = margin(t = 10)),
        legend.position = "right"
        )
```


###### Source Code

```
---
title: "Pygsuia and LRM1b - Growth Data"
subtitle: "Data analysis"
author: "Ada Behncké Serra"
date: "today"
date-format: "long"
format: 
    html:
        code-fold: false
        code-tools: true
        code-overflow: wrap
        toc: true
        toc-depth: 6
        self-contained: true
        anchor-sections: true
        smooth-scroll: true
        theme: simplex
editor: visual
---

# Setup

Load required packages

```{r Packages, message=FALSE, warning=FALSE}

if(!require(pacman)){install.packages("pacman")} #'pacman' eases loading/installation of packages
p_load(readr, #for data import 
       tidyverse, #for data wrangling and visualisation
       car, #tools and functions for linear regression analysis
       rstatix, #pipe-friendly stats tools
       ggpubr) #for additional plotting aesthetic options

```

Load data

```{r Load data, message=FALSE, warning=FALSE}

Data_Growth <- read_delim("Data/growthcurve_anoxic_Pygsuia_LRM1b_sep2024.csv",
                          delim = ";", escape_double = FALSE, 
                          col_types = cols(
                            Treatment = col_factor(levels = c("Control", "Nitrate", "Sulfate")),
                            Culture = col_factor(levels = c("Pygsuia", "LRM1b")), 
                            Biorep = col_factor(levels = c("A", "B", "C"))
                            ),
                          trim_ws = TRUE)
```

------------------------------------------------------------------------

# Growth parameters

We start by calculating means and standard error of the mean (SEM) of the technical replicates. W aree interested in keeping the biological replicates separate, as each one may behave differently. While the 95% confidence interval is a more informative measure of where the true mean of the population lies, with such a small sample (n = 3) it will be even more imprecise than the SEM. To give an idea of the goodness of our mean estimate, given our small sample, the SEM will suffice.

```{r Means and SEM, message=FALSE, warning=FALSE}

Data_growth_means <- Data_Growth %>% 
  group_by(Culture, Treatment, Time_d, Biorep) %>% 
  summarise(
    Cell_density_mean = mean(Cells_ul),
    Cell_density_SEM = (sd(Cells_ul)/3)
  )
head(Data_growth_means)
```

## Initial growth rate

To compare growth rates we are interested in the rate of increase at the exponential phase of microbial growth. We do not have the time resolution to fit an exponential curve to our initial data (almost all cultures are peaking already at our second time point, i.e. 3 days), so we are just going to linearly estimate a growth rate that we will call `µ`

We will estimate the initial growth rate as `µ = (Nmax - N0)/(t_Nmax - t0)`, where `Nmax`is the observed peak cell density (cells/mL), `N0`is the initial cell density (cells/mL), `t_Nmax`is the time point at which the peak cell density is reached (d), and `t0`is the initial time point (d = 0). Since t0 is always 0 (in our case), we can ignore the term in the formula.

To estimate the SEM associated with `µ`, we propagate the uncertainties of `Nmax`and `N0` (`t_Nmax` and `t0`do not contribute to the uncertainty propagation because they do not have an associated uncertainty). Observe that, to propagate the uncertainty, we are assuming that N0 and Nmax are independent (which they probably are not entirely), but we will have to live with that.

We start by creating functions to calculate `µ` and estimate its standard error.

```{r Growth rate function}

calc_growthrate <- function (Nmax, Nmin, N0, t_Nmax, t_Nmin) {
  #create an empty vector to fill with values
  mu <- rep(0, length(t_Nmax))
  for (i in length(t_Nmax)) {
     #if t_Nmax is zero, we want to calculate a death rate
    if (t_Nmax[i] == 0) {
      mu[i] <- (Nmin[i] - N0[i])/(t_Nmin[i])
      } 
    else {
      mu[i] <- (Nmax[i] - N0[i])/(t_Nmax[i])
      }
  }
 
  return(mu)
}

calc_growthrate_SEM <- function (Nmax_SEM, Nmin_SEM, N0_SEM, t_Nmax, t_Nmin) {
  #create an empty vector to fill with values
  muSEM <- rep(0, length(t_Nmax))
  for (i in length(t_Nmax)) {
     #if t_Nmax is zero, we want to calculate a death rate
    if (t_Nmax[i] == 0) {
      muSEM[i] <- sqrt((Nmin_SEM[i]^2 + N0_SEM[i]^2)/t_Nmin[i]^2)
      } 
    else {
      muSEM[i] <- sqrt((Nmax_SEM[i]^2 + N0_SEM[i]^2)/t_Nmax[i]^2)
      }
  }
 
  return(muSEM)
}

  
```

Now we use the functions we defined above to make a table summarising the growth parameters.

```{r Summarise Growth Parameters}

Growth_params <- Data_growth_means %>% 
  group_by(Culture, Treatment, Biorep) %>% 
  summarise(
    N0 = Cell_density_mean[Time_d == 0], #cell density at time t=0 and its SEM
    N0_SEM = Cell_density_SEM[Time_d == 0],
    Nmax= max(Cell_density_mean), #peak cell density
    Nmin = min(Cell_density_mean), #min cell density
    Nmax_SEM = Cell_density_SEM[which.max(Cell_density_mean)], #corresponding SEM
    Nmin_SEM = Cell_density_SEM[which.min(Cell_density_mean)],
    t_Nmax = Time_d[which.max(Cell_density_mean)], #time of the peak cell density
    t_Nmin = Time_d[which.min(Cell_density_mean)], #time of the min cell density
    µ = calc_growthrate(Nmax, Nmin, N0, t_Nmax, t_Nmin), #apply the function we defined
    µ_SEM = calc_growthrate_SEM(Nmax_SEM, Nmin_SEM, N0_SEM, t_Nmax, t_Nmin)
  )
head(Growth_params)

```

------------------------------------------------------------------------

# Comparing treatments and strains

## ANOVA

Now we can use ANOVA to compare growth rates between treatments and strains. Since wee have two grouping factors (Culture and Treatment), we will be doing a two way ANOVA. We are also interested in the interaction between the two factors, i.e., whether the treatments affect the strains differently. Keep in mind, though, that we are comparing calculated growth rates with big uncertainties.

```{r ANOVA}

Growthrate_anova <- aov(µ ~ Culture * Treatment, data = Growth_params)

summary(Growthrate_anova)

```

## Check assumptions

We can only trust the results of a model it the data satisfies the assumptions that the model builds on. In the case of an ANOVA, these assumptions are the normality of residuals and the homogeneity of variances.

```{r Assumption Checks}

#Normality of residuals check: formally (Shapiro-Wilk's test) and visually

shapiro.test(residuals(Growthrate_anova)) #Shapiro-Wilk test
qqnorm(residuals(Growthrate_anova)) #plot residuals
qqline(residuals(Growthrate_anova))

#Homogeneity of variances check: formally (Levene's test)
leveneTest(µ ~ Culture * Treatment, data = Growth_params)

```

Since the p-value of the Shapiro-Wilk normality test is \>0.05, we can't say that the data is not normally distributed. The residuals do not look bad either. The normality assumption is thus satisfied.

Th p-value for the Levene's test is also \>0.05. Thus, we can't reject the null hypothesis and we consider the variances to be equal amongst groups. The homogeneity of variances assumption is thus satisfied.

## Post-Hoc tests

Since we found significant differences among groups with the ANOVA test, we use Post-Hoc tests to examine the specific group differences. We will use the Tukey's HSD method for pairwise comparisons.

```{r Post-hoc TukeyHSD}

summary(Growthrate_anova)

post_hoc_tukey_Growthrate <- as.data.frame(tukey_hsd(Growthrate_anova))

post_hoc_tukey_Growthrate

```

We can see that, strictly speaking in terms of p-value, all groups are significantly different except the control vs. sulfate treatment and control vs. nitrate treatment for LRM1b.

However, given that our growth rate estimates are HIGHLY INACCURATE, significant differences with p-values in the order of 10\^-2 or 10\^-3 should be regarded skeptically, especially if coupled with relatively small effect sizes.

# Figures

## Preliminary figures

We want to plot the two strains separately, so we will need separate datasets for each strain. We start by preparing these two datasets. We will also want to add some jitter to the x-axis to avoid overlapping of the data points.

```{r Setup plot data}

Data_plot_Pygsuia <- Data_growth_means %>% 
  filter(Culture == 'Pygsuia') %>% 
  mutate(
    Time_d_jittered = Time_d + rnorm(n(), mean = 0, sd = 0.15),  # Jitter on the x-axis
  )


Data_plot_LRM1b <- Data_growth_means %>% 
  filter(Culture == 'LRM1b') %>% 
  mutate(
    Time_d_jittered = Time_d + rnorm(n(), mean = 0, sd = 0.15),  # Jitter on the x-axis
  )

```

Now, we make the plots. We add some jitter to x-axis to avoid overlap between the points

```{r Plot Pygsuia}

ggplot(data = Data_plot_Pygsuia,
       aes(x = Time_d_jittered, y = Cell_density_mean, col = Treatment)) +
  geom_point() +  # Plot jittered points
  geom_line(aes(group = interaction(Biorep, Treatment), color = Treatment)) +
  geom_errorbar(aes(ymin = Cell_density_mean - Cell_density_SEM, 
                    ymax = Cell_density_mean + Cell_density_SEM), 
                width = 0.01) + # Add error bars
  scale_x_continuous(breaks = c(0, 3, 5, 7, 10), #set x-axis breaks
                     labels = c(0, 3, 5, 7, 10)) +
  coord_cartesian(ylim = c(0, 30000)) + #adjust zoom
  scale_y_continuous(breaks = c(0,10000, 20000, 30000),
                     labels = c(0,
                                expression(10^4),
                                expression("2·"~10^4),
                                expression("3·"~10^4)
                                )) +
  labs(x = "Time (d)",
       y = expression("Cell Density (cells·"~mL^-1~")")) +
  theme_pubr()+
  scale_colour_viridis_d(option = "magma",
                         begin = 0, end =  0.7) +
  theme(axis.title.y = element_text(margin = margin(r = 10)),
        axis.title.x = element_text(margin = margin(t = 10)),
        legend.position = "right"
        )
  

```

```{r Plot LRM1b}

ggplot(data = Data_plot_LRM1b,
       aes(x = Time_d_jittered, y = Cell_density_mean, col = Treatment)) +
  geom_point() +  # Plot jittered points
  geom_line(aes(group = interaction(Biorep, Treatment), color = Treatment)) +
  geom_errorbar(aes(ymin = Cell_density_mean - Cell_density_SEM, 
                    ymax = Cell_density_mean + Cell_density_SEM), 
                width = 0.01) + # Add error bars
  scale_x_continuous(breaks = c(0, 3, 5, 7, 10), #set x-axis breaks
                     labels = c(0, 3, 5, 7, 10)) +
  coord_cartesian(ylim = c(0, 20000)) + #adjust zoom
  scale_y_continuous(breaks = c(0,10000, 20000),
                     labels = c(0,
                                expression(10^4),
                                expression("2·"~10^4)
                                )) +
  labs(x = "Time (d)",
       y = expression("Cell Density (cells·"~mL^-1~")")) +
  theme_pubr()+
  scale_colour_viridis_d(option = "magma",
                         begin = 0, end =  0.7) +
  theme(axis.title.y = element_text(margin = margin(r = 10)),
        axis.title.x = element_text(margin = margin(t = 10)),
        legend.position = "right"
        )
  

```
```
