## Supplementary DataFile S7 for "Anaerobic protist survival in microcosms is dependent on microbiome metabolic function"

**Supplementary Data File S7**

Genome-resolved metagenomics reveal the potential for metabolic cross-feeding between anaerobic protists and bacteria

16S phylogenies of breviate associated bacteria. For Arcobacteracea and Desulfovibrioacaea, analyses of uncultured+cultured representatives and only cultured representatives were conducted. Breviate-associated sequences are coloured. Uncultured sequences from public databases are shown in grey.

Uncollapsed 18S phylogeny displayed in Figure 1A.

See https://doi.org/10.17044/scilifelab.28254575 for raw tree files and commands.

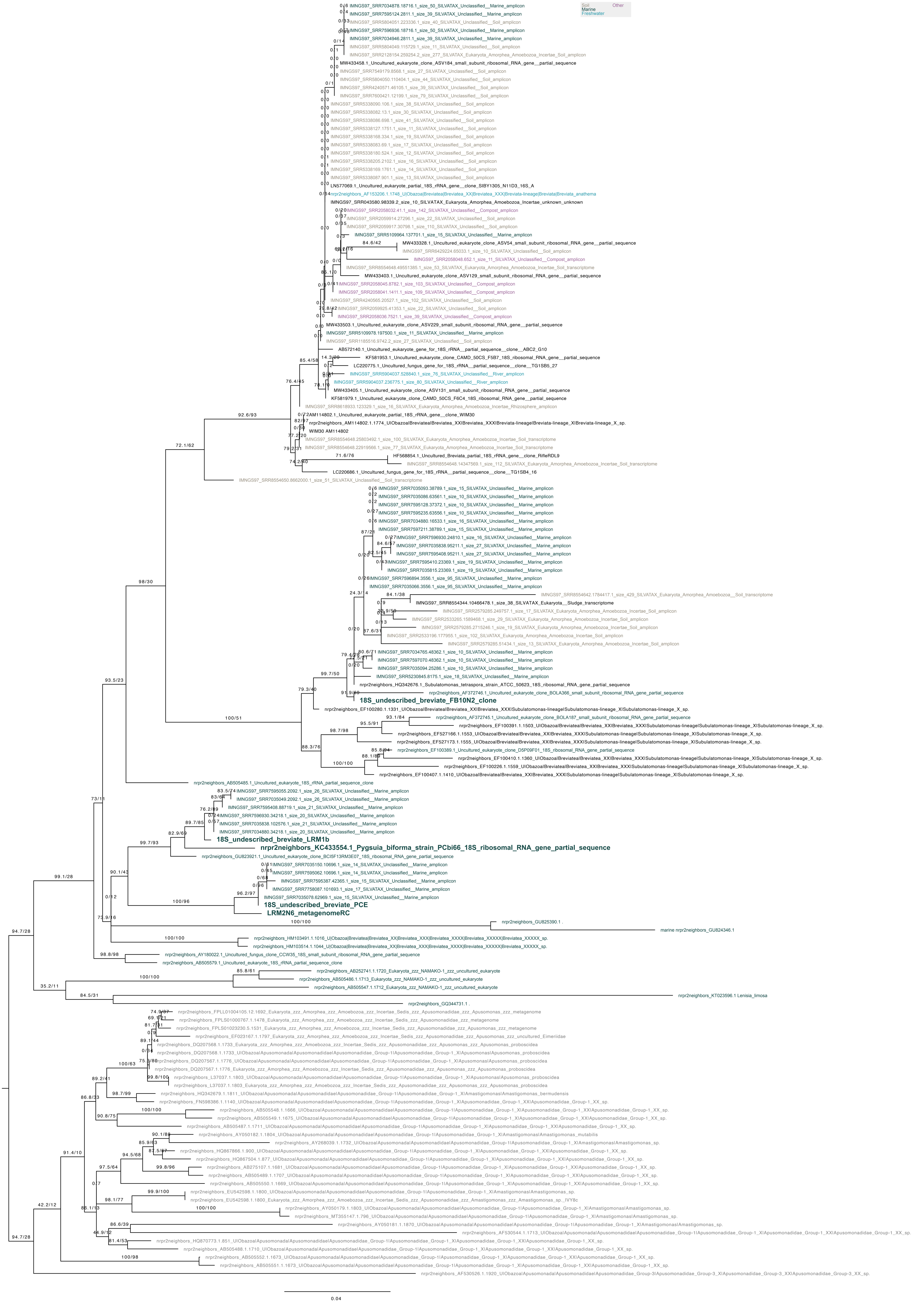
Soil
Other
Marine
Freshwater
IMNGS97_SRR7034878.18716.1_size_50_SILVATAX_Unclassified__Marine_amplicon
IMNGS97_SRR7595124.2811.1_size_39_SILVATAX_Unclassified__Marine_amplicon
IMNGS97_SRR5804051.223336.1_size_40_SILVATAX_Unclassified__Soil_amplicon
IMNGS97_SRR7596936.18716.1_size_50_SILVATAX_Unclassified__Marine_amplicon
IMNGS97_SRR7034946.2811.1_size_39_SILVATAX_Unclassified__Marine_amplicon
IMNGS97_SRR5804049.115729.1_size_11_SILVATAX_Unclassified__Soil_amplicon
IMNGS97_SRR2128154.259254.2_size_277_SILVATAX_Eukaryota_Amorphea_Amoebozoa_Incertae_Soil_amplicon
MW433458.1_Uncultured_eukaryote_clone_ASV184_small_subunit_ribosomal_RNA_gene__partial_sequence
IMNGS97_SRR7549179.8568.1_size_27_SILVATAX_Unclassified__Soil_amplicon
IMNGS97_SRR5804050.110404.1_size_44_SILVATAX_Unclassified__Soil_amplicon
IMNGS97_SRR4240571.46105.1_size_39_SILVATAX_Unclassified__Soil_amplicon
IMNGS97_SRR7600421.12199.1_size_79_SILVATAX_Unclassified__Soil_amplicon
IMNGS97_SRR5338090.106.1_size_38_SILVATAX_Unclassified__Soil_amplicon
IMNGS97_SRR5338082.13.1_size_30_SILVATAX_Unclassified__Soil_amplicon
IMNGS97_SRR5338086.698.1_size_41_SILVATAX_Unclassified__Soil_amplicon
IMNGS97_SRR5338127.1751.1_size_11_SILVATAX_Unclassified__Soil_amplicon
IMNGS97_SRR5338168.334.1_size_19_SILVATAX_Unclassified__Soil_amplicon
IMNGS97_SRR5338083.69.1_size_17_SILVATAX_Unclassified__Soil_amplicon
IMNGS97_SRR5338180.524.1_size_12_SILVATAX_Unclassified__Soil_amplicon
IMNGS97_SRR5338205.2102.1_size_16_SILVATAX_Unclassified__Soil_amplicon
IMNGS97_SRR5338169.1761.1_size_14_SILVATAX_Unclassified__Soil_amplicon
IMNGS97_SRR5338087.901.1_size_13_SILVATAX_Unclassified__Soil_amplicon
LN577069.1_Uncultured_eukaryote_partial_18S_rRNA_gene__clone_SIBY1305_N11D3_16S_A
nrpr2neighbors_AF153206.1.1748_U|Obazoa|Breviatea|Breviatea_XX|Breviatea_XXX|Breviata-lineage|Breviata|Breviata_anathema
IMNGS97_SRR043580.98339.2_size_10_SILVATAX_Eukaryota_Amorphea_Amoebozoa_Incertae_unknown_unknown
IMNGS97_SRR2058032.41.1_size_142_SILVATAX_Unclassified__Compost_amplicon
IMNGS97_SRR2059914.27296.1_size_22_SILVATAX_Unclassified__Soil_amplicon
IMNGS97_SRR2059917.30798.1_size_110_SILVATAX_Unclassified__Soil_amplicon
IMNGS97_SRR5109964.137701.1_size_15_SILVATAX_Unclassified__Marine_amplicon
MW433328.1_Uncultured_eukaryote_clone_ASV54_small_subunit_ribosomal_RNA_gene__partial_sequence
IMNGS97_SRR6429224.65033.1_size_10_SILVATAX_Unclassified__Soil_amplicon
IMNGS97_SRR2058048.652.1_size_11_SILVATAX_Unclassified__Compost_amplicon
IMNGS97_SRR8554648.49551385.1_size_53_SILVATAX_Eukaryota_Amorphea_Amoebozoa_Incertae_Soil_transcriptome
MW433403.1_Uncultured_eukaryote_clone_ASV129_small_subunit_ribosomal_RNA_gene__partial_sequence
IMNGS97_SRR2058045.8782.1_size_103_SILVATAX_Unclassified__Compost_amplicon
IMNGS97_SRR2058041.1411.1_size_109_SILVATAX_Unclassified__Compost_amplicon
IMNGS97_SRR4240565.20527.1_size_102_SILVATAX_Unclassified__Soil_amplicon
IMNGS97_SRR2059925.41353.1_size_22_SILVATAX_Unclassified__Soil_amplicon
IMNGS97_SRR2058036.7521.1_size_39_SILVATAX_Unclassified__Compost_amplicon
MW433503.1_Uncultured_eukaryote_clone_ASV229_small_subunit_ribosomal_RNA_gene__partial_sequence
IMNGS97_SRR5109978.197500.1_size_11_SILVATAX_Unclassified__Marine_amplicon
IMNGS97_SRR1185516.9742.2_size_27_SILVATAX_Unclassified__Soil_amplicon
AB572140.1_Uncultured_eukaryote_gene_for_18S_rRNA__partial_sequence__clone__ABC2_G10
KF581953.1_Uncultured_eukaryote_clone_CAMD_50CS_F5B7_18S_ribosomal_RNA_gene__partial_sequence
LC220775.1_Uncultured_fungus_gene_for_18S_rRNA__partial_sequence__clone__TG1SB5_27
IMNGS97_SRR5904037.528840.1_size_76_SILVATAX_Unclassified__River_amplicon
IMNGS97_SRR5904037.236775.1_size_80_SILVATAX_Unclassified__River_amplicon
MW433405.1_Uncultured_eukaryote_clone_ASV131_small_subunit_ribosomal_RNA_gene__partial_sequence
KF581979.1_Uncultured_eukaryote_clone_CAMD_50CS_F6C4_18S_ribosomal_RNA_gene__partial_sequence
IMNGS97_SRR8618933.123329.1_size_16_SILVATAX_Eukaryota_Amorphea_Amoebozoa_Incertae_Rhizosphere_amplicon
AM114802.1_Uncultured_eukaryote_partial_18S_rRNA_gene__clone_WIM30
nrpr2neighbors_AM114802.1.1774_U|Obazoa|Breviatea|Breviatea_XX|Breviatea_XXX|Breviata-lineage|Breviata-lineage_X|Breviata-lineage_X_sp.
WIM30 AM114802
IMNGS97_SRR8554648.25803492.1_size_100_SILVATAX_Eukaryota_Amorphea_Amoebozoa_Incertae_Soil_transcriptome
IMNGS97_SRR8554648.22919566.1_size_77_SILVATAX_Eukaryota_Amorphea_Amoebozoa_Incertae_Soil_transcriptome
HF568854.1_Uncultured_Breviata_partial_18S_rRNA_gene__clone_RifleRDL9
IMNGS97_SRR8554648.14347569.1_size_112_SILVATAX_Eukaryota_Amorphea_Amoebozoa_Incertae_Soil_transcriptome
LC220686.1_Uncultured_fungus_gene_for_18S_rRNA__partial_sequence__clone__TG1SB4_16
IMNGS97_SRR8554650.8662000.1_size_51_SILVATAX_Unclassified__Soil_transcriptome
IMNGS97_SRR7035093.38789.1_size_15_SILVATAX_Unclassified__Marine_amplicon
IMNGS97_SRR7035086.63561.1_size_10_SILVATAX_Unclassified__Marine_amplicon
IMNGS97_SRR7595128.37372.1_size_10_SILVATAX_Unclassified__Marine_amplicon
IMNGS97_SRR7595235.63556.1_size_10_SILVATAX_Unclassified__Marine_amplicon
IMNGS97_SRR7034880.16533.1_size_16_SILVATAX_Unclassified__Marine_amplicon
IMNGS97_SRR7597211.38789.1_size_15_SILVATAX_Unclassified__Marine_amplicon
IMNGS97_SRR7596930.24810.1_size_16_SILVATAX_Unclassified__Marine_amplicon
IMNGS97_SRR7035838.95211.1_size_27_SILVATAX_Unclassified__Marine_amplicon
IMNGS97_SRR7595408.95211.1_size_27_SILVATAX_Unclassified__Marine_amplicon
IMNGS97_SRR7595410.23369.1_size_19_SILVATAX_Unclassified__Marine_amplicon
IMNGS97_SRR7035815.23369.1_size_19_SILVATAX_Unclassified__Marine_amplicon
IMNGS97_SRR7596894.3556.1_size_95_SILVATAX_Unclassified__Marine_amplicon
IMNGS97_SRR7035066.3556.1_size_95_SILVATAX_Unclassified__Marine_amplicon
IMNGS97_SRR8554642.1784417.1_size_429_SILVATAX_Eukaryota_Amorphea_Amoebozoa__Soil_transcriptome
IMNGS97_SRR8554344.10466478.1_size_38_SILVATAX_Eukaryota__Sludge_transcriptome
IMNGS97_SRR2579285.249757.1_size_17_SILVATAX_Eukaryota_Amorphea_Amoebozoa_Incertae_Soil_amplicon
IMNGS97_SRR2533265.1589468.1_size_29_SILVATAX_Eukaryota_Amorphea_Amoebozoa_Incertae_Soil_amplicon
IMNGS97_SRR2579285.2715246.1_size_19_SILVATAX_Eukaryota_Amorphea_Amoebozoa_Incertae_Soil_amplicon
IMNGS97_SRR2533196.177955.1_size_102_SILVATAX_Eukaryota_Amorphea_Amoebozoa_Incertae_Soil_amplicon
IMNGS97_SRR2579285.51434.1_size_13_SILVATAX_Eukaryota_Amorphea_Amoebozoa_Incertae_Soil_amplicon
IMNGS97_SRR7034765.48362.1_size_10_SILVATAX_Unclassified__Marine_amplicon
IMNGS97_SRR7597070.48362.1_size_10_SILVATAX_Unclassified__Marine_amplicon
IMNGS97_SRR7035094.25286.1_size_10_SILVATAX_Unclassified__Marine_amplicon
IMNGS97_SRR5230845.8175.1_size_18_SILVATAX_Unclassified__Marine_amplicon
nrpr2neighbors_HQ342676.1_Subulatomonas_tetraspora_strain_ATCC_50623_18S_ribosomal_RNA_gene_partial_sequence
nrpr2neighbors_AF372746.1_Uncultured_eukaryote_clone_BOLA366_small_subunit_ribosomal_RNA_gene_partial_sequence
18S_undescribed_breviate_FB10N2_clone
nrpr2neighbors_EF100280.1.1331_U|Obazoa|Breviatea|Breviatea_XX|Breviatea_XXX|Subulatomonas-lineage|Subulatomonas-lineage_X|Subulatomonas-lineage_X_sp.
nrpr2neighbors_AF372745.1_Uncultured_eukaryote_clone_BOLA187_small_subunit_ribosomal_RNA_gene_partial_sequence
nrpr2neighbors_EF100391.1.1503_U|Obazoa|Breviatea|Breviatea_XX|Breviatea_XXX|Subulatomonas-lineage|Subulatomonas-lineage_X|Subulatomonas-lineage_X_sp.
nrpr2neighbors_EF527166.1.1553_U|Obazoa|Breviatea|Breviatea_XX|Breviatea_XXX|Subulatomonas-lineage|Subulatomonas-lineage_X|Subulatomonas-lineage_X_sp.
nrpr2neighbors_EF527173.1.1555_U|Obazoa|Breviatea|Breviatea_XX|Breviatea_XXX|Subulatomonas-lineage|Subulatomonas-lineage_X|Subulatomonas-lineage_X_sp.
nrpr2neighbors_EF100389.1_Uncultured_eukaryote_clone_D5P09F01_18S_ribosomal_RNA_gene_partial_sequence
nrpr2neighbors_EF100410.1.1360_U|Obazoa|Breviatea|Breviatea_XX|Breviatea_XXX|Subulatomonas-lineage|Subulatomonas-lineage_X|Subulatomonas-lineage_X_sp.
nrpr2neighbors_EF100226.1.1559_U|Obazoa|Breviatea|Breviatea_XX|Breviatea_XXX|Subulatomonas-lineage|Subulatomonas-lineage_X|Subulatomonas-lineage_X_sp.
nrpr2neighbors_EF100407.1.1410_U|Obazoa|Breviatea|Breviatea_XX|Breviatea_XXX|Subulatomonas-lineage|Subulatomonas-lineage_X|Subulatomonas-lineage_X_sp.
nrpr2neighbors_AB505485.1_Uncultured_eukaryote_18S_rRNA_partial_sequence_clone
IMNGS97_SRR7595055.2092.1_size_26_SILVATAX_Unclassified__Marine_amplicon
IMNGS97_SRR7035049.2092.1_size_26_SILVATAX_Unclassified__Marine_amplicon
IMNGS97_SRR7595408.88719.1_size_21_SILVATAX_Unclassified__Marine_amplicon
IMNGS97_SRR7596930.34218.1_size_20_SILVATAX_Unclassified__Marine_amplicon
IMNGS97_SRR7035838.102576.1_size_21_SILVATAX_Unclassified__Marine_amplicon
IMNGS97_SRR7034880.34218.1_size_20_SILVATAX_Unclassified__Marine_amplicon
18S_undescribed_breviate_LRM1b
nrpr2neighbors_KC433554.1_Pygsuia_biforma_strain_PCbi66_18S_ribosomal_RNA_gene_partial_sequence
nrpr2neighbors_GU823921.1_Uncultured_eukaryote_clone_BCI5F13RM3E07_18S_ribosomal_RNA_gene_partial_sequence
IMNGS97_SRR7035150.10696.1_size_14_SILVATAX_Unclassified__Marine_amplicon
IMNGS97_SRR7595062.10696.1_size_14_SILVATAX_Unclassified__Marine_amplicon
IMNGS97_SRR7595387.42365.1_size_15_SILVATAX_Unclassified__Marine_amplicon
IMNGS97_SRR7758087.101693.1_size_17_SILVATAX_Unclassified__Marine_amplicon
IMNGS97_SRR7035078.62969.1_size_15_SILVATAX_Unclassified__Marine_amplicon
18S_undescribed_breviate_PCE
LRM2N6_metagenomeRC
nrpr2neighbors_GU825390.1 .
marine nrpr2neighbors_GU824346.1
nrpr2neighbors_HM103491.1.1016_U|Obazoa|Breviatea|Breviatea_XX|Breviatea_XXX|Breviatea_XXXX|Breviatea_XXXXX|Breviatea_XXXXX_sp.
nrpr2neighbors_HM103514.1.1044_U|Obazoa|Breviatea|Breviatea_XX|Breviatea_XXX|Breviatea_XXXX|Breviatea_XXXXX|Breviatea_XXXXX_sp.
nrpr2neighbors_AY180022.1_Uncultured_fungus_clone_CCW35_18S_small_subunit_ribosomal_RNA_gene_partial_sequence
nrpr2neighbors_AB505579.1_Uncultured_eukaryote_18S_rRNA_partial_sequence_clone
nrpr2neighbors_AB252741.1.1720_Eukaryota_zzz_NAMAKO-1_zzz_uncultured_eukaryote
nrpr2neighbors_AB505486.1.1713_Eukaryota_zzz_NAMAKO-1_zzz_uncultured_eukaryote
nrpr2neighbors_AB505547.1.1712_Eukaryota_zzz_NAMAKO-1_zzz_uncultured_eukaryote
nrpr2neighbors_KT023596.1 Lenisia_limosa
nrpr2neighbors_GQ344731.1 .
nrpr2neighbors_FPLL01004105.12.1692_Eukaryota_zzz_Amorphea_zzz_Amoebozoa_zzz_Incertae_Sedis_zzz_Apusomonadidae_zzz_Apusomonas_zzz_metagenome
nrpr2neighbors_FPLS01000767.1.1478_Eukaryota_zzz_Amorphea_zzz_Amoebozoa_zzz_Incertae_Sedis_zzz_Apusomonadidae_zzz_metagenome
nrpr2neighbors_FPLS01023230.5.1531_Eukaryota_zzz_Amorphea_zzz_Amoebozoa_zzz_Incertae_Sedis_zzz_Apusomonadidae_zzz_Apusomonas_zzz_metagenome
nrpr2neighbors_EF023167.1.1797_Eukaryota_zzz_Amorphea_zzz_Amoebozoa_zzz_Incertae_Sedis_zzz_Apusomonadidae_zzz_Apusomonas_zzz_uncultured_Eimeriidae
nrpr2neighbors_DQ207568.1.1733_Eukaryota_zzz_Amorphea_zzz_Amoebozoa_zzz_Incertae_Sedis_zzz_Apusomonadidae_zzz_Apusomonas_zzz_Apusomonas_proboscidea
nrpr2neighbors_DQ207568.1.1733_U|Obazoa|Apusomonada|Apusomonadidae|Apusomonadidae_Group-1|Apusomonadidae_Group-1_X|Apusomonas|Apusomonas_proboscidea
nrpr2neighbors_DQ207567.1.1776_U|Obazoa|Apusomonada|Apusomonadidae|Apusomonadidae_Group-1|Apusomonadidae_Group-1_X|Apusomonas|Apusomonas_proboscidea
nrpr2neighbors_DQ207567.1.1776_Eukaryota_zzz_Amorphea_zzz_Amoebozoa_zzz_Incertae_Sedis_zzz_Apusomonadidae_zzz_Apusomonas_zzz_Apusomonas_proboscidea
nrpr2neighbors_L37037.1.1803_U|Obazoa|Apusomonada|Apusomonadidae|Apusomonadidae_Group-1|Apusomonadidae_Group-1_X|Apusomonas|Apusomonas_proboscidea
nrpr2neighbors_L37037.1.1803_Eukaryota_zzz_Amorphea_zzz_Amoebozoa_zzz_Incertae_Sedis_zzz_Apusomonadidae_zzz_Apusomonas_zzz_Apusomonas_proboscidea
nrpr2neighbors_HQ342679.1.1811_U|Obazoa|Apusomonada|Apusomonadidae|Apusomonadidae_Group-1|Apusomonadidae_Group-1_X|Amastigomonas|Amastigomonas_bermudensis
nrpr2neighbors_FN598386.1.1140_U|Obazoa|Apusomonada|Apusomonadidae|Apusomonadidae_Group-1|Apusomonadidae_Group-1_X|Apusomonadidae_Group-1_XX|Apusomonadidae_Group-1_XX_sp.
nrpr2neighbors_AB505548.1.1666_U|Obazoa|Apusomonada|Apusomonadidae|Apusomonadidae_Group-1|Apusomonadidae_Group-1_X|Apusomonadidae_Group-1_XX|Apusomonadidae_Group-1_XX_sp.
nrpr2neighbors_AB505549.1.1675_U|Obazoa|Apusomonada|Apusomonadidae|Apusomonadidae_Group-1|Apusomonadidae_Group-1_X|Apusomonadidae_Group-1_XX|Apusomonadidae_Group-1_XX_sp.
nrpr2neighbors_AB505487.1.1711_U|Obazoa|Apusomonada|Apusomonadidae|Apusomonadidae_Group-1|Apusomonadidae_Group-1_X|Apusomonadidae_Group-1_XX|Apusomonadidae_Group-1_XX_sp.
nrpr2neighbors_AY050182.1.1804_U|Obazoa|Apusomonada|Apusomonadidae|Apusomonadidae_Group-1|Apusomonadidae_Group-1_X|Amastigomonas|Amastigomonas_mutabilis
nrpr2neighbors_AY268039.1.1732_U|Obazoa|Apusomonada|Apusomonadidae|Apusomonadidae_Group-1|Apusomonadidae_Group-1_X|Amastigomonas|Amastigomonas_sp.
nrpr2neighbors_HQ867866.1.900_U|Obazoa|Apusomonada|Apusomonadidae|Apusomonadidae_Group-1|Apusomonadidae_Group-1_X|Apusomonadidae_Group-1_XX|Apusomonadidae_Group-1_XX_sp.
nrpr2neighbors_HQ867504.1.877_U|Obazoa|Apusomonada|Apusomonadidae|Apusomonadidae_Group-1|Apusomonadidae_Group-1_X|Apusomonadidae_Group-1_XX|Apusomonadidae_Group-1_XX_sp.
nrpr2neighbors_AB275107.1.1681_U|Obazoa|Apusomonada|Apusomonadidae|Apusomonadidae_Group-1|Apusomonadidae_Group-1_X|Apusomonadidae_Group-1_XX|Apusomonadidae_Group-1_XX_sp.
nrpr2neighbors_AB505489.1.1707_U|Obazoa|Apusomonada|Apusomonadidae|Apusomonadidae_Group-1|Apusomonadidae_Group-1_X|Apusomonadidae_Group-1_XX|Apusomonadidae_Group-1_XX_sp.
nrpr2neighbors_AB505550.1.1669_U|Obazoa|Apusomonada|Apusomonadidae|Apusomonadidae_Group-1|Apusomonadidae_Group-1_X|Apusomonadidae_Group-1_XX|Apusomonadidae_Group-1_XX_sp.
nrpr2neighbors_EU542598.1.1800_U|Obazoa|Apusomonada|Apusomonadidae|Apusomonadidae_Group-1|Apusomonadidae_Group-1_X|Amastigomonas|Amastigomonas_sp.
nrpr2neighbors_EU542598.1.1800_Eukaryota_zzz_Amorphea_zzz_Amoebozoa_zzz_Incertae_Sedis_zzz_Amastigomonas_zzz_Amastigomonas_sp._IVY8c
nrpr2neighbors_AY050179.1.1803_U|Obazoa|Apusomonada|Apusomonadidae|Apusomonadidae_Group-1|Apusomonadidae_Group-1_X|Amastigomonas|Amastigomonas_sp.
nrpr2neighbors_MT355147.1.796_U|Obazoa|Apusomonada|Apusomonadidae|Apusomonadidae_Group-1|Apusomonadidae_Group-1_X|Amastigomonas|Amastigomonas_sp.
nrpr2neighbors_AY050181.1.1870_U|Obazoa|Apusomonada|Apusomonadidae|Apusomonadidae_Group-1|Apusomonadidae_Group-1_X|Amastigomonas|Amastigomonas_sp.
nrpr2neighbors_AF530544.1.1713_U|Obazoa|Apusomonada|Apusomonadidae|Apusomonadidae_Group-1|Apusomonadidae_Group-1_X|Apusomonadidae_Group-1_XX|Apusomonadidae_Group-1_XX_sp.
nrpr2neighbors_HQ870773.1.851_U|Obazoa|Apusomonada|Apusomonadidae|Apusomonadidae_Group-1|Apusomonadidae_Group-1_X|Apusomonadidae_Group-1_XX|Apusomonadidae_Group-1_XX_sp.
nrpr2neighbors_AB505488.1.1710_U|Obazoa|Apusomonada|Apusomonadidae|Apusomonadidae_Group-1|Apusomonadidae_Group-1_X|Apusomonadidae_Group-1_XX|Apusomonadidae_Group-1_XX_sp.
nrpr2neighbors_AB505552.1.1673_U|Obazoa|Apusomonada|Apusomonadidae|Apusomonadidae_Group-1|Apusomonadidae_Group-1_X|Apusomonadidae_Group-1_XX|Apusomonadidae_Group-1_XX_sp.
nrpr2neighbors_AB505551.1.1673_U|Obazoa|Apusomonada|Apusomonadidae|Apusomonadidae_Group-1|Apusomonadidae_Group-1_X|Apusomonadidae_Group-1_XX|Apusomonadidae_Group-1_XX_sp.
nrpr2neighbors_AF530526.1.1920_U|Obazoa|Apusomonada|Apusomonadidae|Apusomonadidae_Group-3|Apusomonadidae_Group-3_X|Apusomonadidae_Group-3_XX|Apusomonadidae_Group-3_XX_sp.
0.04

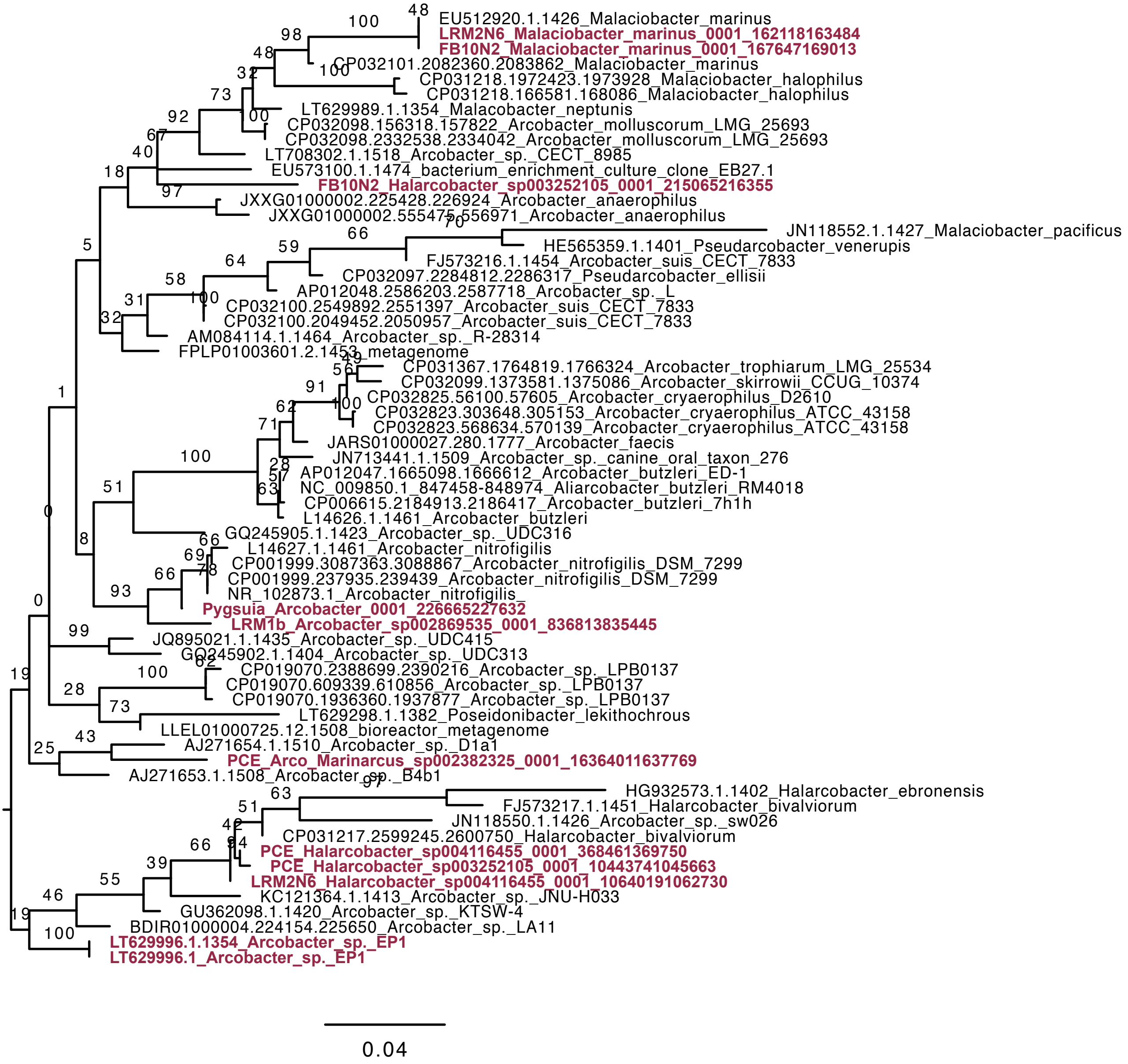
EU512920.1.1426_Malaciobacter_marinus
LRM2N6_Malaciobacter_marinus_0001_162118163484
FB10N2_Malaciobacter_marinus_0001_167647169013
CP032101.2082360.2083862_Malaciobacter_marinus
CP031218.1972423.1973928_Malaciobacter_halophilus
CP031218.166581.168086_Malaciobacter_halophilus
LT629989.1.1354_Malacobacter_neptunis
CP032098.156318.157822_Arcobacter_molluscorum_LMG_25693
CP032098.2332538.2334042_Arcobacter_molluscorum_LMG_25693
LT708302.1.1518_Arcobacter_sp._CECT_8985
EU573100.1.1474_bacterium_enrichment_culture_clone_EB27.1
FB10N2_Halarcobacter_sp003252105_0001_215065216355
JXXG01000002.225428.226924_Arcobacter_anaerophilus
JXXG01000002.555475.556971_Arcobacter_anaerophilus
JN118552.1.1427_Malaciobacter_pacificus
HE565359.1.1401_Pseudarcobacter_venerupis
FJ573216.1.1454_Arcobacter_suis_CECT_7833
CP032097.2284812.2286317_Pseudarcobacter_ellisii
AP012048.2586203.2587718_Arcobacter_sp._L
CP032100.2549892.2551397_Arcobacter_suis_CECT_7833
CP032100.2049452.2050957_Arcobacter_suis_CECT_7833
AM084114.1.1464_Arcobacter_sp._R-28314
FPLP01003601.2.1453_metagenome
CP031367.1764819.1766324_Arcobacter_trophiarum_LMG_25534
CP032099.1373581.1375086_Arcobacter_skirrowii_CCUG_10374
CP032825.56100.57605_Arcobacter_cryaerophilus_D2610
CP032823.303648.305153_Arcobacter_cryaerophilus_ATCC_43158
CP032823.568634.570139_Arcobacter_cryaerophilus_ATCC_43158
JARS01000027.280.1777_Arcobacter_faecis
JN713441.1.1509_Arcobacter_sp._canine_oral_taxon_276
AP012047.1665098.1666612_Arcobacter_butzleri_ED-1
NC_009850.1_847458-848974_Aliarcobacter_butzleri_RM4018
CP006615.2184913.2186417_Arcobacter_butzleri_7h1h
L14626.1.1461_Arcobacter_butzleri
GQ245905.1.1423_Arcobacter_sp._UDC316
L14627.1.1461_Arcobacter_nitrofigilis
CP001999.3087363.3088867_Arcobacter_nitrofigilis_DSM_7299
CP001999.237935.239439_Arcobacter_nitrofigilis_DSM_7299
NR_102873.1_Arcobacter_nitrofigilis_
Pygsuia_Arcobacter_0001_226665227632
LRM1b_Arcobacter_sp002869535_0001_836813835445
JQ895021.1.1435_Arcobacter_sp._UDC415
GQ245902.1.1404_Arcobacter_sp._UDC313
CP019070.2388699.2390216_Arcobacter_sp._LPB0137
CP019070.609339.610856_Arcobacter_sp._LPB0137
CP019070.1936360.1937877_Arcobacter_sp._LPB0137
LT629298.1.1382_Poseidonibacter_lekithochrous
LLEL01000725.12.1508_bioreactor_metagenome
AJ271654.1.1510_Arcobacter_sp._D1a1
PCE_Arco_Marinarcus_sp002382325_0001_16364011637769
AJ271653.1.1508_Arcobacter_sp._B4b1
HG932573.1.1402_Halarcobacter_ebronensis
FJ573217.1.1451_Halarcobacter_bivalviorum
JN118550.1.1426_Arcobacter_sp._sw026
CP031217.2599245.2600750_Halarcobacter_bivalviorum
PCE_Halarcobacter_sp004116455_0001_368461369750
PCE_Halarcobacter_sp003252105_0001_10443741045663
LRM2N6_Halarcobacter_sp004116455_0001_10640191062730
KC121364.1.1413_Arcobacter_sp._JNU-H033
GU362098.1.1420_Arcobacter_sp._KTSW-4
BDIR01000004.224154.225650_Arcobacter_sp._LA11
LT629996.1.1354_Arcobacter_sp._EP1
LT629996.1_Arcobacter_sp._EP1
0.04

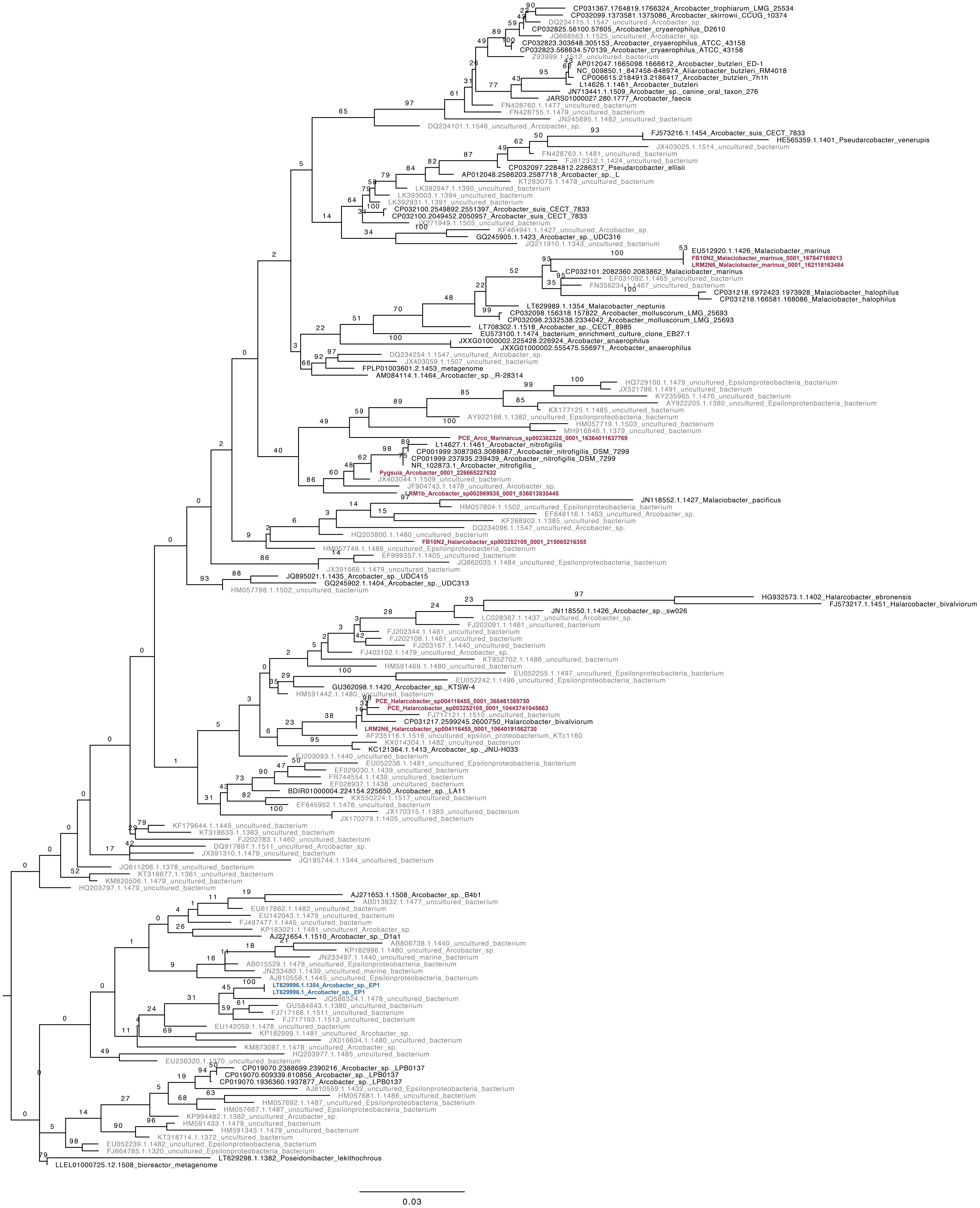
CP031367.1764819.1766324_Arcobacter_trophiarum_LMG_25534
CP032099.1373581.1375086_Arcobacter_skirrowii_CCUG_10374
DQ234115.1.1547_uncultured_Arcobacter_sp.
CP032825.56100.57605_Arcobacter_cryaerophilus_D2610
JQ668563.1.1525_uncultured_Arcobacter_sp.
CP032823.303648.305153_Arcobacter_cryaerophilus_ATCC_43158
CP032823.568634.570139_Arcobacter_cryaerophilus_ATCC_43158
Z93999.1.1512_uncultured_bacterium
AP012047.1665098.1666612_Arcobacter_butzleri_ED-1
NC_009850.1_847458-848974_Aliarcobacter_butzleri_RM4018
CP006615.2184913.2186417_Arcobacter_butzleri_7h1h
L14626.1.1461_Arcobacter_butzleri
JN713441.1.1509_Arcobacter_sp._canine_oral_taxon_276
JARS01000027.280.1777_Arcobacter_faecis
FN428760.1.1477_uncultured_bacterium
FN428755.1.1479_uncultured_bacterium
JN245695.1.1482_uncultured_bacterium
DQ234101.1.1546_uncultured_Arcobacter_sp.
FJ573216.1.1454_Arcobacter_suis_CECT_7833
HE565359.1.1401_Pseudarcobacter_venerupis
JX403025.1.1514_uncultured_bacterium
FN428763.1.1481_uncultured_bacterium
FJ612312.1.1424_uncultured_bacterium
CP032097.2284812.2286317_Pseudarcobacter_ellisii
AP012048.2586203.2587718_Arcobacter_sp._L
KT283075.1.1478_uncultured_bacterium
LK392947.1.1390_uncultured_bacterium
LK393003.1.1394_uncultured_bacterium
LK392931.1.1391_uncultured_bacterium
CP032100.2549892.2551397_Arcobacter_suis_CECT_7833
CP032100.2049452.2050957_Arcobacter_suis_CECT_7833
JX271949.1.1505_uncultured_bacterium
KF464941.1.1427_uncultured_Arcobacter_sp.
GQ245905.1.1423_Arcobacter_sp._UDC316
JQ211910.1.1343_uncultured_bacterium
EU512920.1.1426_Malaciobacter_marinus
FB10N2_Malaciobacter_marinus_0001_167647169013
LRM2N6_Malaciobacter_marinus_0001_162118163484
CP032101.2082360.2083862_Malaciobacter_marinus
EF031092.1.1465_uncultured_bacterium
FN356234.1.1467_uncultured_bacterium
CP031218.1972423.1973928_Malaciobacter_halophilus
CP031218.166581.168086_Malaciobacter_halophilus
LT629989.1.1354_Malacobacter_neptunis
CP032098.156318.157822_Arcobacter_molluscorum_LMG_25693
CP032098.2332538.2334042_Arcobacter_molluscorum_LMG_25693
LT708302.1.1518_Arcobacter_sp._CECT_8985
EU573100.1.1474_bacterium_enrichment_culture_clone_EB27.1
JXXG01000002.225428.226924_Arcobacter_anaerophilus
JXXG01000002.555475.556971_Arcobacter_anaerophilus
DQ234254.1.1547_uncultured_Arcobacter_sp.
JX403059.1.1507_uncultured_bacterium
FPLP01003601.2.1453_metagenome
AM084114.1.1464_Arcobacter_sp._R-28314
HQ729100.1.1479_uncultured_Epsilonproteobacteria_bacterium
JX521786.1.1491_uncultured_bacterium
KY235965.1.1476_uncultured_bacterium
KX177125.1.1485_uncultured_bacterium
AY922205.1.1380_uncultured_Epsilonproteobacteria_bacterium
AY922188.1.1382_uncultured_Epsilonproteobacteria_bacterium
HM057719.1.1503_uncultured_bacterium
MH916846.1.1379_uncultured_bacterium
PCE_Arco_Marinarcus_sp002382325_0001_16364011637769
L14627.1.1461_Arcobacter_nitrofigilis
CP001999.3087363.3088867_Arcobacter_nitrofigilis_DSM_7299
CP001999.237935.239439_Arcobacter_nitrofigilis_DSM_7299
NR_102873.1_Arcobacter_nitrofigilis_
Pygsuia_Arcobacter_0001_226665227632
JX403044.1.1509_uncultured_bacterium
JF904743.1.1478_uncultured_Arcobacter_sp.
LRM1b_Arcobacter_sp002869535_0001_836813835445
JN118552.1.1427_Malaciobacter_pacificus
HM057804.1.1502_uncultured_Epsilonproteobacteria_bacterium
EF648116.1.1463_uncultured_Arcobacter_sp.
KF268902.1.1385_uncultured_bacterium
DQ234096.1.1547_uncultured_Arcobacter_sp.
HQ203800.1.1480_uncultured_bacterium
FB10N2_Halarcobacter_sp003252105_0001_215065216355
HM057749.1.1488_uncultured_Epsilonproteobacteria_bacterium
EF999357.1.1405_uncultured_bacterium
JX391668.1.1479_uncultured_bacterium
JQ862035.1.1484_uncultured_Epsilonproteobacteria_bacterium
JQ895021.1.1435_Arcobacter_sp._UDC415
GQ245902.1.1404_Arcobacter_sp._UDC313
HM057798.1.1502_uncultured_bacterium
HG932573.1.1402_Halarcobacter_ebronensis
FJ573217.1.1451_Halarcobacter_bivalviorum
JN118550.1.1426_Arcobacter_sp._sw026
LC028387.1.1437_uncultured_Arcobacter_sp.
FJ202091.1.1461_uncultured_bacterium
FJ202344.1.1461_uncultured_bacterium
FJ202108.1.1461_uncultured_bacterium
FJ203167.1.1440_uncultured_bacterium
FJ403102.1.1479_uncultured_Arcobacter_sp.
KT952702.1.1486_uncultured_bacterium
HM591469.1.1480_uncultured_bacterium
EU052255.1.1497_uncultured_Epsilonproteobacteria_bacterium
EU052242.1.1496_uncultured_Epsilonproteobacteria_bacterium
GU362098.1.1420_Arcobacter_sp._KTSW-4
HM591442.1.1480_uncultured_bacterium
PCE_Halarcobacter_sp004116455_0001_368461369750
PCE_Halarcobacter_sp003252105_0001_10443741045663
FJ717121.1.1510_uncultured_bacterium
CP031217.2599245.2600750_Halarcobacter_bivalviorum
LRM2N6_Halarcobacter_sp004116455_0001_10640191062730
AF235116.1.1516_uncultured_epsilon_proteobacterium_KTc1160
KX014304.1.1482_uncultured_bacterium
KC121364.1.1413_Arcobacter_sp._JNU-H033
FJ203093.1.1440_uncultured_bacterium
EU052236.1.1481_uncultured_Epsilonproteobacteria_bacterium
EF029030.1.1439_uncultured_bacterium
FR744554.1.1439_uncultured_bacterium
EF028937.1.1438_uncultured_bacterium
BDIR01000004.224154.225650_Arcobacter_sp._LA11
KX550224.1.1517_uncultured_bacterium
EF645952.1.1476_uncultured_bacterium
JX170315.1.1383_uncultured_bacterium
JX170279.1.1405_uncultured_bacterium
KF179644.1.1445_uncultured_bacterium
KT318633.1.1383_uncultured_bacterium
FJ202783.1.1460_uncultured_bacterium
DQ917897.1.1511_uncultured_Arcobacter_sp.
JX391310.1.1479_uncultured_bacterium
JQ195744.1.1344_uncultured_bacterium
JQ611206.1.1378_uncultured_bacterium
KT318677.1.1361_uncultured_bacterium
KM820506.1.1479_uncultured_bacterium
HQ203797.1.1479_uncultured_bacterium
AJ271653.1.1508_Arcobacter_sp._B4b1
AB013832.1.1477_uncultured_bacterium
EU617862.1.1482_uncultured_bacterium
EU142043.1.1479_uncultured_bacterium
FJ497477.1.1445_uncultured_bacterium
KP183021.1.1481_uncultured_Arcobacter_sp.
AJ271654.1.1510_Arcobacter_sp._D1a1
AB806738.1.1440_uncultured_bacterium
KP182996.1.1480_uncultured_Arcobacter_sp.
JN233497.1.1440_uncultured_marine_bacterium
AB015529.1.1478_uncultured_Epsilonproteobacterium
JN233480.1.1439_uncultured_marine_bacterium
AJ810558.1.1445_uncultured_Epsilonproteobacteria_bacterium
LT629996.1.1354_Arcobacter_sp._EP1
LT629996.1_Arcobacter_sp._EP1
JQ586324.1.1478_uncultured_bacterium
GU584643.1.1380_uncultured_bacterium
FJ717168.1.1511_uncultured_bacterium
FJ717163.1.1513_uncultured_bacterium
EU142059.1.1478_uncultured_bacterium
KP182999.1.1481_uncultured_Arcobacter_sp.
JX016634.1.1480_uncultured_bacterium
KM873097.1.1478_uncultured_Arcobacter_sp.
HQ203977.1.1485_uncultured_bacterium
EU236320.1.1370_uncultured_bacterium
CP019070.2388699.2390216_Arcobacter_sp._LPB0137
CP019070.609339.610856_Arcobacter_sp._LPB0137
CP019070.1936360.1937877_Arcobacter_sp._LPB0137
AJ810559.1.1432_uncultured_Epsilonproteobacteria_bacterium
HM057681.1.1486_uncultured_bacterium
HM057692.1.1487_uncultured_Epsilonproteobacteria_bacterium
HM057667.1.1487_uncultured_Epsilonproteobacteria_bacterium
KP994482.1.1382_uncultured_Arcobacter_sp.
HM591433.1.1479_uncultured_bacterium
HM591345.1.1479_uncultured_bacterium
KT318714.1.1372_uncultured_bacterium
EU052239.1.1482_uncultured_Epsilonproteobacteria_bacterium
FJ664785.1.1320_uncultured_Epsilonproteobacteria_bacterium
LT629298.1.1382_Poseidonibacter_lekithochrous
LLEL01000725.12.1508_bioreactor_metagenome
0.03

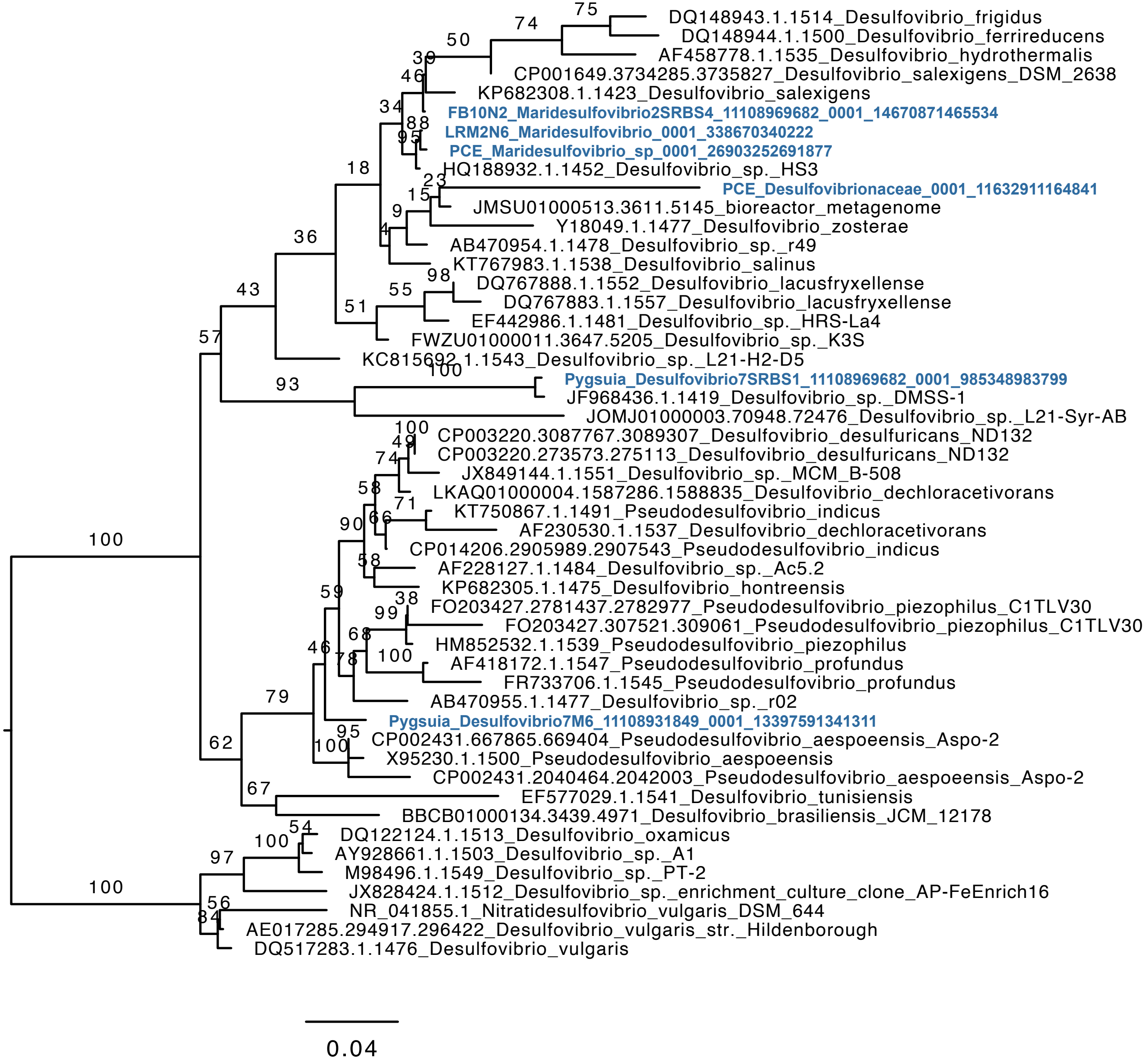

DQ148943.1.1514_Desulfovibrio_frigidus
DQ148944.1.1500_Desulfovibrio_ferrireducens
AF458778.1.1535_Desulfovibrio_hydrothermalis
CP001649.3734285.3735827_Desulfovibrio_salexigens_DSM_2638
KP682308.1.1423_Desulfovibrio_salexigens
FB10N2_Maridesulfovibrio2SRBS4_11108969682_0001_14670871465534
LRM2N6_Maridesulfovibrio_0001_338670340222
PCE_Maridesulfovibrio_sp_0001_26903252691877
HQ188932.1.1452_Desulfovibrio_sp._HS3
PCE_Desulfovibrionaceae_0001_11632911164841
JMSU01000513.3611.5145_bioreactor_metagenome
Y18049.1.1477_Desulfovibrio_zosterae
AB470954.1.1478_Desulfovibrio_sp._r49
KT767983.1.1538_Desulfovibrio_salinus
DQ767888.1.1552_Desulfovibrio_lacusfryxellense
DQ767883.1.1557_Desulfovibrio_lacusfryxellense
EF442986.1.1481_Desulfovibrio_sp._HRS-La4
FWZU01000011.3647.5205_Desulfovibrio_sp._K3S
KC815692.1.1543_Desulfovibrio_sp._L21-H2-D5
Pygsuia_Desulfovibrio7SRBS1_11108969682_0001_985348983799
JF968436.1.1419_Desulfovibrio_sp._DMSS-1
JOMJ01000003.70948.72476_Desulfovibrio_sp._L21-Syr-AB
CP003220.3087767.3089307_Desulfovibrio_desulfuricans_ND132
CP003220.273573.275113_Desulfovibrio_desulfuricans_ND132
JX849144.1.1551_Desulfovibrio_sp._MCM_B-508
LKAQ01000004.1587286.1588835_Desulfovibrio_dechloracetivorans
KT750867.1.1491_Pseudodesulfovibrio_indicus
AF230530.1.1537_Desulfovibrio_dechloracetivorans
CP014206.2905989.2907543_Pseudodesulfovibrio_indicus
AF228127.1.1484_Desulfovibrio_sp._Ac5.2
KP682305.1.1475_Desulfovibrio_hontreensis
FO203427.2781437.2782977_Pseudodesulfovibrio_piezophilus_C1TLV30
FO203427.307521.309061_Pseudodesulfovibrio_piezophilus_C1TLV30
HM852532.1.1539_Pseudodesulfovibrio_piezophilus
AF418172.1.1547_Pseudodesulfovibrio_profundus
FR733706.1.1545_Pseudodesulfovibrio_profundus
AB470955.1.1477_Desulfovibrio_sp._r02
Pygsuia_Desulfovibrio7M6_11108931849_0001_13397591341311
CP002431.667865.669404_Pseudodesulfovibrio_aespoeensis_Aspo-2
X95230.1.1500_Pseudodesulfovibrio_aespoeensis
CP002431.2040464.2042003_Pseudodesulfovibrio_aespoeensis_Aspo-2
EF577029.1.1541_Desulfovibrio_tunisiensis
BBCB01000134.3439.4971_Desulfovibrio_brasiliensis_JCM_12178
DQ122124.1.1513_Desulfovibrio_oxamicus
AY928661.1.1503_Desulfovibrio_sp._A1
M98496.1.1549_Desulfovibrio_sp._PT-2
JX828424.1.1512_Desulfovibrio_sp._enrichment_culture_clone_AP-FeEnrich16
NR_041855.1_Nitratidesulfovibrio_vulgaris_DSM_644
AE017285.294917.296422_Desulfovibrio_vulgaris_str._Hildenborough
DQ517283.1.1476_Desulfovibrio_vulgaris
0.04

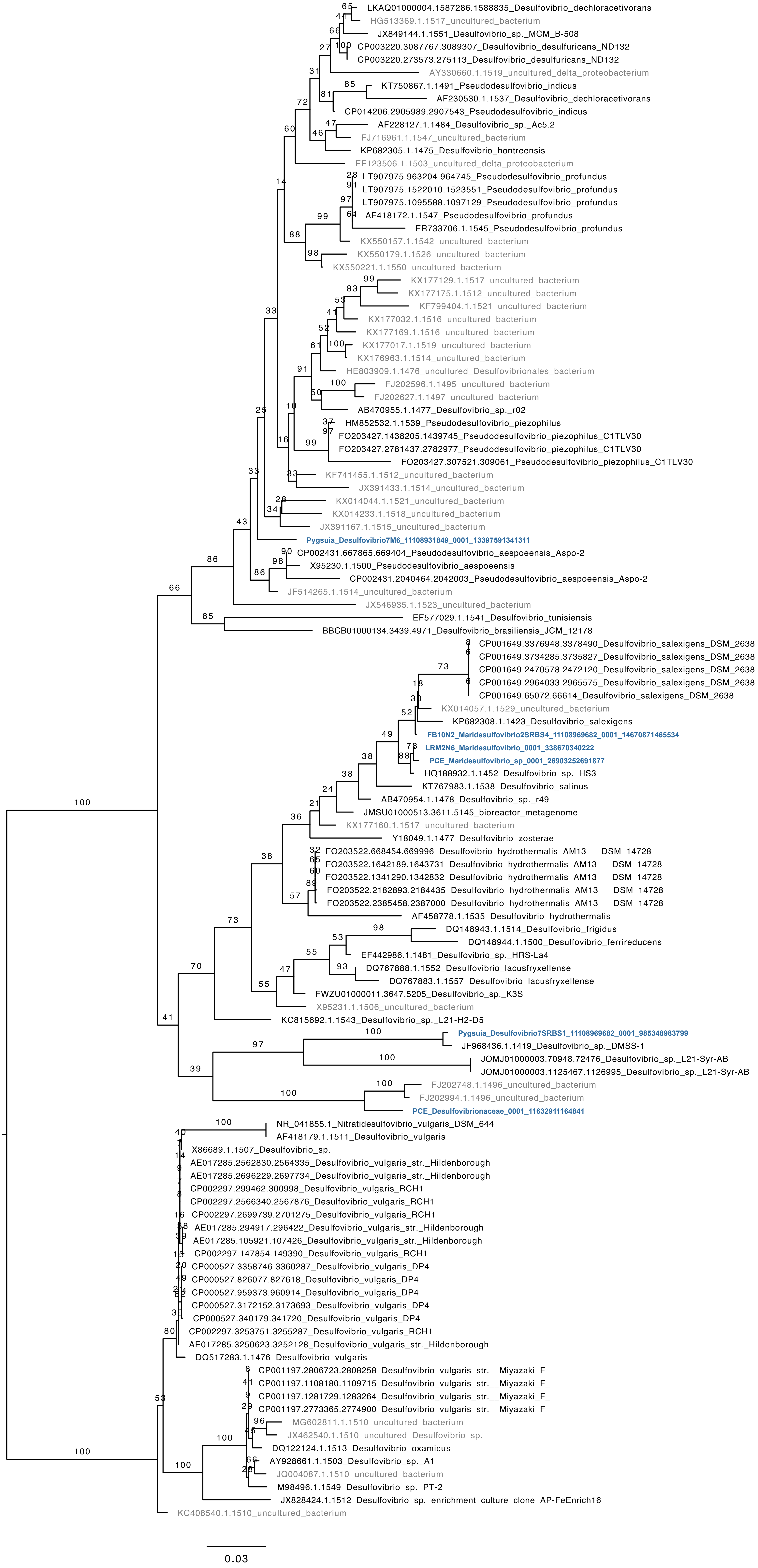

LKAQ01000004.1587286.1588835_Desulfovibrio_dechloracetivorans
HG513369.1.1517_uncultured_bacterium
JX849144.1.1551_Desulfovibrio_sp._MCM_B-508
CP003220.3087767.3089307_Desulfovibrio_desulfuricans_ND132
CP003220.273573.275113_Desulfovibrio_desulfuricans_ND132
AY330660.1.1519_uncultured_delta_proteobacterium
KT750867.1.1491_Pseudodesulfovibrio_indicus
AF230530.1.1537_Desulfovibrio_dechloracetivorans
CP014206.2905989.2907543_Pseudodesulfovibrio_indicus
AF228127.1.1484_Desulfovibrio_sp._Ac5.2
FJ716961.1.1547_uncultured_bacterium
KP682305.1.1475_Desulfovibrio_hontreensis
EF123506.1.1503_uncultured_delta_proteobacterium
LT907975.963204.964745_Pseudodesulfovibrio_profundus
LT907975.1522010.1523551_Pseudodesulfovibrio_profundus
LT907975.1095588.1097129_Pseudodesulfovibrio_profundus
AF418172.1.1547_Pseudodesulfovibrio_profundus
FR733706.1.1545_Pseudodesulfovibrio_profundus
KX550157.1.1542_uncultured_bacterium
KX550179.1.1526_uncultured_bacterium
KX550221.1.1550_uncultured_bacterium
KX177129.1.1517_uncultured_bacterium
KX177175.1.1512_uncultured_bacterium
KF799404.1.1521_uncultured_bacterium
KX177032.1.1516_uncultured_bacterium
KX177169.1.1516_uncultured_bacterium
KX177017.1.1519_uncultured_bacterium
KX176963.1.1514_uncultured_bacterium
HE803909.1.1476_uncultured_Desulfovibrionales_bacterium
FJ202596.1.1495_uncultured_bacterium
FJ202627.1.1497_uncultured_bacterium
AB470955.1.1477_Desulfovibrio_sp._r02
HM852532.1.1539_Pseudodesulfovibrio_piezophilus
FO203427.1438205.1439745_Pseudodesulfovibrio_piezophilus_C1TLV30
FO203427.2781437.2782977_Pseudodesulfovibrio_piezophilus_C1TLV30
FO203427.307521.309061_Pseudodesulfovibrio_piezophilus_C1TLV30
KF741455.1.1512_uncultured_bacterium
JX391433.1.1514_uncultured_bacterium
KX014044.1.1521_uncultured_bacterium
KX014233.1.1518_uncultured_bacterium
JX391167.1.1515_uncultured_bacterium
Pygsuia_Desulfovibrio7M6_11108931849_0001_13397591341311
CP002431.667865.669404_Pseudodesulfovibrio_aespoeensis_Aspo-2
X95230.1.1500_Pseudodesulfovibrio_aespoeensis
CP002431.2040464.2042003_Pseudodesulfovibrio_aespoeensis_Aspo-2
JF514265.1.1514_uncultured_bacterium
JX546935.1.1523_uncultured_bacterium
EF577029.1.1541_Desulfovibrio_tunisiensis
BBCB01000134.3439.4971_Desulfovibrio_brasiliensis_JCM_12178
CP001649.3376948.3378490_Desulfovibrio_salexigens_DSM_2638
CP001649.3734285.3735827_Desulfovibrio_salexigens_DSM_2638
CP001649.2470578.2472120_Desulfovibrio_salexigens_DSM_2638
CP001649.2964033.2965575_Desulfovibrio_salexigens_DSM_2638
CP001649.65072.66614_Desulfovibrio_salexigens_DSM_2638
KX014057.1.1529_uncultured_bacterium
KP682308.1.1423_Desulfovibrio_salexigens
FB10N2_Maridesulfovibrio2SRBS4_11108969682_0001_14670871465534
LRM2N6_Maridesulfovibrio_0001_338670340222
PCE_Maridesulfovibrio_sp_0001_26903252691877
HQ188932.1.1452_Desulfovibrio_sp._HS3
KT767983.1.1538_Desulfovibrio_salinus
AB470954.1.1478_Desulfovibrio_sp._r49
JMSU01000513.3611.5145_bioreactor_metagenome
KX177160.1.1517_uncultured_bacterium
Y18049.1.1477_Desulfovibrio_zosterae
FO203522.668454.669996_Desulfovibrio_hydrothermalis_AM13___DSM_14728
FO203522.1642189.1643731_Desulfovibrio_hydrothermalis_AM13___DSM_14728
FO203522.1341290.1342832_Desulfovibrio_hydrothermalis_AM13___DSM_14728
FO203522.2182893.2184435_Desulfovibrio_hydrothermalis_AM13___DSM_14728
FO203522.2385458.2387000_Desulfovibrio_hydrothermalis_AM13___DSM_14728
AF458778.1.1535_Desulfovibrio_hydrothermalis
DQ148943.1.1514_Desulfovibrio_frigidus
DQ148944.1.1500_Desulfovibrio_ferrireducens
EF442986.1.1481_Desulfovibrio_sp._HRS-La4
DQ767888.1.1552_Desulfovibrio_lacusfryxellense
DQ767883.1.1557_Desulfovibrio_lacusfryxellense
FWZU01000011.3647.5205_Desulfovibrio_sp._K3S
X95231.1.1506_uncultured_bacterium
KC815692.1.1543_Desulfovibrio_sp._L21-H2-D5
Pygsuia_Desulfovibrio7SRBS1_11108969682_0001_985348983799
JF968436.1.1419_Desulfovibrio_sp._DMSS-1
JOMJ01000003.70948.72476_Desulfovibrio_sp._L21-Syr-AB
JOMJ01000003.1125467.1126995_Desulfovibrio_sp._L21-Syr-AB
FJ202748.1.1496_uncultured_bacterium
FJ202994.1.1496_uncultured_bacterium
PCE_Desulfovibrionaceae_0001_11632911164841
NR_041855.1_Nitratidesulfovibrio_vulgaris_DSM_644
AF418179.1.1511_Desulfovibrio_vulgaris
X86689.1.1507_Desulfovibrio_sp.
AE017285.2562830.2564335_Desulfovibrio_vulgaris_str._Hildenborough
AE017285.2696229.2697734_Desulfovibrio_vulgaris_str._Hildenborough
CP002297.299462.300998_Desulfovibrio_vulgaris_RCH1
CP002297.2566340.2567876_Desulfovibrio_vulgaris_RCH1
CP002297.2699739.2701275_Desulfovibrio_vulgaris_RCH1
AE017285.294917.296422_Desulfovibrio_vulgaris_str._Hildenborough
AE017285.105921.107426_Desulfovibrio_vulgaris_str._Hildenborough
CP002297.147854.149390_Desulfovibrio_vulgaris_RCH1
CP000527.3358746.3360287_Desulfovibrio_vulgaris_DP4
CP000527.826077.827618_Desulfovibrio_vulgaris_DP4
CP000527.959373.960914_Desulfovibrio_vulgaris_DP4
CP000527.3172152.3173693_Desulfovibrio_vulgaris_DP4
CP000527.340179.341720_Desulfovibrio_vulgaris_DP4
CP002297.3253751.3255287_Desulfovibrio_vulgaris_RCH1
AE017285.3250623.3252128_Desulfovibrio_vulgaris_str._Hildenborough
DQ517283.1.1476_Desulfovibrio_vulgaris
CP001197.2806723.2808258_Desulfovibrio_vulgaris_str.__Miyazaki_F_
CP001197.1108180.1109715_Desulfovibrio_vulgaris_str.__Miyazaki_F_
CP001197.1281729.1283264_Desulfovibrio_vulgaris_str.__Miyazaki_F_
CP001197.2773365.2774900_Desulfovibrio_vulgaris_str.__Miyazaki_F_
MG602811.1.1510_uncultured_bacterium
JX462540.1.1510_uncultured_Desulfovibrio_sp.
DQ122124.1.1513_Desulfovibrio_oxamicus
AY928661.1.1503_Desulfovibrio_sp._A1
JQ004087.1.1510_uncultured_bacterium
M98496.1.1549_Desulfovibrio_sp._PT-2
JX828424.1.1512_Desulfovibrio_sp._enrichment_culture_clone_AP-FeEnrich16
KC408540.1.1510_uncultured_bacterium
0.03

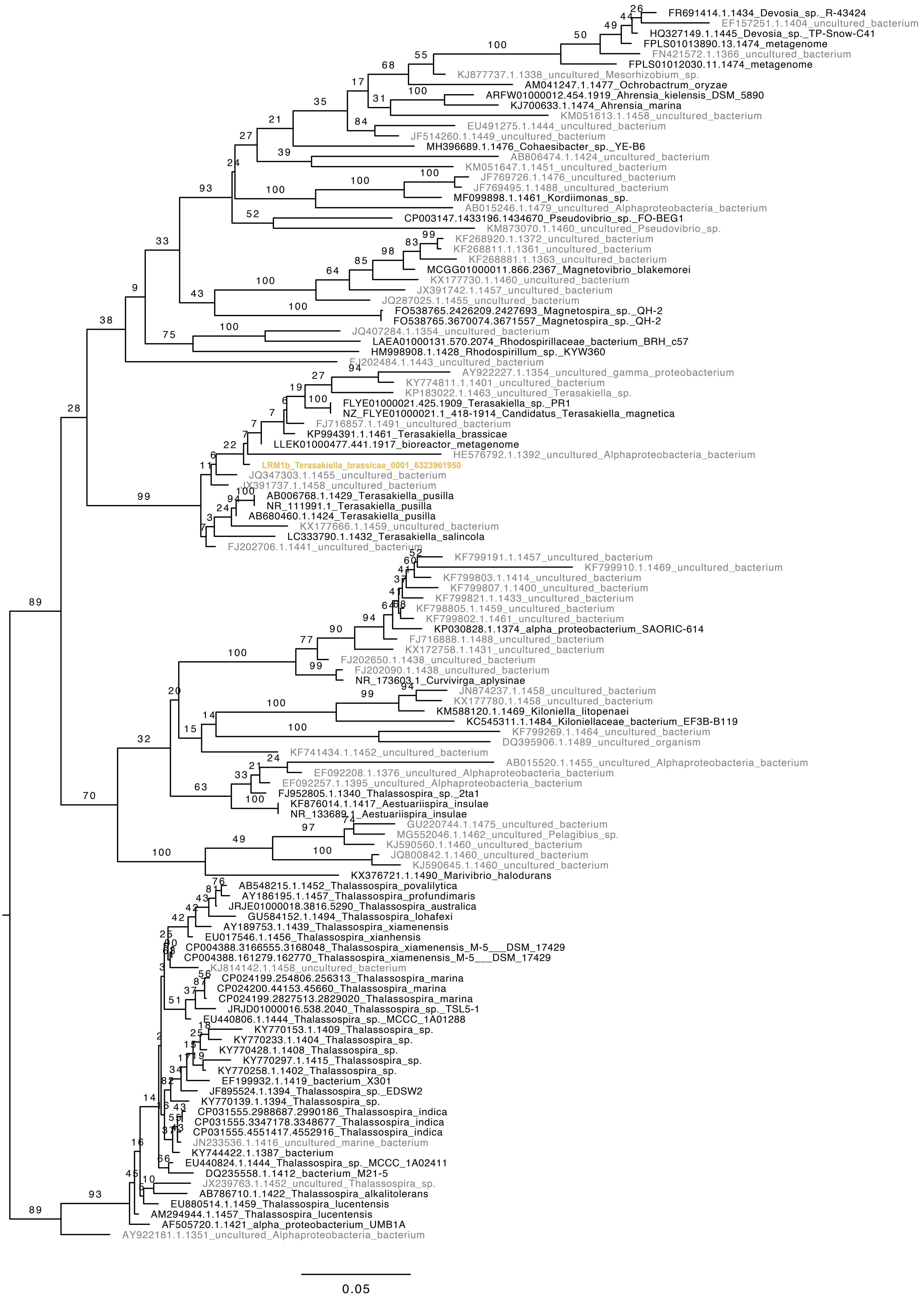
FR691414.1.1434_Devosia_sp._R-43424
EF157251.1.1404_uncultured_bacterium
HQ327149.1.1445_Devosia_sp._TP-Snow-C41
FPLS01013890.13.1474_metagenome
FN421572.1.1366_uncultured_bacterium
FPLS01012030.11.1474_metagenome
KJ877737.1.1338_uncultured_Mesorhizobium_sp.
AM041247.1.1477_Ochrobactrum_oryzae
ARFW01000012.454.1919_Ahrensia_kielensis_DSM_5890
KJ700633.1.1474_Ahrensia_marina
KM051613.1.1458_uncultured_bacterium
EU491275.1.1444_uncultured_bacterium
JF514260.1.1449_uncultured_bacterium
MH396689.1.1476_Cohaesibacter_sp._YE-B6
AB806474.1.1424_uncultured_bacterium
KM051647.1.1451_uncultured_bacterium
JF769726.1.1476_uncultured_bacterium
JF769495.1.1488_uncultured_bacterium
MF099898.1.1461_Kordiimonas_sp.
AB015246.1.1479_uncultured_Alphaproteobacteria_bacterium
CP003147.1433196.1434670_Pseudovibrio_sp._FO-BEG1
KM873070.1.1460_uncultured_Pseudovibrio_sp.
KF268920.1.1372_uncultured_bacterium
KF268811.1.1361_uncultured_bacterium
KF268881.1.1363_uncultured_bacterium
MCGG01000011.866.2367_Magnetovibrio_blakemorei
KX177730.1.1460_uncultured_bacterium
JX391742.1.1457_uncultured_bacterium
JQ287025.1.1455_uncultured_bacterium
FO538765.2426209.2427693_Magnetospira_sp._QH-2
FO538765.3670074.3671557_Magnetospira_sp._QH-2
JQ407284.1.1354_uncultured_bacterium
LAEA01000131.570.2074_Rhodospirillaceae_bacterium_BRH_c57
HM998908.1.1428_Rhodospirillum_sp._KYW360
FJ202484.1.1443_uncultured_bacterium
AY922227.1.1354_uncultured_gamma_proteobacterium
KY774811.1.1401_uncultured_bacterium
KP183022.1.1463_uncultured_Terasakiella_sp.
FLYE01000021.425.1909_Terasakiella_sp._PR1
NZ_FLYE01000021.1_418-1914_Candidatus_Terasakiella_magnetica
FJ716857.1.1491_uncultured_bacterium
KP994391.1.1461_Terasakiella_brassicae
LLEK01000477.441.1917_bioreactor_metagenome
HE576792.1.1392_uncultured_Alphaproteobacteria_bacterium
LRM1b_Terasakiella_brassicae_0001_6323961950
JQ347303.1.1455_uncultured_bacterium
JX391737.1.1458_uncultured_bacterium
AB006768.1.1429_Terasakiella_pusilla
NR_111991.1_Terasakiella_pusilla
AB680460.1.1424_Terasakiella_pusilla
KX177666.1.1459_uncultured_bacterium
LC333790.1.1432_Terasakiella_salincola
FJ202706.1.1441_uncultured_bacterium
KF799191.1.1457_uncultured_bacterium
KF799910.1.1469_uncultured_bacterium
KF799803.1.1414_uncultured_bacterium
KF799807.1.1400_uncultured_bacterium
KF799821.1.1433_uncultured_bacterium
KF798805.1.1459_uncultured_bacterium
KF799802.1.1461_uncultured_bacterium
KP030828.1.1374_alpha_proteobacterium_SAORIC-614
FJ716888.1.1488_uncultured_bacterium
KX172758.1.1431_uncultured_bacterium
FJ202650.1.1438_uncultured_bacterium
FJ202090.1.1438_uncultured_bacterium
NR_173603.1_Curvivirga_aplysinae
JN874237.1.1458_uncultured_bacterium
KX177780.1.1458_uncultured_bacterium
KM588120.1.1469_Kiloniella_litopenaei
KC545311.1.1484_Kiloniellaceae_bacterium_EF3B-B119
KF799269.1.1464_uncultured_bacterium
DQ395906.1.1489_uncultured_organism
KF741434.1.1452_uncultured_bacterium
AB015520.1.1455_uncultured_Alphaproteobacteria_bacterium
EF092208.1.1376_uncultured_Alphaproteobacteria_bacterium
EF092257.1.1395_uncultured_Alphaproteobacteria_bacterium
FJ952805.1.1340_Thalassospira_sp._2ta1
KF876014.1.1417_Aestuariispira_insulae
NR_133689.1_Aestuariispira_insulae
GU220744.1.1475_uncultured_bacterium
MG552046.1.1462_uncultured_Pelagibius_sp.
KJ590560.1.1460_uncultured_bacterium
JQ800842.1.1460_uncultured_bacterium
KJ590645.1.1460_uncultured_bacterium
KX376721.1.1490_Marivibrio_halodurans
AB548215.1.1452_Thalassospira_povalilytica
AY186195.1.1457_Thalassospira_profundimaris
JRJE01000018.3816.5290_Thalassospira_australica
GU584152.1.1494_Thalassospira_lohafexi
AY189753.1.1439_Thalassospira_xiamenensis
EU017546.1.1456_Thalassospira_xianhensis
CP004388.3166555.3168048_Thalassospira_xiamenensis_M-5___DSM_17429
CP004388.161279.162770_Thalassospira_xiamenensis_M-5___DSM_17429
KJ814142.1.1458_uncultured_bacterium
CP024199.254806.256313_Thalassospira_marina
CP024200.44153.45660_Thalassospira_marina
CP024199.2827513.2829020_Thalassospira_marina
JRJD01000016.538.2040_Thalassospira_sp._TSL5-1
EU440806.1.1444_Thalassospira_sp._MCCC_1A01288
KY770153.1.1409_Thalassospira_sp.
KY770233.1.1404_Thalassospira_sp.
KY770428.1.1408_Thalassospira_sp.
KY770297.1.1415_Thalassospira_sp.
KY770258.1.1402_Thalassospira_sp.
EF199932.1.1419_bacterium_X301
JF895524.1.1394_Thalassospira_sp._EDSW2
KY770139.1.1394_Thalassospira_sp.
CP031555.2988687.2990186_Thalassospira_indica
CP031555.3347178.3348677_Thalassospira_indica
CP031555.4551417.4552916_Thalassospira_indica
JN233536.1.1416_uncultured_marine_bacterium
KY744422.1.1387_bacterium
EU440824.1.1444_Thalassospira_sp._MCCC_1A02411
DQ235558.1.1412_bacterium_M21-5
JX239763.1.1452_uncultured_Thalassospira_sp.
AB786710.1.1422_Thalassospira_alkalitolerans
EU880514.1.1459_Thalassospira_lucentensis
AM294944.1.1457_Thalassospira_lucentensis
AF505720.1.1421_alpha_proteobacterium_UMB1A
AY922181.1.1351_uncultured_Alphaproteobacteria_bacterium
0.05
